# Accurate error control in high dimensional association testing using conditional false discovery rates

**DOI:** 10.1101/414318

**Authors:** James Liley, Chris Wallace

## Abstract

High-dimensional hypothesis testing is ubiquitous in the biomedical sciences, and informative covariates may be employed to improve power. The conditional false discovery rate (cFDR) is widely-used approach suited to the setting where the covariate is a set of p-values for the equivalent hypotheses for a second trait. Although related to the Benjamini-Hochberg procedure, it does not permit any easy control of type-1 error rate, and existing methods are over-conservative. We propose a new method for type-1 error rate control based on identifying mappings from the unit square to the unit interval defined by the estimated cFDR, and splitting observations so that each map is independent of the observations it is used to test. We also propose an adjustment to the existing cFDR estimator which further improves power. We show by simulation that the new method more than doubles potential improvement in power over unconditional analyses compared to existing methods. We demonstrate our method on transcriptome-wide association studies, and show that the method can be used in an iterative way, enabling the use of multiple covariates successively. Our methods substantially improve the power and applicability of cFDR analysis.

## 1 Introduction

In the ‘omics’ approach to biology, a large number *n* of descriptive variables are considered in the analysis of a biological system, intended to provide a near-exhaustive characterisation of the system under consideration. Typically only a small proportion of the investigated variables are associated with the behaviour of the system, and we seek to identify this subset of variables, along with the magnitude and direction of their associated effect sizes. A first step is generally to test each hypothesis in a frequentist framework, generating a corresponding set of p-values. Often, additional information is available in the form of an external covariate, which assigns a numerical value to each hypothesis which has different (unknown) distributions amongst associations and non-associations. Information from such covariates can be incorporated into hypothesis testing to improve power in detecting associations.

A range of procedures have been proposed for this type of analysis. An important consideration is the form of the (two-dimensional) rejection rule applied to the p-value-covariate pairs. An optimal procedure (in terms of minimising type 2 error and controlling type 1 error) determines rejection regions on the basis of a ratio of bivariate probability densities (PDFs) of the p-value and covariate under the null and under the alternative. One approach to the problem at hand is to estimate this ratio directly [1, 2, 3]. Other approaches include ‘filtering’ on covariate values [4], weighting hypotheses according to the value of the covariate [5, 6, 7, 8, 9], modulating a univariate test of p-values in response to the covariate in some other way [10, 11, 12], and binning covariates in order to treat each bin separately [13]. Since covariates can be of many types (continuous, categorical; univariate, multivariate; known or unknown distributional properties) and can relate to the p-values in a range of ways, this array of methods is necessary to manage the range of problem types.

The conditional false discovery rate (cFDR) circumvents the difficulties of estimating PDFs by approximating the optimal ratio using cumulative density functions (CDFs) [14]. In this case, the covariate is generally a set of p-values arising from an analogous procedure on the same variables for a second ‘conditional’ trait with an unknown degree of similarity to the trait giving rise to the primary set of p-values (which we call the ‘principal’ trait). The method has been extensively used in genomics [14, 15, 16, 17, 18, 19, 20, 21, 22, 23, 24, 25, 26, 27, 28, 29, 30, 31, 32, 33, 34, 35, 36, 37, 38]. Formally, the cFDR is a posterior probability of non-association with the principal trait given that p-values for the principal and conditional traits fall below p-value thresholds *p*, *q* respectively. It is readily estimated using empirical CDFs (ECDFs) [14].

The cFDR is a useful Bayesian quantity in its own right. Generally, the cFDR is used in effectively a frequentist way: roughly, for each observed p-value pair (*p_i_, q_i_*), we estimate the cFDR at (*p*, *q*) = (*p_i_, q_i_*) and reject the null hypothesis if this estimated value is less than some threshold *α*. This process is nearly analogous to the Benjamini-Hochberg procedure (B-H) [39] on a single set of of p-values *p_i_*, but unlike BH, it does not control the FDR at *α* (nor any other conventional measure of type-1 error rate). In a previous paper [23] we proposed a rough method to approximately control FDR in this setting, but our method was drastically conservative.

The main contribution of this paper is to propose a much improved type-1 error rate control strategy for cFDR, which improves power relative to previous methods. Our method transforms cFDR estimates into ‘v-values’, v-values’, which function analogously to p-values and can be used to control FDR or family-wise error rate (FWER). In four secondary contributions, we a) propose an improvement to the existing estimator which improves power, b) show several asymptotic results about the method and demonstrate that the effect of certain troublesome properties is small, c) enable and demonstrate iterative use of the procedure, and d) compare the general cFDR method with PDF-based, parametric and kernel density estimator (KDE)-based approaches. An R package is provided.

In this paper, we begin by describing a motivating example using transcriptome-wide association studies (TWAS). We then summarise the cFDR and its estimator, and describe its relation to the B-H procedure. We then describe our method to transform cFDR estimates into p-value-like quantities, and discuss how the cFDR approach relates to similar methods in the field. We evaluate the type-1 error rate control and power of the method, and finally describe an iterated form of the procedure for use with multiple sets of covariates.

### 1.1 Motivating example

We consider a transcript-wide association study (TWAS) [40] of breast cancer BRCA [41]) and ovarian cancer (OCA [42]), which are epidemiologically and biologically similar diseases [43]. TWAS test for association between levels of predicted expression of transcripts (gene products) in various tissues between cases (BRCA or OCA) and controls. For each transcript-tissue pair, the TWAS generates a p-value against the null hypothesis that the predicted mean expression of that transcript in that tissue is the same in case and control populations, according to a transcript-prediction rule learnt from independent data. The TWAS in question test around 10,000 gene transcripts across 45 tissues (though many transcript-tissue pairs are missing), and after we restrict to transcript-tissue pairs common to both studies, we are left with a set of ≈ 10^5^ p-values *p_BRCA_*, *p_OCA_* for association with BRCA and OCA respectively (further detail is given in supplementary material section 9.1). We wish to find which of the variables are associated with BRCA, and thus investigate a null hypothesis 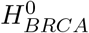 of non-association. Given established genetic correlations BRCA and OCA, we hope to leverage the OCA TWAS results to increase power in this search. We will assume that we have no prior knowledge that any variables are more likely to be BRCA- or OCA-associated, that absolute Z-scores *z_BRCA_* = Φ^−1^(*p_BRCA_*/2), *z_OCA_* = −Φ^−1^ (*p_OCA_*/2) have a block-diagonal correlation structure where block locations are known, and that under a null hypothesis 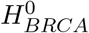 of no association with BRCA, *z_BRCA_* and *z_OCA_* are independent.

A straightforward approach is to apply the Benjamini-Hochberg (B-H) procedure to the values *p_BRCA_* (figure 1, panel A). BRCA and OCA tend to have associations at the same variables, suggesting that a rejection region should reflect this to improve power. A natural way to do this is to only consider those variables for which *z*_OCA_ exceeds some threshold, which allows rejection of 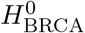 at a looser *z*_BRCA_ threshold (figure 1, panel B). FDR control is maintained under the independence assumption above [4]. However, this procedure is problematic: a threshold on *z*_OCA_ must be chosen *a priori*, and variables with *z*_OCA_ falling below the red line have 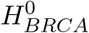 retained automatically.

**Figure 1:**
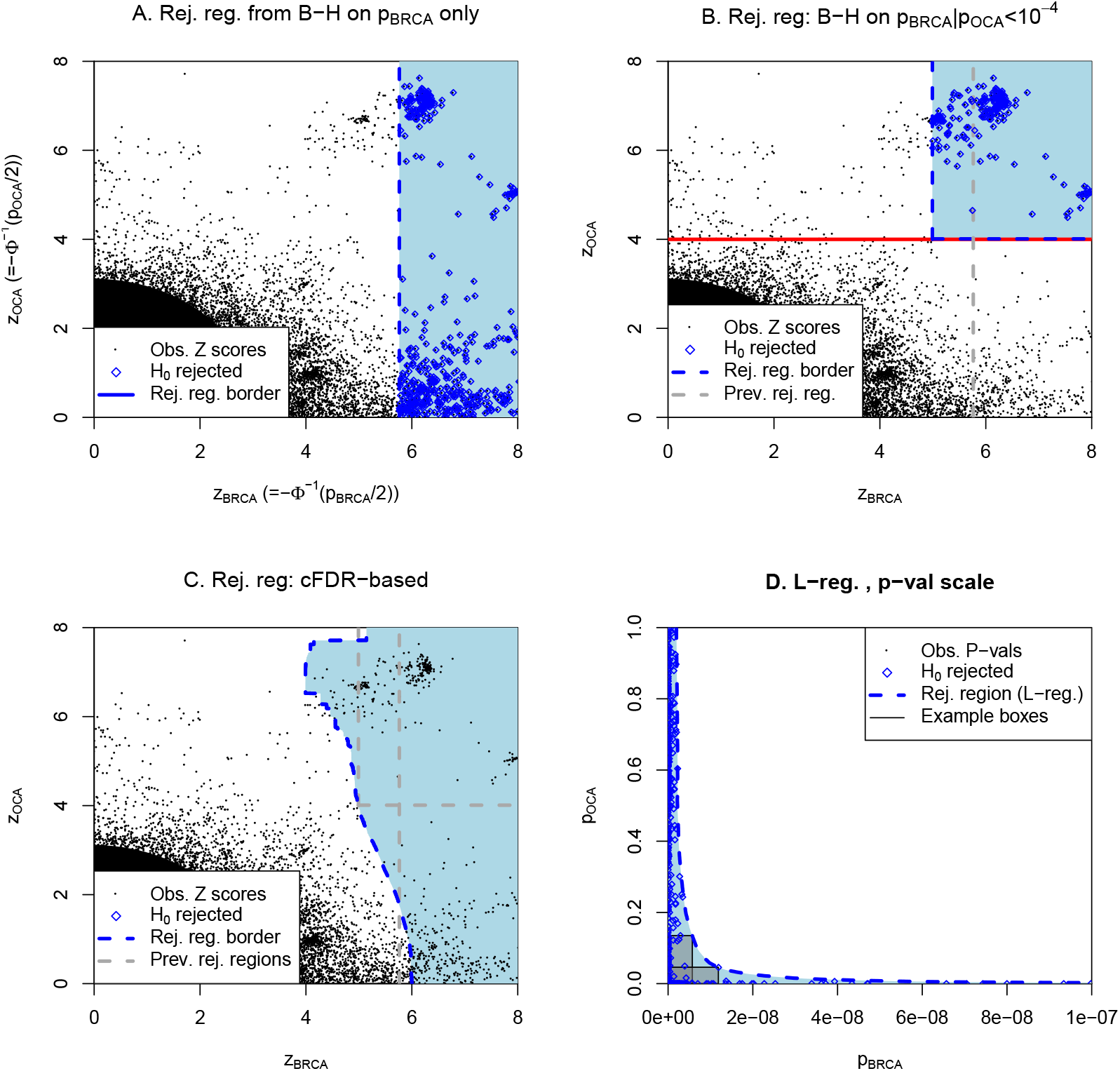
Illustration of cFDR approach using example data from TWAS study of breast cancer (BRCA), conditioning on ovarian cancer (OCA). Each plot shows test statistics from BRCA (x axis) and OCA (y axis) on either the Z (**A**-**C**) or p-value (**D**) scale. All rejection regions use methods to control FDR at < 1 × 10^−6^. **A** Benjamini-Hochberg (B-H) procedure applied to BRCA statistics alone, leads to a rejection region to the right of the blue dashed vertical line. **B** B-H applied to those variables for which *z*_OCA_ exceeds the threshold shown by a solid red line. **C** cFDR procedure: for the *i*th values *z_BRCA_*(*i*) *z_OCA_*(*i*), a B-H procedure aiming to control the FDR at *α* is conducted on only the variables for which *z_OCA_* ≥ *z_OCA_*(*i*), and if the *i*th null hypothesis is rejected during this procedure, it is rejected overall. We term the rejection region corresponding to this value *α* an ‘L-region’ *L*(*α*), shown as the shaded region. **D** The exposition that follows using p-values rather than Z scores, and so we reproduce the data and *L*(*α*) on the p-value scale. On this scale, the estimated cFDR at a point *p_BRCA_, p_OCA_* can be considered an estimate of the FDR corresponding to a fixed rejection region given by the box with *p_BRCA_, p_OCA_* as its top-right corner, and the L-region *L*(*α*) roughly as the locus of top-right corners of boxes with estimated cFDR equal to *α*. Two such boxes are illustrated on the figure.

The cFDR procedure circumvents this problem (figure 1, panel C). The associated rejection region, which we term an L-region, ‘adapts’ to the joint distribution of *z_BRCA_* and *z_OCA_*. For small *α*, the L-region approximates an optimal rejection region (appendix 8.1). A major shortcoming is that although a B-H procedure with FDR controlled at *α* is repeatedly used to generate the L-region, the overall FDR is not controlled at *α*, nor any straightforward function of *α* ([23]; appendix 8.2).

In this paper, we demonstrate a straightforward and effective way to control the type-1 error rate (specifically FDR or FWER) in the cFDR procedure. As *α* varies from 0 to 1, the leftmost boundary of the L-region ‘sweeps’ across the entire (+,+) quadrant, and for *α*_1_ < *α*_2_, we have *L*(*α*_1_) ⊆ *L*(*α*_2_). Thus we can associate each point (x, y) in the (+,+) quadrant with the smallest L-region containing it, which will generally have (x, y) on its leftmost border. Loosely, we control FDR by estimating the probability that each point would lie within its associated region under 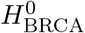. We term this the v-value, which has similar properties to a p-value and can be used in the B-H procedure.

Care must be taken when applying rejection rules to the same data on which those rules were determined, so we use a leave-one-out procedure which avoids this problem (section 3, appendix 8.4). We show that the rejection region generated by the cFDR approximates the best-possible rejection region (section 2.2, appendix 8.1), and that rejection regions converge reasonably fast as the number of variables under consideration increases (section 8.3). The rejection region is non-parametric, and we show that the cFDR method can outperform parametric methods (section 5.2). Finally, the v-values may be considered ‘adjusted’ p-values, which enables straightforward iteration of the method with further sets of p-values at the same variables, discussed in section 5.5.

## 2 Review of cFDR estimator

### 2.1 Definitions

Assume that we have results from *n* pairs of hypothesis tests against two series of null hypotheses 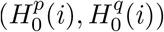 in the form of a set *S* of bivariate p-values *S* = (*p_i_, q_i_*), *i* = 1..*n*. In our motivating example, 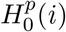 and 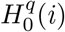 are non-association of the *i*th tissue-gene pair with BRCA and OCA respectively. We consider 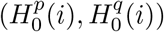 to be realisations of independent Bernoulli random variables 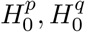 satisfying 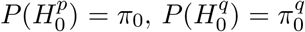, and *p_i_*, *q_i_* to be IID realisations of random variables *P*, *Q* satisfying:

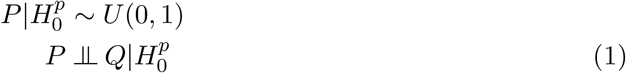

although assumption (1) can be relaxed. We denote

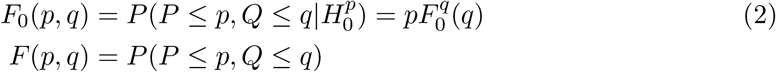

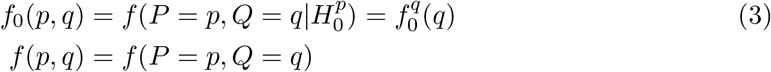

where the separability of (2) and (3) is due to assumption (1).

### 2.2 Optimal procedure

Under 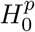, the probability of a random instance of (*P*, *Q*) falling in a region *R* is *∫_R_ f*_0_(*p*, *q*)*dpdq*. To find an ideal two-dimensional rejection region for hypothesis testing, we wish to fix this value at a level *α* while maximising the probability *∫_R_ f*_0_(*p*, *q*)*dpdq*. This optimal region (or one such optimal region) is given by the set of points {(*p*, *q*) : *f*_0_(*p*, *q*)/*f*(*p*, *q*) ≥ *k_α_*} for some *k_α_* (a formal statement and proof are given in appendix 8.1, and this is also shown in various forms in [1, 2, 3]). In equivalent terms, an optimal decision rule for the set *S* would rank p-value pairs according to *f*_0_(*p_i_, q_i_*)/*f*(*p_i_, q_i_*) or equivalently 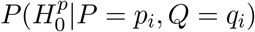.

A natural approach is to estimate *f*_0_ and *f* using a parametric approximation [1, 3] or local approximations using kernel density estimates (KDEs) [2] or spline models [44]. However, PDFs are difficult to estimate in general, and there may be little reason to believe parametric assumptions are satisfied; in our motivating example (figure 1, panels A-C) there is little reason to think that a smooth rejection region would be optimal.

### 2.3 Conditional false discovery rate

The conditional false-discovery rate [14] takes an alternative approach of instead ranking points by an estimate of *F*_0_(*p*, *q*)/*F*(*p*, *q*). This estimate is obtained by estimating the monotonically-related quantity:

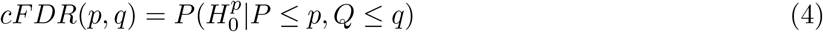

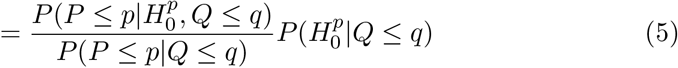

Suppose we have a multiset *X* of p-value pairs (*p_i_, q_i_*). If almost all these pairs are iid realisations (*p_i_, q_i_*) of (*P*, *Q*), then for fixed *p*, *q*, the empirical CDFs

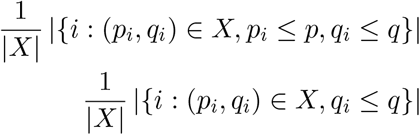

are consistent estimators of *P*(*P* ≤ *p*, *Q* ≤ *q*), *P*(*Q* ≤ *q*) respectively. Given assumption (1), we have 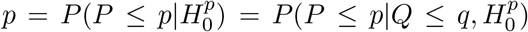 and (for the moment) we may conservatively approximate 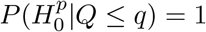. Given *X*, we thus define the estimated cFDR (denoted 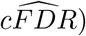), as a function of two variables (*p*, *q*) ∈ (0, 1):

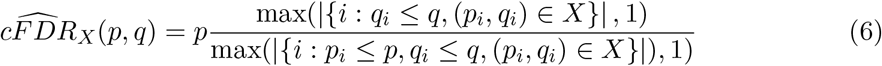

For fixed *p*, *q*, 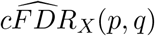 is a generally-biased but consistent estimator of 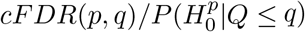, which converges uniformly on fixed regions at a rate of *O*(*n*^−1/2^) (see appendix 8.3), and it is usually a downwards-biased (conservative) estimator of *cFDR*(*p*, *q*).

Approximating 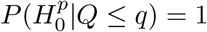 in equation (6) disregards any variation on 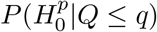 with *q*, so we introduce at this stage an estimate of 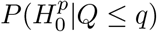, which we can multiply with 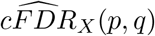 to improve the accuracy of approximation of *cFDR*(*p*, *q*). Our estimate is

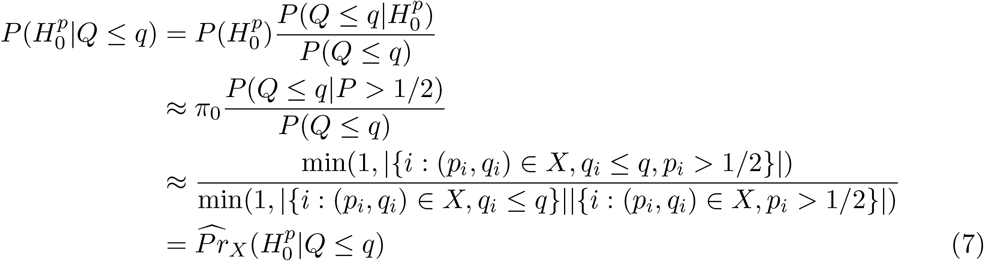

where we approximate 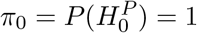. We denote

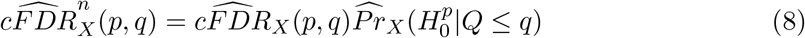

Estimating *π*_0_ (rather than setting *π*_0_ = 1) would uniformly scale all estimates of *cFDR*(*p*, *q*), which has no effect on our rejection procedure.

In the hypothesis testing setting, we aim to use 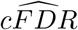 or 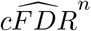 to construct a decision rule on our set *S* of observed p-value pairs (we will forego the *n* superscript from now, with the understanding that it may be added). A simple approach is to reject 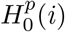 if 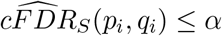, but 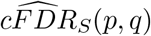 is not monotonically increasing with *p* and we do not want to reject 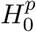 for some (*p_i_, q_i_*) and not for some other pair (*p_j_, q_j_*) with *q_i_* = *q_j_* but *p_j_* < *p_i_*. Hence, we can use the decision rule (as per [14])

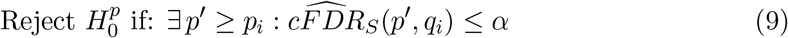

This enables a rejection region with a single rightmost boundary, as shown in panels D in figure 1. It closely parallels the B-H [39] procedure (B-H) on a set of p-values. Suppose we have a set *S*_1_ of p-values *p*_1_, *p*_2_, …, *p_n_*, and define

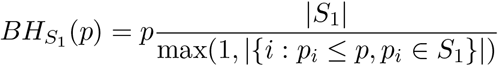

Then the B-H procedure can be written as

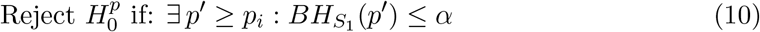

The B-H procedure controls FDR at *α*, and if it is performed with *S*_1_ as the subset of *S* for which *q_i_* ≤ *γ* (where *γ* is a threshold chosen independently of *S*), the FDR will still be controlled at *α* if assumption (1) is satisfied [4, 5]. The rejection procedure (9) is equivalent to repeatedly performing this ‘thresholded’ B-H at *γ* = *q_i_*, and using that decision rule for point (*p_i_, q_i_*) (panel C, figure 1).

When procedure (9) is used, the FDR is no longer controlled at *α*, and indeed can exceed *α* by an arbitrary proportion. This is most easily seen in the extreme case

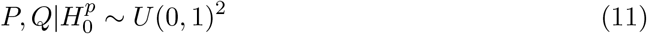

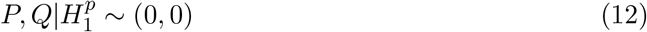

in which we can show that the FDR *α_TRUE_* of rejection procedure 9 applied to 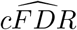 satisfies

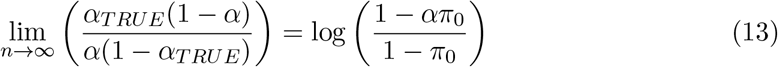

and when applied to 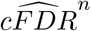 satisfies

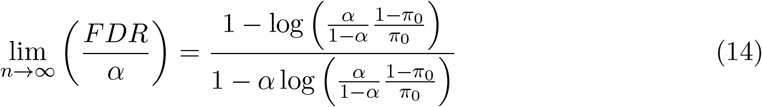

with the corollary that the actual FDR when using rejection procedure 9 can be an arbitrarily large multiple of *α*. These formulae are proved in theorem 6, appendix 8.2.

Previous work using the cFDR generally interprets it in a Bayesian context, without requiring a bound on FDR or FWER. In [23] we introduced a method to choose an *α*\* such that rejection criterion (9) roughly controlled the FDR at *α*, but that method was overly conservative, generally controlling FDR at a far lower level than needed.

## 3 Map from p-value pairs to v-values

We identify a ‘rejection region’ associated with 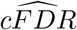 by adding a ‘test point’ (*p*, *q*) to a set of points *X* and considering the region for which a hypothesis corresponding to (*p*, *q*) is rejected under (9) with *S* = *X* + (*p*, *q*).

The function 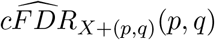 is now defined on the unit square. It is difficult to use, however: when considered as a function of *p* with fixed *q*, it does not monotonically increase with *p*. We thus define

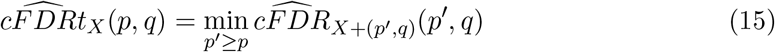

and define the ‘L-region’ *L_X_*(*α*):

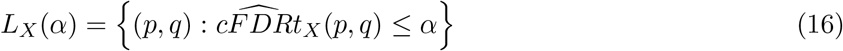

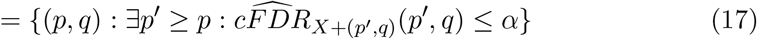

and define the ‘L-curve’ as the rightmost boundary of this region. We note that

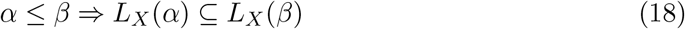

We now show the following, and include the brief proof:

### Theorem 1.

*Assume that P*, 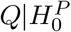 *is a bivariate continuous random variable with support* [0, 1]^2^, *and set μ*_0_ *as its induced measure. Suppose X* = (*p_i_, q_i_*) ∈ [0, 1]^2^, *i* ∈ 1..*n is a fixed finite set of points. Define L_X_*(*α*) *as per equation 16, with 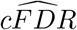 defined as per either equation 6 or equation 8, and define the ‘v-value’ as a function of p, q* ∈ (0, 1)^2^

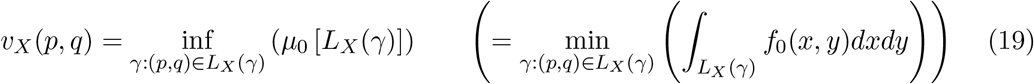

*the second definition being valid in our more restrictive context (section 2). Then for α* ∈ (0, 1)

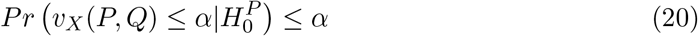

*Proof*. Since *X* is finite *L_X_*(*α*) is Lesbegue-measurable, so the integral in 19 is well-defined.

Given *α* ∈ (0, 1), let

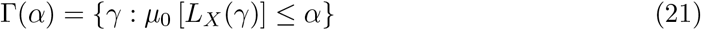

Suppose there exists *γ*(*α*) ∈ Γ(*α*) with (*p*, *q*) ∈ *L_X_*(*γ*(*α*)). Then from definition 19 we have

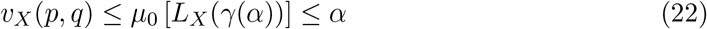

so, using property 18

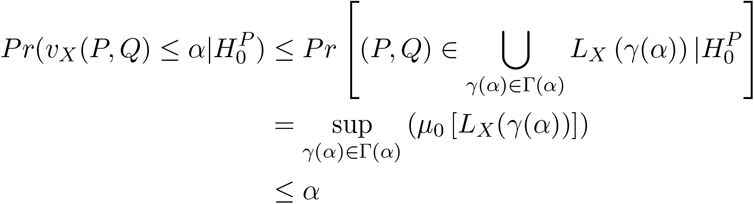

If there exists *γ* such that *α* = *μ*_0_[*L*(*γ*)] then (*p*, *q*) ∈ *L_X_*(*γ*) ⇔ *v_X_*(*p*, *q*) ≤ *α*, so equality is achieved in 20

### Remark 1.1.

*With* Γ *defined as per 21, and setting μ*_1_ *as the induced measure of P*, 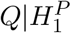, *the power of the rejection procedure*

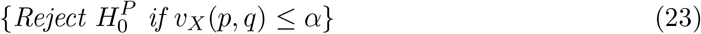

*is*

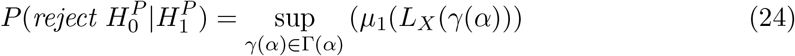

*and the type-1 error rate is* ≤ *α*

Theorem 1 assumes the probability measure *F_0_* of *P*, 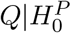 is known. In practice, it must be estimated. This can be readily done given assumption (1), as will be shown in section 3.2. We can think of the function *v_X_*(*p*, *q*) as a map from the unit square to the unit interval, where the map is defined by the points *X*.

We note that property 20 indicates that the value *v_X_*(*p*, *q*) is interpretable as a p-value against 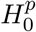, using 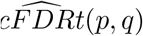 as a test statistic (it may even be thought of as a definition of a p-value [45]). In this sense, it is slightly conservative due to the ‘≤ *α*’ rather than ‘= *α*’ in 20. Resultant v-values may thus be used in the B-H procedure to control FDR, or used with a Šidák correction to control FWER. For the remainder of this paper, we will seek to control FDR by the B-H procedure.

In order to use theorem 1 to generate a p-value for a test point (*p*, *q*) we must assume *X* is ‘fixed’. In practice, this means (*p_i_, q_i_*) must be independent of values (*p*, *q*); hence, not in *X*.

Given our set of points *S* = (*p_i_, q_i_*), this is easily managed: to test (*p_i_, q_i_*), we simply leave (*p_i_, q_i_*) out of the points used to define the we use on (*p_i_, q_i_*) itself. That is, given our set *S* of datapoints as above, we define the ‘leave-one-out’ v-value

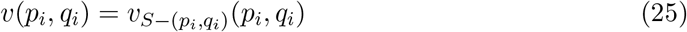

The problem can also be managed by leaving out blocks of points; for a partition of 1..*n* into blocks 1, 2, …, *k*, supposing *i* is in block *b*(*i*), the ‘block-out’ v-value is defined as

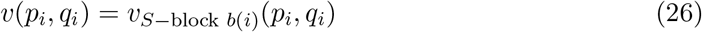

If observations (*p_i_, q_i_*) are not independent but have a block-diagonal correlation structure, then this procedure is necessary in order to ensure property (20) holds for (*p*, *q*) = (*p_i_, q_i_*): since each observation (*p_i_, q_i_*) carries information about the other p-value pairs it is correlated with, removing it will not remove the influence of point (*p_i_, q_i_*) on the map. In this case, blocks should be chosen so that p-value pairs are independent between blocks, but possibly dependent within blocks. Such structure arises often in -omics experiments; in genetics, independence of allele counts may be assumed between chromosomes, but not generally within.

For comparison, we also define the ‘naive’ v-value

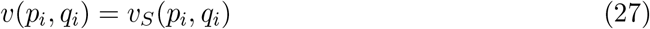

where *p_i_, q_i_* is in *S*.

In the subsequent section, we note several asymptotic properties of L-regions and v-values. We note that consistency of L-region estimation is not necessary for type-1 error rate control: there is no requirement in theorem 1 for the values in *X* to have the same distribution as *P*, *Q*. L-regions are also identical under monotonic transformations of the function 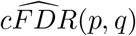, so the method is unaffected by the approximation 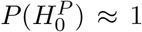 in equation 7.

### 3.1 Asymptotic properties of L-regions and v-values

We describe several asymptotic properties of L-regions and v-values. Estimator 7 is not generally consistent, so we focus our attention on properties of regions defined using 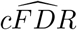 rather than 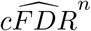. We divide out the quantity estimated in 7, write *F*(*q*) = *Pr*(*Q* ≤ *q*) and define

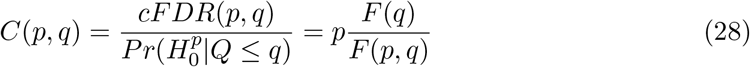

which we generally assume to be differentiable on the unit square. We recall definition 15 and note the following (proved in appendix 8.3:

#### Theorem 2.

*Let R be the region of the unit square for which F*(*p*, *q*) ≥ *γ* > 0 *and F*(*q*) > 0. *Then on R*, 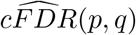 *converges uniformly to C*(*p*, *q*), *and if ∂C*(*p*, *q*)/*∂_p_* ≥ 0, *then so does* 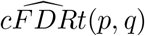

This theorem indicates that the empirical L-curve converges to a contour of *C*(*p*, *q*), where the contour exists. Although the region *R* does not cover the entire unit square, in practice it usually has Lesbegue measure 0: if 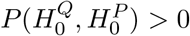 and *P*, 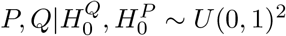 then *F*(*p*, *q*) is bounded away from 0 on (0, 1)^2^. We note that *C*(*p*, *q*) is meaningful only when *F*(*p*, *q*) > 0.

Because of the uniform convergence, this can be translated into a statement about v-values. Given an L-region *L*(*α*), we define the M-region as the ‘expected’ L-region:

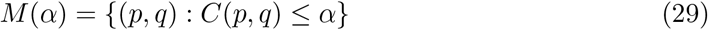

and the ‘error’ on the v-value *v* = *∫*_*L*(*α*)_ *f*_0_(*p*, *q*)*dpdq* as

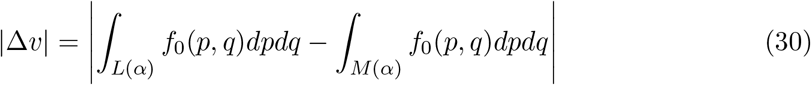

Now we have

#### Theorem 3.

*Define R as in theorem 2, and further assume that* 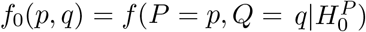 *is known and on R we have ∂C*(*p*, *q*)/*∂p ≥ γ*_2_ > 0. *Write R^c^* = [0, 1]^2^ \ *R. Then the maximum error on any v-value is*

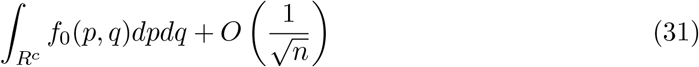

thus, as *n* → ∞, v-values based on 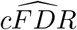 converge at a rate *O*(*n*^−1/2^) to those that would be obtained using *C*(*p*, *q*), plus a fixed error. We also note that under the conditions in both theorems, if *R* has negligible Lesbegue measure, and there exists *γ* such that

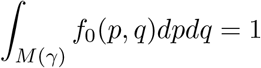

then as *n* → ∞ the power of rejection procedure 23 satisfies

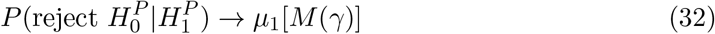

where *μ*_1_ is defined as in equation 23. Finally, we note that neither consistency nor unbiasedness of the 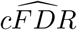 estimator is necessary for the p-value property in theorem 1 to hold. Proofs of theorems 2 and 3 are given in appendix 8.3.

### 3.2 Estimation of *P*, 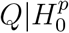

Recalling equation (3), we may write 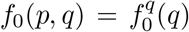. To estimate 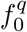, we assume that 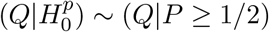, and approximate the latter with a mixture-Gaussian distribution

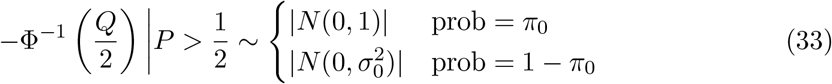

where *N*(*μ, σ*) is the normal distribution with mean *μ* and variance *σ*^2^. Estimates 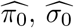 of *π*_0_ and *σ*_0_ can be readily made using an expectation-maximisation algorithm [46], using the values *q_i_* for which the corresponding *p_i_* is ≥ 1/2. We then estimate the density *f*_0_(*p*, *q*) of *P*, 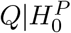 as:

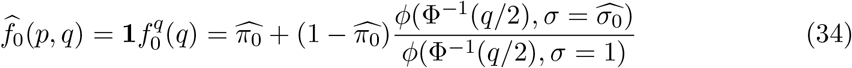

where *ϕ* is the normal density function with SD *σ*. If *P*, *Q* have a known dependence under 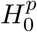, an alternative distribution can be used for computing *v*(*L*) (see supplementary material, section 9.2). The PDF 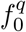 could be estimated in other ways; for example, a kernel density estimate [47].

### 3.3 Correlation between v-values

Decision rules based on multiple p-values generally require adjustment if p-values are dependent (eg [39]). If v-values are obtained by the leave-one-out procedure (25) they are slightly pairwise dependent. The dependence is small; if *X*′ = *X* − (*p_i_, q_i_*) − (*p_j_, q_j_*) the values *v_X′_* (*p_i_, q_i_*), *v_X′_ (p_j_, q_j_*) are independent, so the pairwise dependence between v-values corresponding to (*p_i_, q_i_*), (*p_j_/q_j_*) only arises from the differences *v_X′+(p_j_, q_j_)_*(*p_i_, q_i_*) − *v_X′_*(*p_i_, q_i_*), *v_X′+(p_i_, q_i_)_*(*p_j_, q_j_*) − *v_X′_*(*p_j_, q_j_*); that is, the effect of a single point ((*p_j_, q_j_*), (*p_i_, q_i_*) respectively) on the map *v_X′_* defined by |*X′*| = |*X*| − 2 points. Indeed, we show that the expected change to v-values on adding a single new point is small:

#### Theorem 4.

*Suppose we add a point* (*p*, q\**) *to a set of n points* (*p_i_, q_i_*), *considered as realisations of P, Q, and conditions are satisfied for convergence of v-values as above. Let* Δ*v*(*L*(*α*)) *be the shift in a v-value corresponding to an L-curve L*(*α*) *after adding* (*p*, q\**). *Then*

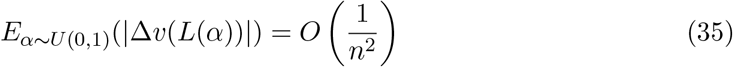

The proof is given in appendix 8.4. When v-values are defined using block-out as in (26), v-values are independent within-block but dependent between blocks. The Benjamini-Hochberg procedure is also sensitive to higher-order (non-pairwise) dependence between v-values, but we show by simulation in section 5, residual dependence does not generally lead to failure of FDR control, even when we increase dependence by enforcing correlation between observations *p_i_, p_j_* and between *q_i_, q_j_*.

### 3.4 Algorithm

We can now present our final algorithm.

**Algorithm 1.**
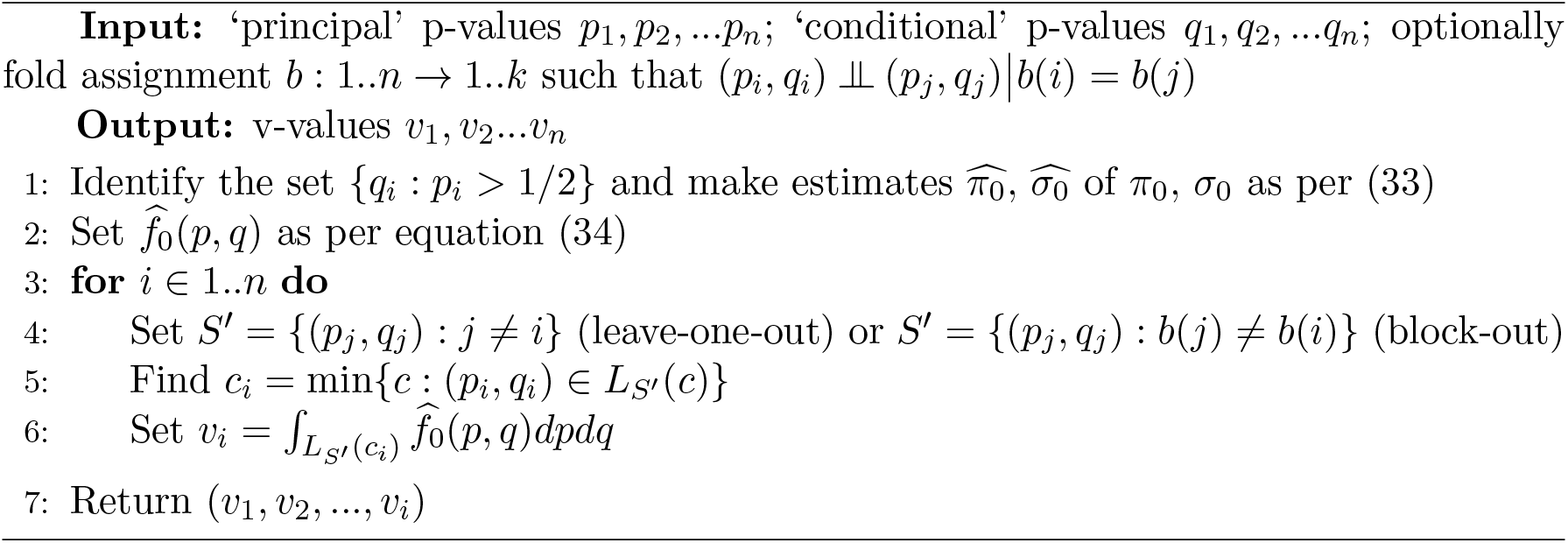
Controlling type-1 error rate in cFDR

We can interpret *v_i_* as ‘the probability that a randomly-chosen (*p*, *q*) pair has a more extreme 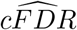 value than 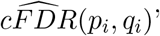; that is, as a p-value. This allows straightforward FWER or FDR control, especially as v-values are almost independent. The v-values order hypotheses such that a rejection rule {reject 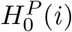 if *v_i_* ≤ *α*} has near-optimal power, in terms of corresponding to near-optimal forms for rejection regions. Much of our methodology can also be used if values *q_i_* are not p-values for some second trait, as long as they fall in (0, 1). However, approximation 33 may be inappropriate if this is not the case.

## 4 Relation to other methods

A wide range of approaches have been proposed for the problem of high-dimensional association testing using an informative covariate. Given the correspondingly wide variation in problems of this type, the optimal method is likely to depend on circumstance. In general, we will take *P, p*, 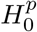 to refer to p-values and hypotheses for the trait of primary interest, and *Q, q* to refer to the covariate.

### 4.1 Determination of rejection region form

The simplest approach to covariate-based testing is ‘independent filtering’ [4] in which attention is restricted to the set {(*p_i_, q_i_*) : *q_i_* ≥ *q*_0_}, with the B-H procedure then applied to the corresponding subset of values of *p_i_*. This procedure is equivalent to rejection regions which are a series of rectangles with upper border at *q* = *q*_0_. Independent filtering is clearly non-optimal, but is well-suited to some problem types [4].

As discussed above, a range of approaches aim to approximate the optimal rejection regions based on *f*_0_/*f*. In [1] and [3], parametrisation leads to rejection regions constricted to a particular parametric class; in [1] that of oracle rejection regions under mixture-Gaussian forms of *f*_0_ and *f*. In [2] and [44], boundaries of rejection regions are necessarily smooth at a scale corresponding to the smoothing kernel width, but can take otherwise arbitrary forms. An alternative approach is to ‘bin’ covariates [5, 13] which leads to L-curves which are step-functions with steps spaced according to the resolution of the bins.

An approach in [11] estimates 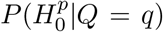 for each *q* to modulate a B-H type test for each observation. The entire effect of the covariate in this method is encompassed through the value of 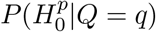, which necessarily relies on point-estimates of the PDF 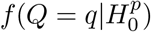, and hence the method is dependent on the accuracy of this estimate.

Another common approach to covariate modulation is the weighted Benjamini-Hochberg procedure [7], in which each p-value *p_i_* is reweighted to a value *p_i_/w_i_* (where ∑ *w_i_* = 1) and the standard B-H procedure is then applied to the values *p_i_/w_i_*. Our method can be interpreted in these terms, setting *w_i_* = *v_i_/p_i_*, but this is rather unnatural; there is no clear way to interpret what the ratio *v_i_/p_i_* means, and this approach does not make use of the ‘p-value property’ in equation (20).

The use of empirical CDFs to generate rejection regions has the advantage of making use of the global distribution of *P*, *Q*, while spline- and kernel-density based estimates can generally only use local observations. The cFDR-based method has the obvious disadvantage of not converging to the optimum rejection region, and it can be less powerful than parametric approaches if parametric assumptions hold. However, using CDFs rather than PDFs allows faster convergence of rejection regions with *n*, and this favours the cFDR approach if *n* is small, particularly if CDF- and PDF-based regions are similar and PDFs are difficult to model well.

Under certain circumstances, contours of CDF- and PDF-based methods are similar. A precise statement, proof, and demonstration is given in appendix 8.5.

### 4.2 Censoring of points

In general terms, the process determining a decision rule to be used on observation (*p_i_, q_i_*) cannot easily make use of the datapoint (*p_i_, q_i_*) itself, since the use of the point biases the choice of decision rule in some way. Approaches by [1, 2] censor the points used in the decision rule to those already rejected in a stepwise approach, and a method in [3] masks the information available for the decision rule by effectively adding the point (1 – *p_i_, q_i_*) to the dataset.

Since cFDR uses the entire dataset to estimate empirical CDFs, complex censoring can require that the cFDR estimator be changed in a non-trivial way. In particular, there is no obvious way to apply the methods proposed by [2] or [3]. We propose avoiding the problem by leaving out the point (*p_i_, q_i_*) directly (equations (25),(26)), at the cost of residual correlation in resultant v-values. While crude, this corresponds to a near-minimum censorship of points, and the resultant correlation tends to be small enough to ignore (see appendix 8.4).

### 4.3 Asymmetry and management of extreme outliers

An important property of the cFDR-based method is asymmetry, in that 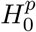 cannot generally be rejected based on a low *q_i_* alone (this can be seen by noting that 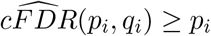 and *p_i_* can only exceed 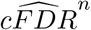 in rare circumstances). Parametric approaches such as those in [1, 3] are not generally robust to this; for example, in [1], an extremely low *q_i_* could lead to rejecting 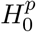 even if the corresponding *p_i_* were close to 1 and *P*, *Q* were independent (given that the degree of dependence is estimated). This property of the cFDR is very important when *p_i_* and *q_i_* are derived from GWAS on different diseases; it is entirely possible and even expected that a very strong association in the conditional trait is not an association with the principal trait. This property also differentiates our approach from meta-analysis of two sets of p-values.

### 4.4 Relation to original FDR-controlling method for cFDR

In a paper in 2015 [23], we identified the problem of failure of FDR control at *α* when using a rejection rule 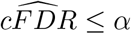 and proposed a rough solution. We proposed identifying L-curves and estimating f_0_ as above, and for each L-region *L_S_*(*α\**), identifying a rectangle *R*(*α\**) contained within it with vertices (0, 0), (0, *q_r_*), (*p_r_*, 0), (*p_r_, q_r_*). Since *R*(*α\**) ⊆ *L_S_*(*α\**), we have

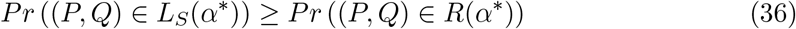

and 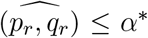, so the FDR associated with rejecting any (*p*, *q*) pairs falling in *L_S_*(*α\**) was approximately

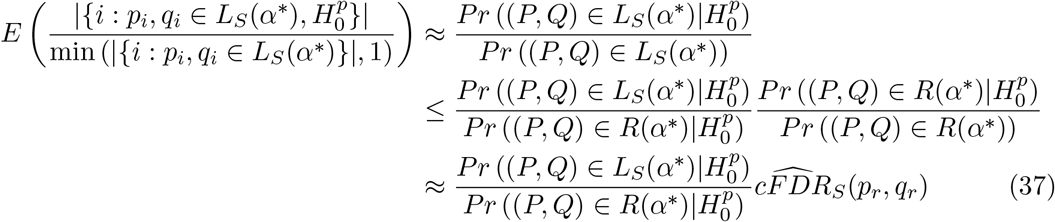

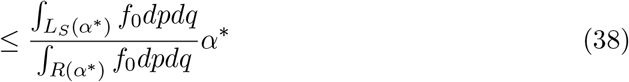

To approximately control FDR at *α*, our procedure found *α\** so that expression (38) was ≤ *α* and rejection 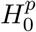 whenever (*p_i_, q_i_*) ∈ *L_S_*(*α\**).

As well as being approximate, this procedure was conservative due to inequality (36). Our new method avoids this conservative assumption, and is on firmer theoretical ground. Furthermore, our old method precluded use of 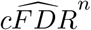 given approximation (37). We show by simulation below that this results in substantial improvement in power in our new method.

## 5 Assessment of performance

In this section, we address five main points. Firstly, we demonstrate that our new method controls type-1 error rate (FDR) appropriately, and that the censoring approach of (25) and (26) is necessary. Secondly, we demonstrate that power is substantially improved relative to our previous method for fixed level of FDR control, and that use of 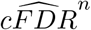 over 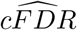 improves power further. We then demonstrate that in settings where parametric assumptions are not satisfied, rejection regions based on 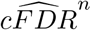 can correspond to a more powerful procedure than rejection regions based on alternative CDF or PDF estimators. We examine the effect of correlation between observations *p_i_, q_i_* on our main FDR-controlling methods, and demonstrate that the disadvantage of using our leave-one-out method (equation 25) instead of the leave-out-block method (equation 26) out method in the presence of correlation is loss of power rather than loss of FDR control. Finally, we assess the degree of shared association between *P* and *Q* which is necessary for our method to give an advantage over p-values alone.

In each simulation, we sampled a set of values *S* = (*p_i_, q_i_*), *i* ∈ 1..*n*. The sampling schema we used itself depended on a series of underlying parameters, which were themselves sampled from a joint distribution specified in 1, or a conditional distribution of it. We also separately considered several fixed values of parameters.

We first chose a fixed total number of hypotheses *n*, then split these into four classes of fixed size: *C*_1_ of size 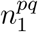 associated in both *P* and *Q*, *C*_2_ of size 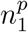 associated only with *P*, *C*_3_ of size 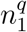 associated only with *Q* and *C*_4_ associated with neither *P* nor *Q*. Within each class, samples (*p_i_, q_i_*) were identically distributed.

For *i* ∈ *C*_1_, *C*_2_, we sampled *p_i_* (determined by *d*, *s_p_*) by first simulating Z scores:

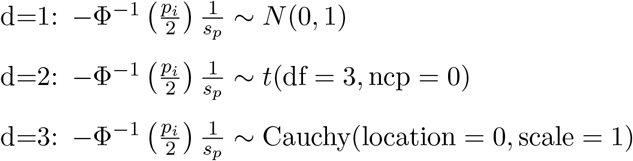

where 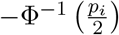 can be considered a *Z*-score corresponding to *p_i_*, and *s_p_* a scaling factor for the distribution. We set the distribution of *q_i_* ∈ *C*_1_, *C*_3_ similarly, with *s_q_* in place of *s_p_*. The values *p_i_*, *q_i_* for *i* ∈ *C*_4_ were sampled from *U*(0, 1).

Although effect sizes are often assumed to follow normal distributions, real data is often noisier, with longer tails, and recent work suggests non-normal distributions may be a better fit in the case of GWAS data [48]. We chose the alternative distributions (normal, t (3df), and Cauchy) to span behaviours from ‘well-behaved’ (normal) to ‘long-tailed’ (t) to ‘chaotic’ (Cauchy) to survey a wider range of possibilities.

Samples *p_i_*, *p_j_* and *q_i_, q_j_* were generally independent (unless otherwise specified), but we also sampled under two patterns of dependence. Firstly, we simulated a ‘block’ correlation structure in which we divided samples into three blocks, and within each block sampled z-scores *z_p_i__*, *z_p_j__*, *z_q_i__*, *z_q_j__* corresponding to *p_i_*, *p_j_* and *q_i_, q_j_* such that cor(*z_p_i__, z_p_j__*) = cor(*z_q_i__, z_q_j__*) = *ρ* if *i, j* were in the same block and class, and cor(*z_p_i__, z_p_j__*) = cor(*z_q_i__, z_q_j__*) = 0 otherwise. Secondly, we simulated an equicorrelation structure in which cor(*z_p_i__, z_p_j__*) = cor(*z_q_i__, z_q_j__*) = *ρ* whenever *i* and *j* were in the same class. When *d* ∈ 2, 3, we used the off-diagonal elements of the normalised dependence matrix in the multivariate T distribution in place of correlation.

When relevant, we also sampled parameters from the distribution specified in table 1 conditional on 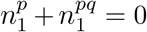; that is, no associations with *P*. We plotted results from these simulations separately to those with parameters drawn from the unconditional distribution.

**Table 1:** Variables used in simulations

| Variable | Description | Sampling distribution |
| --- | --- | --- |
| $n$ | Total number of variables | $10^{U(3,4)}$ (rounded) |
| $n_1^{pq}$ | Number of variables assoc. with $P, Q$ | $U(0, 200)$ (rounded) |
| $n_1^p$ | Number of variables associated with $P$ | $U(0, 200)$ (rounded) |
| $n_1^q$ | Number of variables associated with $Q$ | $U(0, 200)$ (rounded) |
| $s_p$ | Scale for distribution of $P$ (see below) | $U(\frac{3}{2}, 3)$ |
| $s_q$ | Scale for distribution of $Q$ | $U(\frac{3}{2}, 3)$ |
| $d$ | Form of alternative distributions | 1: Normal, 2: t (3df), 3: Cauchy, eq. prob. |

Given a rejection procedure, we defined

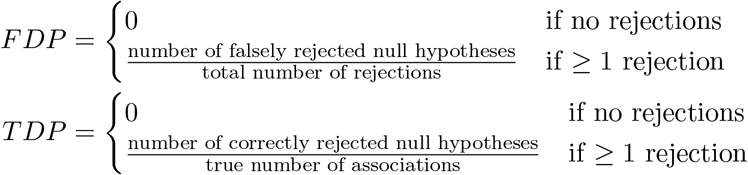

We analysed type 1 error in terms of the estimated FDR, 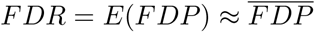, and power in terms of the corresponding true-discovery rate 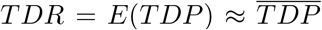. We compared *FDP* and *TDP* between samples by estimating them via a Gaussian-weighted moving average across the independent variable (usually 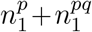). We show 95% pointwise confidence envelopes derived as per [49], except in cases where such envelopes obstruct viewability of the plot. In these cases, we state that values *TDR*(*A*) for one method *A* ‘exceed’ paired values *TDR*(*B*) of another method *B* if in at least six of eight equal subdivisions of the x-axis range the following three conditions hold: *TDR*(*A*) > *TDR*(*B*) more than *TDR*(*B*) > *TDR*(*A*), the mean 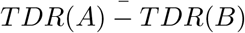 is positive, and a Wilcoxon test of ranks on *TDR*(*A*) − *TDR*(*B*) rejects the null hypothesis of a symmetric distribution around 0 with *p* < 5 × 10^−3^ (or the equivalent with *FDR* in place of *TDR*). Each line on each plot corresponds to *>* 5000 simulation runs. All raw simulation results and analytic code are publically available at https://github.com/jamesliley/cfdr_pipeline.

### 5.1 New FDR-controlling procedure leads to greater power than previous method, and adjustment improves power further

We first compared FDR control amongst five methods, aiming to control the FDR at either *α* = 0.1 or *α* = 0.01:

1. the B-H method applied to the values *p_i_*, labelled ‘P-val’
2. the B-H method applied to ‘naive’ v-values *v*(*p_i_, q_i_*) = *v_S_*(*p_i_, q_i_*) as per equation 27 for reference, labelled ‘Naive’
3. the B-H method applied to ‘leave-one-out’ v-values *v*(*p_i_, q_i_*) = *v*_*S*−(*p_i_, q_i_*)_(*p_i_, q_i_*) as per equation 25, labelled ‘LOO’
4. the B-H method applied to block-out v-values (after randomly separating observations into three equally sized subdivisions, so (*p_i_, q_i_*) is in subdivision *b*(*i*), defining v-values *v*_*S−b*(*i*)_(*p_i_, q_i_*)) as per equation 26, labelled ‘LOB’
5. our previous method for FDR control applied to (*p_i_, q_i_*), labeled ‘Orig.’

We sampled simulation parameters according to table 1, or the corresponding conditional distribution of table 1 with 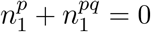.

Expected *FDP* was consistent with the FDR control level when using leave-one-out v values or ‘block-out v-values (rejection procedures 3,4). When using the ‘naive’ v-values *v_S_*(*p_i_, q_i_*) (rejection procedure 2), FDR was not controlled at the requisite level. The FDR using methods 3 and 4 exceeded the FDR of our original method (rejection procedure 5), indicating that our original method was conservative. FDR control was maintained when using the approximation of *f*_0_ in equation (33). Results are shown in figure 2.

**Figure 2:**
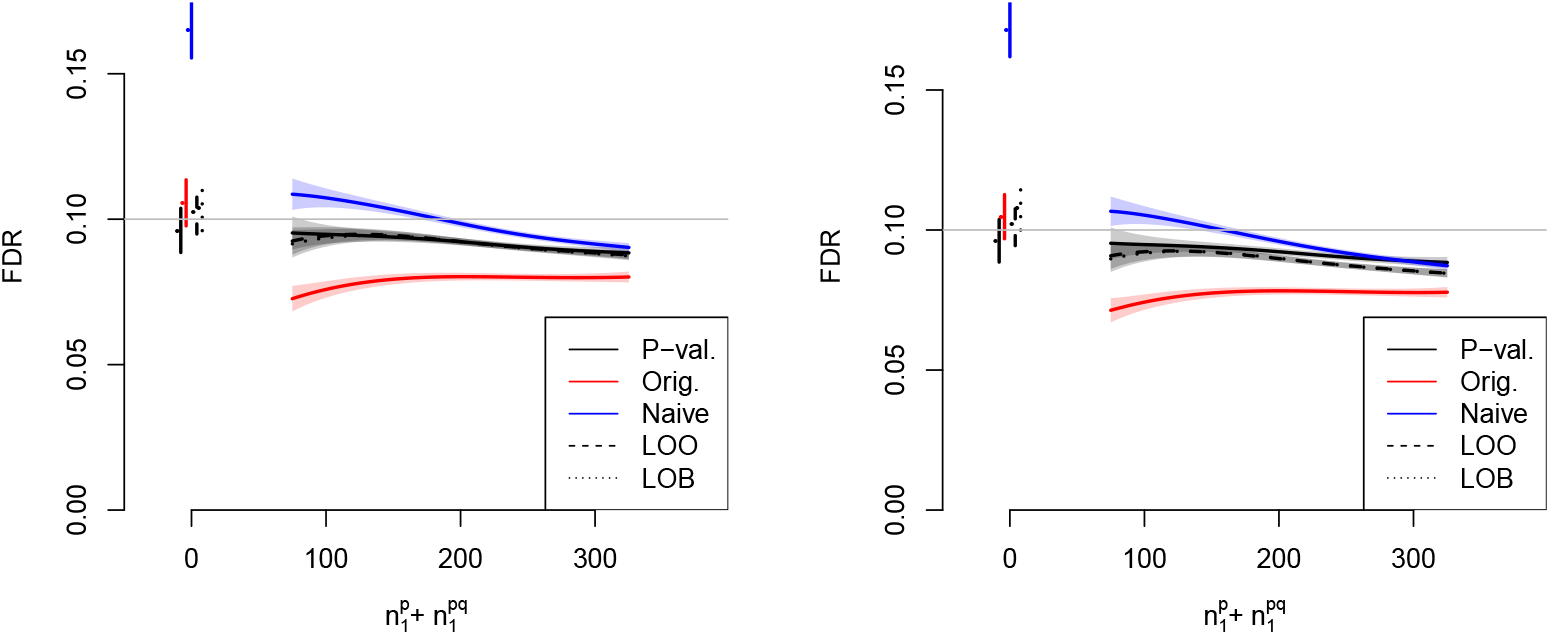
FDR control of various methods against 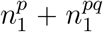, the total number of variables associated with *P* (the primary study under consideration). The horizontal line shows *α* = 0.1, the desired FDR control level. Simulations in the left panel integrate L-regions over the the true distribution *f*_0_; simulations in the right panel integrate over the estimated distribution as per equation (33). Shaded regions indicate 95% confidence envelopes. Curves show moving weighted averages using a Gaussian kernel with SD 15% of the X axis range. Lines on the left indicate FDR control with 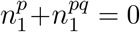. A corresponding plot with *α* = 0.01 is shown in supplementary figure 19.

Having established the validity of rejection methods 3, 4, we compared the power of ‘adjusted’ 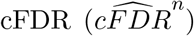 and non-adjusted cFDR 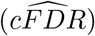 using the leave-one-out v-value (rejection procedure 3) and the power of our previous method, rejection procedure 5, applied to 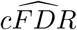 (labelled ‘Orig’). The TDR of 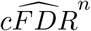 exceeded the power of 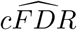, which in turn exceeded the power of our previous rejection procedure on 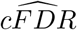 (figure 3).

**Figure 3:**
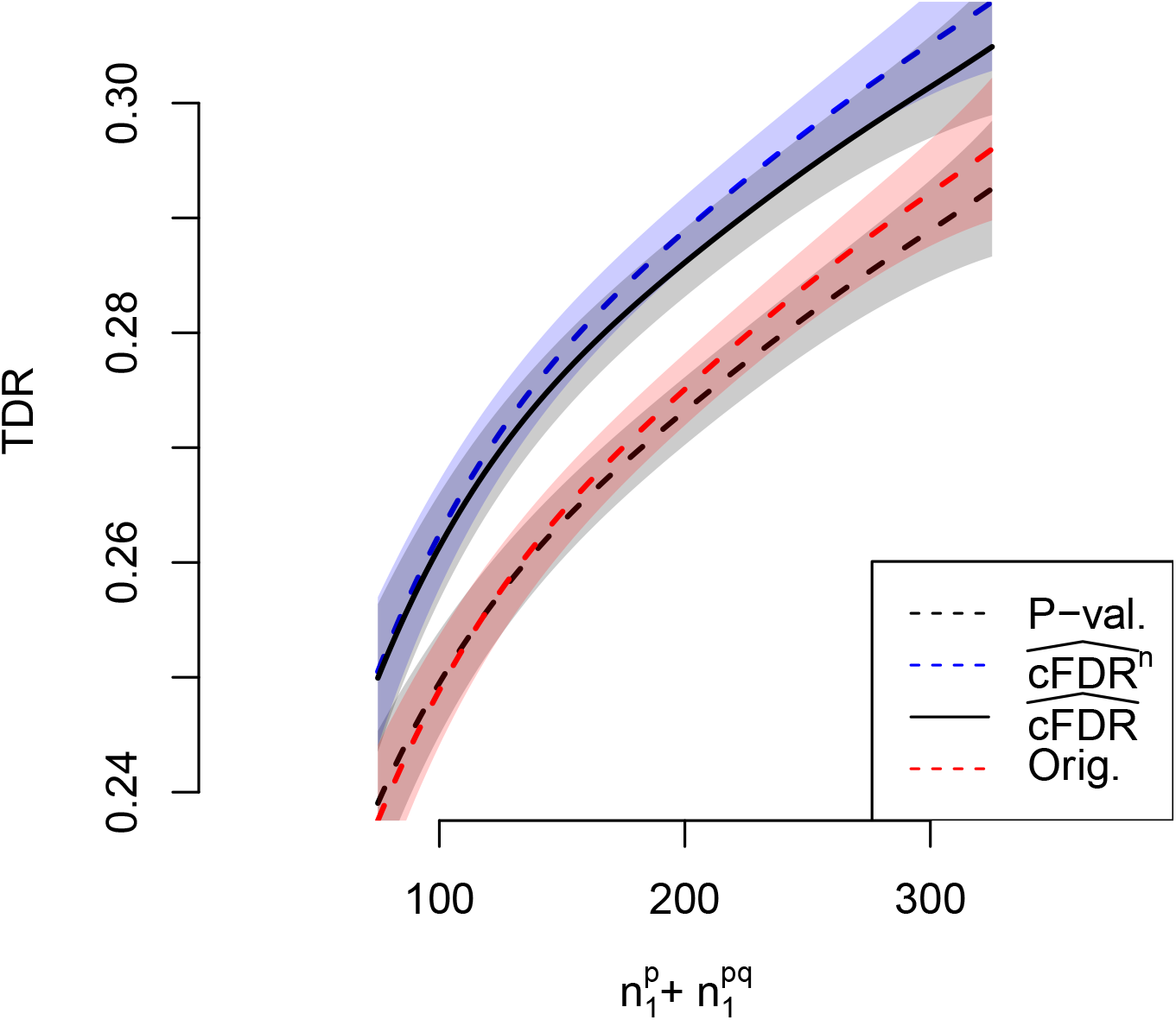
TDR of various methods against 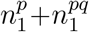, the total number of variables associated with *P* (the primary study under consideration), at FDR control level *α* = 0.1. Shaded areas show 95% pointwise confidence envelopes. A corresponding plot with *α* = 0.01 is shown in supplementary figure 20. Curves show moving weighted averages using a Gaussian kernel with SD 15% of the X axis range.

We report FDR and TDR for p-values, 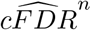 and oracle cfdr for a range of fixed simulation parameters in table 2

**Table 2:**
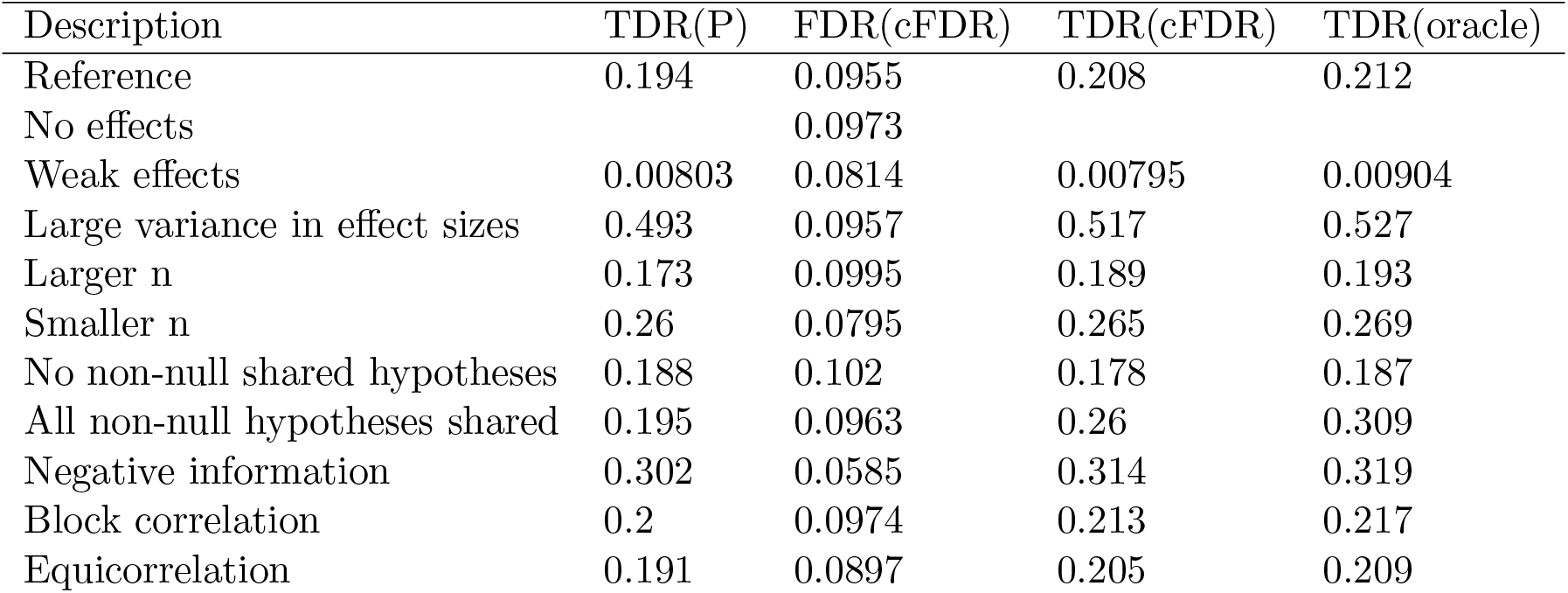
FDR and TDR of p-value, 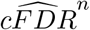, and oracle cfdr (best possible procedure) using leave-one-out v-values (equation 25) for a range of simulation parameters, controlling FDR at *α* = 0.1. All descriptions are relative to ‘Reference’ which has the following parameter values: 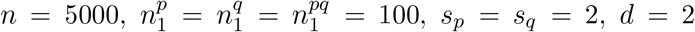, *s_p_* = *s_q_* = 2, *d* = 2. TDR is undefined if 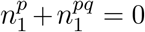. ‘Negative information’ means *fewer*-than random shared associations, rather than more 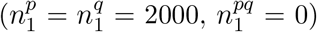. Complete parameter values, confidence intervals, and more detailed results are shown in supplementary table 3.

### 5.2 PDF- based estimation leads to a less powerful procedure than CDFbased estimation

As *n* → ∞, consistent estimators of 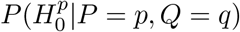 will converge to optimal rejection regions while estimators of 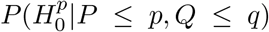 will not, and hence the former will ultimately be more powerful. However, we found that under the distribution of simulation parameters in table 1, the ECDF-based estimator 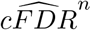 is considerably more powerful than two PDF-based estimators of 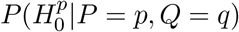.

Results are shown in figure 4. We considered parametric (labelled ‘PDF param’) and KDE-based (labelled ‘PDF KDE’) estimators of 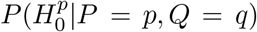. The parametric model was based on a four-Gaussian model detailed in supplementary material, section 9.4. We separated cases in which parametric assumptions were satisfied (ie *d* = 1 in table 1) and in which they were not (*d* = 2, 3). The TDR of 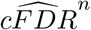 exceeded the TDR of both estimators of 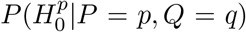. The performance of an oracle CDF procedure (using exact contours of *F*_0_/*F* as rejection regions, labelled ‘CDF oracle’) and an oracle PDF procedure (using exact contours of *f*_0_/*f* as rejection regions, labelled ‘PDF oracle’) are shown for comparison.

**Figure 4:**
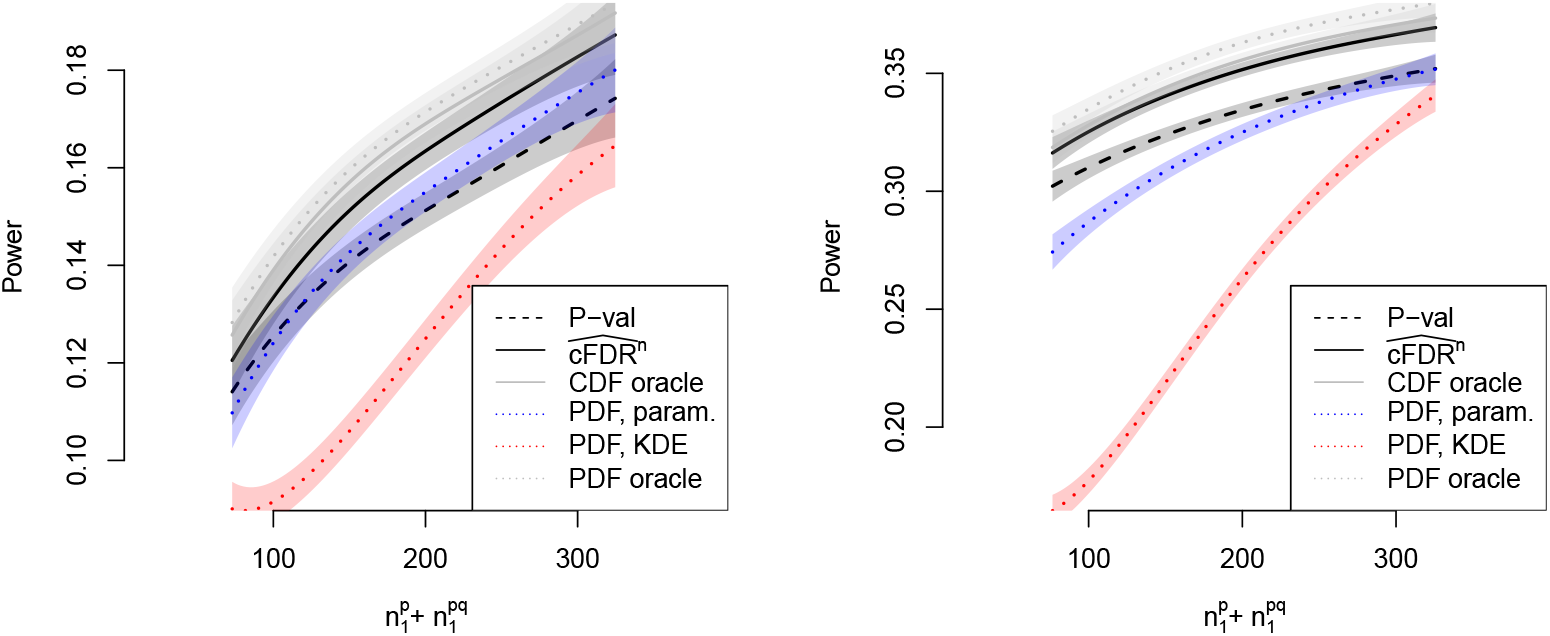
TDR of PDF-based methods against 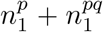, the total number of variables associated with *P* (the primary study under consideration), controlling FDR at *α* = 0.1. In the left panel, parametric assumptions were satisfied (ie *d* = 1 in table 1) and in the right panel, they are not (*d* = 2, 3). Shaded regions show pointwise 95% confidence intervals. A corresponding plot with *α* = 0.01 is shown in supplementary figure 22. Curves show moving weighted averages using a Gaussian kernel with SD 15% of the X axis range.

#### 5.2.1 Parametric- and KDE- based cFDR estimators are less powerful than the ECDF-based estimator

We also examined PDF- and KDE- based estimates of the cFDR rather than the cfdr. Details of the alternative estimators are given in supplementary material, section 9.4.

When parametric assumptions were satisfied (figure 5, left panel), performance of the ECDF-based 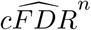, parametric (labelled ‘CDF param’) and KDE-based (labelled ‘CDF KDE’) cFDR estimators was equivocal. When parametric assumptions were not satisfied (*d* = 2, 3 as per table 1; figure 5, right panel), the TDR of the ECDF estimator exceeded the TDR of the the parametric and KDE estimators. The performance of an oracle CDF procedure (using exact contours of *F*_0_/*F* as rejection regions) is shown for comparison.

**Figure 5:**
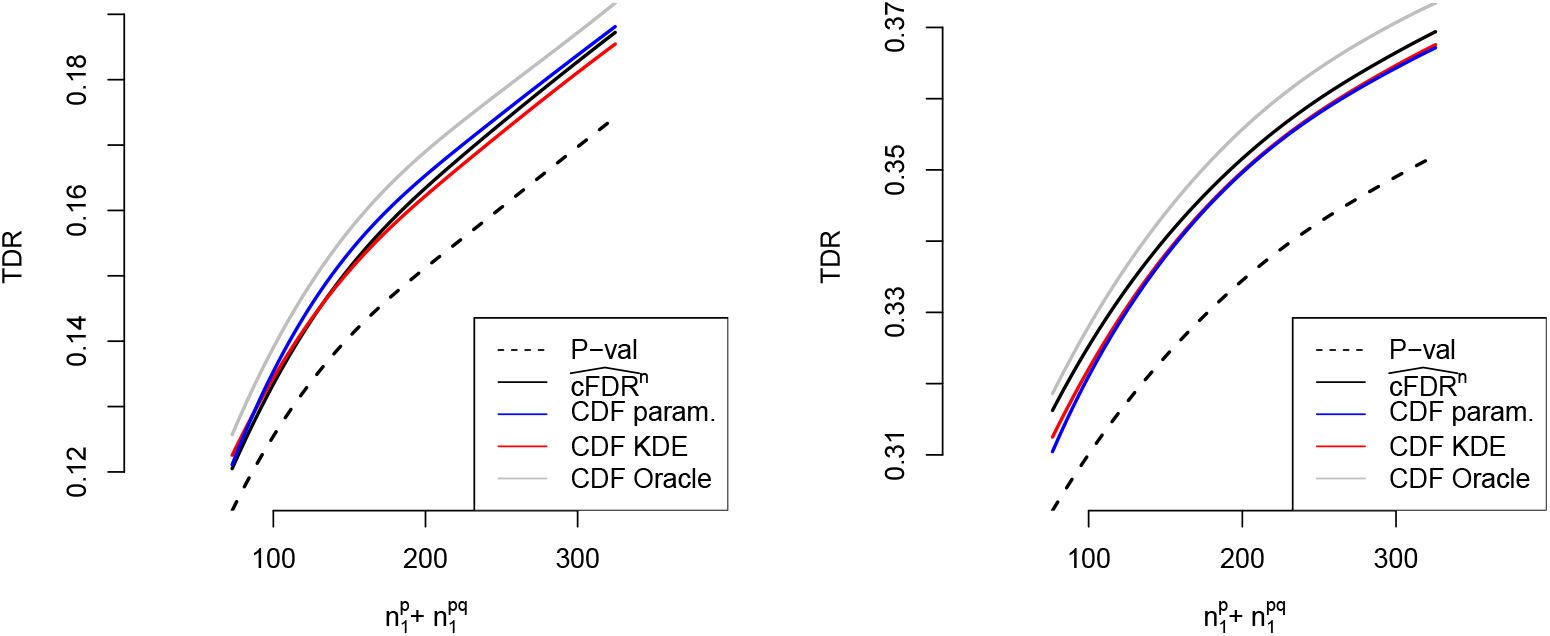
TDR of various methods against 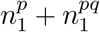, the total number of variables associated with *P* (the primary study under consideration), restricting to simulations in which parametric assumptions were satisfied (left panel) or were not satisfied (right panel), at FDR control level *α* = 0.1. A corresponding plot with *α* = 0.01 is shown in supplementary figure 21. Confidence intervals are omitted for visual clarity Curves show moving weighted averages using a Gaussian kernel with SD 15% of the X axis range.

### 5.3 Correlated samples lead to lower TDR but FDR control is maintained

When *p_i_, q_i_* had either block correlation or equicorrelation, we found that FDR control was maintained when using leave-one-out v-values (equation 25) and when using leave-out-block v-values (equation 26. Under equicorrelation, the TDR of leave-out-block exceeded the TDR of leave-one-out v-values. Under block correlation, although TDR of leave-out-block did not formally exceed the TDR of leave-one-out, a paired Wilcoxon rank-sum test on TDR values rejected the null hypothesis of a symmetric distribution around 0 with *p* < 1 × 10^−6^ in favour of leave-out-block.

Maintenance of FDR control is expected, as the Benjamini-Hochberg procedure controls FDR more conservatively when p-values are positively correlated than when independent. Figure 6 shows FDR and TDR controlling at *α* = 0.1 in the case *ρ* = 0.01, including performance of p-values *p_i_* under the BH procedure for comparison. The case *ρ* = 0.1 is similar and is shown in supplementary figure 23

**Figure 6:**
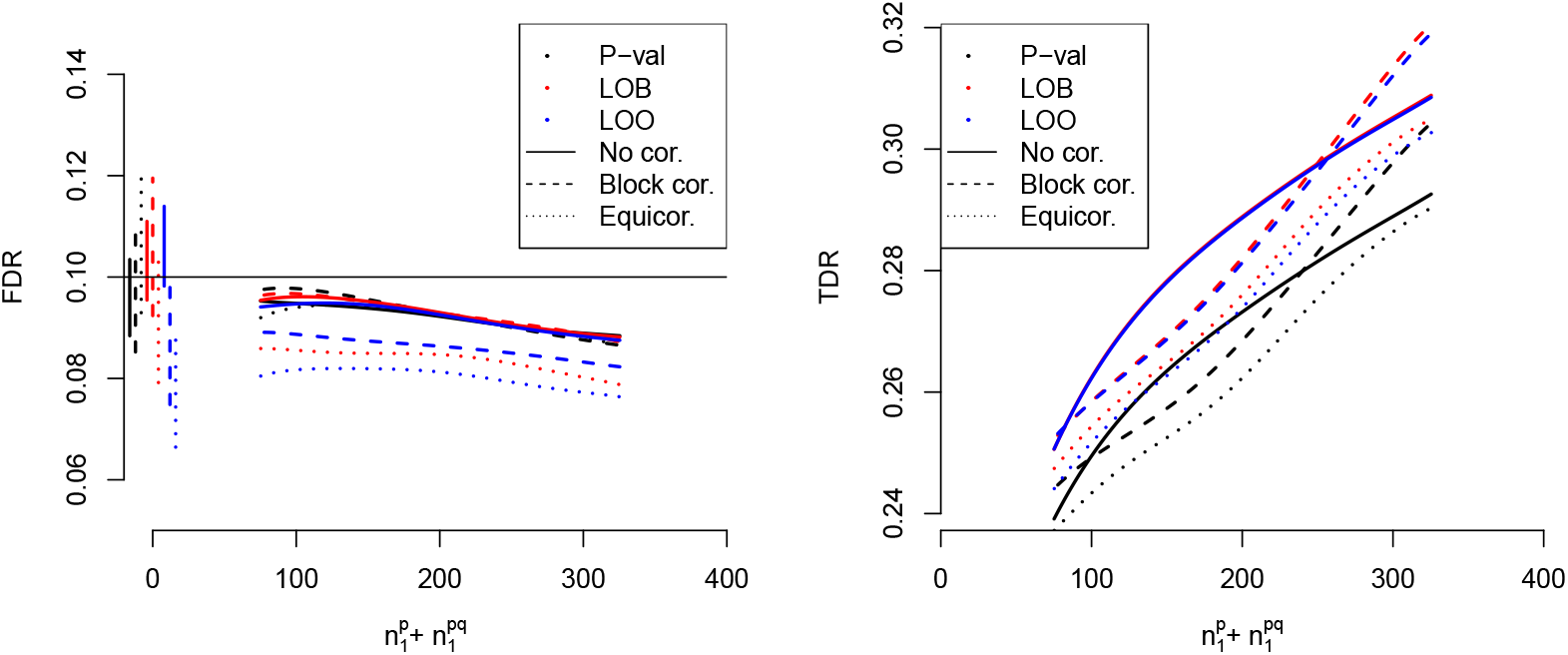
FDR (left) and TDR (right) of FDR-controlling methods leave-out-block (equation 26) and leave-one-out (equation 25) applied to 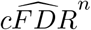, and the BH procedure applied to p-values, under several models of correlation between observations (*ρ* = 0.01). Confidence envelopes are omitted for visual clarity. Vertical lines show FDR with 95% confidence intervals at 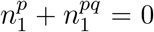 (the p-value appears not to control FDR under equicorrelation, but it is well-known to do so theoretically). A corresponding figure with *ρ* = 0.1 is shown in supplementary figure 23. Curves show moving weighted averages using a Gaussian kernel with SD 15% of the X axis range.

### 5.4 TDR of 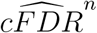 becomes higher than TDR of p-value alone when ≈ 20% of hypotheses are shared

Finally, we assessed the proportion of non-null hypotheses for *P* which needed to be shared with *Q* in order for 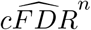 to have an advantage in TDR over only considering *P*. When no non-null hypotheses are shared, the values *q_i_* confer no information on 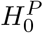, so we expect that use of v-values will add only noise and the TDR of the p-value will be larger than that of the v-value. We plotted the difference in TDR between 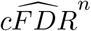 (using leave-one-out v-values) and p-values *p_i_* against 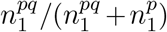, and found that this difference became positive when around 20% of hypotheses were shared (figure 7). This figure is dependent on our simulation parameters: a smaller percentage of hypotheses may lead to an advantage of 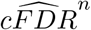 if, for instance, effect sizes were very large.

**Figure 7:**
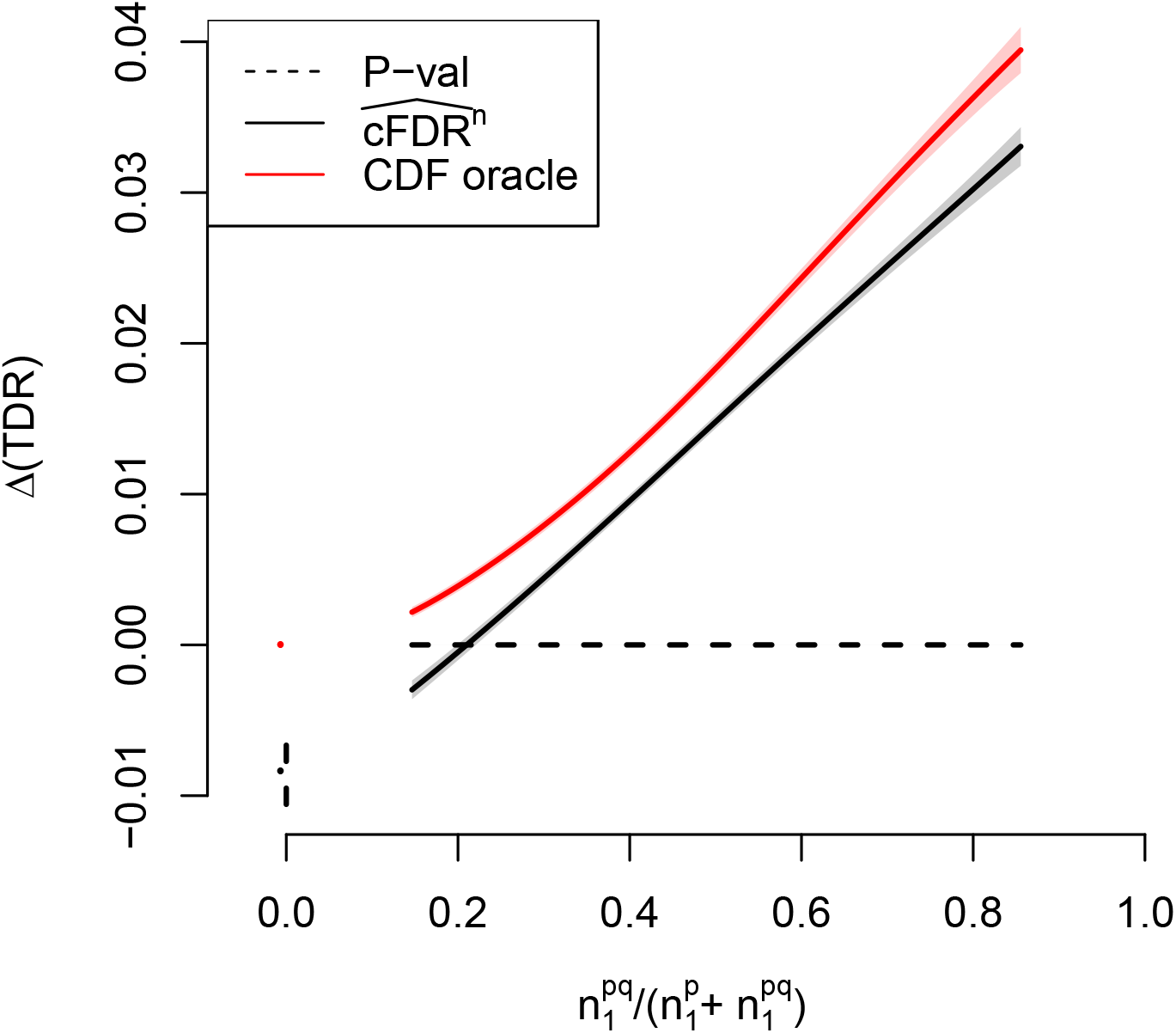
Difference in TDR between 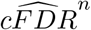 (assessed by leave-one-out v-values) and p-values, controlling FDR at *α* = 0.1, against 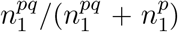 (proportion of non-null hypotheses for *P* which are shared with *Q*). The performance of the oracle CDF method is shown for comparison. Shaded areas show pointwise 95% confidence intervals. Points and lines at the leftmost edge show TDR values and 95% confidence intervals when 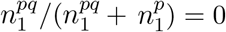. A corresponding figure with *α* = 0.01 is shown in supplementary figure 24. Curves show moving weighted averages using a Gaussian kernel with SD 15% of the X axis range.

### 5.5 Iterated cFDR

Since our proposed method for type-1 error rate control maps p-value/covariate pairs to v-values preserving the p-value property, we are free to use the resultant v-values in a second cFDR-based analysis against a second covariate. This enables immediate and simple adaptation to a setting in which more than one set of covariates are available. In our motivating example, this would allow us to subsequently ‘condition’ on other potentially related diseases as well as OCA.

We simulated a set of p-values {*p*} = {*p_i_, i* ∈ 1..1000}, with 100 true associations 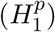 in which p-values were sampled from 2Φ(−|*N*(0, 3^2^)|) (where Φ is the normal CDF) and 900 non-associations 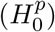 in which p-values were sampled from *U*(0, 1). We then similarly simulated sets of covariates 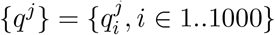 with 100 true associations 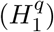, which for even *j* were randomly spaced amongst the 1000 variables (uninformative covariates) and for odd *j* overlapped more-than-randomly with associations with principal p-values (informative covariates), with around 54 shared associations on average (strictly, such that 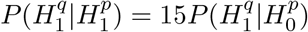).

Starting with *v*_0_ = *p*, we conditioned on each set of {*q^j^*} in succession, so *v_i+1_* = *v*(*v_i_, q^i^*). We used 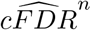 as an estimator and used leave-one-out v-values. Originally 19 of 100 null hypotheses were correctly rejected using *p* alone (*p* < 5 × 10^−5^ = 0.05/1000. On repeated conditioning, almost all all v-values when *H^p^* = 1 tended to 0: 99 null hypotheses were correctly rejected using *v*_500_. Under 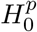, v-values remained uniform on (0, 1) (figure 8). This indicated the potential to greatly strengthen the power of a highdimensional association analysis by repeated conditioning in this manner, even when only half of the sets of covariates are informative.

**Figure 8:**
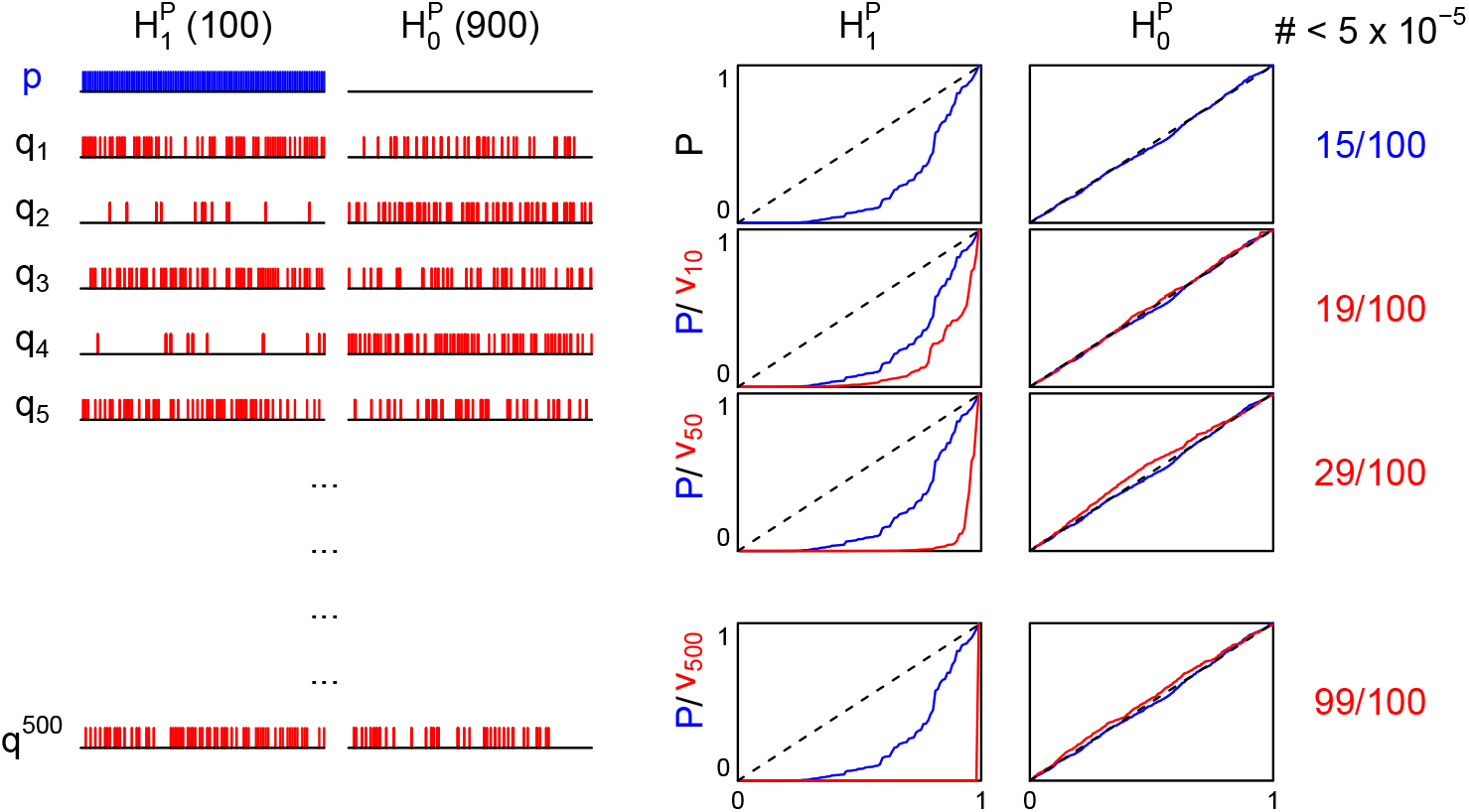
Iterated cFDR. Left part of plot shows where non-null hypotheses fall in *p/q^j^* (blue vert. lines for *p*, red for *q_j_*). Non-null hypotheses are shared more-than-randomly in only every second set *q^j^*. Right part shows *p/v_j_* values (blue/red lines respectively) plotted in ascending order under 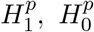, and the number of *p/v_j_* values which reach Bonferroni-corrected significance.

### 5.6 Summary of BRCA analysis

Finally, we return to the motivating example. cFDR rejects more null hypotheses for BRCA (724) than B-H on BRCA data alone (678, figure 1A) or the subset of variables with OCA association (280, figure 1B). The procedure is asymmetrical in that it will not reject a BRCA null hypothesis for a low OCA p-value alone, and can readily be reversed: supplementary figure 18 shows a similar analysis analysing association with OCA.

## 6 Discussion

We present an improvement to the conditional false discovery rate method, a widely-used procedure in genetic discovery. Our new methods essentially involve computing an analogy of the p-value corresponding to the ranking of hypotheses defined by the cFDR estimator. Our method enables the cFDR to be used definitively in the discovery phase of -omics studies with control of a type-1 error rate. The general procedure of multiple p-value testing with a covariate has wide scientific application; see [1, 2, 11] for examples.

The 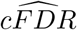 and 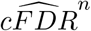 estimators make no distributional assumptions on *P*, *Q*. The type-1 error rate controlling method requires modelling of the PDF of *P*, 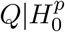, but this requires approximating a univariate PDF 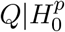. Furthermore, this PDF is only used as an integrand rather than for direct point-estimates. It is reasonable to expect that for approximations to complex PDFs, relative average errors over intervals will be smaller than relative errors at individual points; parametric approximations tend to be smoother than the true distribution at a fine scale, and KDE-based approximations rougher. An obvious shortcoming of cFDR-based methods is the lack of asymptotic optimality. Methods based on consistent estimators of *f*_0_/*f* will eventually outperform any estimator of *F*_0_/*F* for large enough *n* (see supplementary material, section 9.5). However, the ECDF-based cFDR estimator was far stronger than PDF-based estimators at the values of *n* we simulated at (10^3^ − 10^4^). In practical terms, it is important to note that n, being the number of variables, cannot generally be increased indefinitely, as opposed to, for instance, sample size. Essentially, the fitting of L-curves corresponds to a procedure by which the similarity between *P*, *Q* is assessed, and the degree of modulation when moving from *p* to *v* values corresponds to this similarity. Moreover, this assessment of similarity occurs intrinsically on the basis of the joint CDF rather than relying on a parametric description.

L-curves may not change monotonically with *Q*; that is, it may be possible to reject a null hypothesis with p-values (*p*, *q*_1_) and not reject a null hypothesis with p-values (*p*, *q*_2_), *q*_2_ < *q*_1_ (see the lower-left panel of figure 1). It would be possible within our framework of FDR control (theorem 1) to force L-curves to be monotonic with *Q*, and indeed since this would result in straight-up-and-down segments on L-curves, the loss of power due to noise when *P* and *Q* are unrelated (figure 7) may be reduced in this case. However, nonmonotonicity of L-curves is potentially advantageous in a biostatistical setting. Between TWAS or GWAS for similar diseases, it may be the case that shared non-null hypotheses have ‘moderately’ small p-values, corresponding to common general shared medium-risk pathological causes, but non-shared non-null hypotheses have ‘extremely’ small p-values, corresponding to specific high-risk pathologies. Non-monotonic L curves allow this effect to be modelled. Unrestricted L-curves also allow use to be made of q-values such that 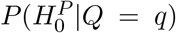 is lower with low *q*, rather than higher: we show this in table 2, row ’Negative information’.

Our proposed ‘iterated cFDR’ procedure can be thought of as a meta-analysis of a series of experiments *E_P_*, *E*_*Q*_1__, *E*_*Q*_2__, … giving rise to p-value sets 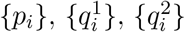, … when only the first set ({*p_i_*}) are known to test the correct hypotheses; that is, be *U*(0, 1) for null hypotheses. It enables us to find the set of non-null hypotheses corresponding to *E_P_* (denoted 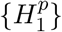), even though the set of non-null hypotheses corresponding to *E_Q_j__* (denoted 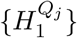) may only partly overlap 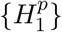, may contain hypotheses not in 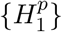, and (half the time) may even carry no information about 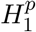 at all. This could be used to refine the set of association statistics {*p_i_*} for a disease of interest by using sets of association statistics 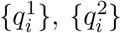, … at the same variables for a range of separate traits. It could also be used to improve power when repeating an -omics study in a new ethnic group by levering on previous studies in different ethnicities.

In summary, our method improves the power of cFDR analyses and allows it to be used confidently in the setting of multiple hypothesis testing. This can enable more efficient use of data, and more information to be gained from the same datasets. Our method contributes to a set of tools for high-dimensional statistical analysis and has wide application across a range of fields in biomedicine and elsewhere.

## 7 Code availability

All functions necessary to apply the methods detailed in this work are available in the R package https://github.com/jamesliley/cfdr

A full pipeline to generate the results in this paper is available in the git repository https://github.com/jamesliley/cfdr.pipeline.

## 8 Acknowledgments

We thank Professor Sylvia Richardson for helpful comments on the structure of this work. CW is funded by the Wellcome Trust (WT107881) and the MRC (MC_UU_00002/4). JL was funded by WT107881 and by Johnson and Johnson, and for part of this work was on the Wellcome Trust PhD programme in Mathematical Genomics and Medicine, funded by the NIHR Cambridge BRC. The funders had no role in study design, data collection and analysis, decision to publish, or preparation of the manuscript.

## Appendix

### 8.1 Optimal procedure

In this section, we show the following result. This is not original; it is shown in various forms in (at least) [1, 2, 3].

#### Theorem 5.

*Let f*_0_ *and f*_1_ *be positive Lesbegue-integrable functions of* (*p*, *q*) *on some region* Ω. *Suppose a Lesbegue-measurable region R*_0_ *satisfies*:

1. 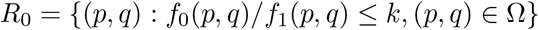
2. 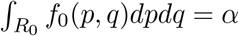
3. 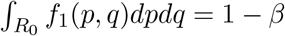

*Then no Lesbegue-measurable region R* ⊂ Ω *satisfies both*

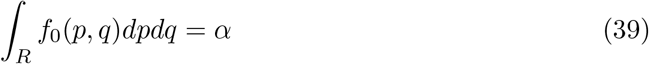

*and*

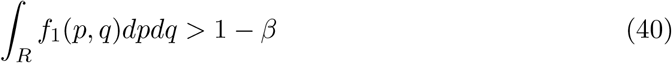

*Proof*. Suppose such a region existed. Then given condition 39, we must have *f*_0_(*p*, *q*)/*f*_1_(*p*, *q*) > *k* in *R* \ *R*_0_, and since the integral of *f*_0_ over *R* is equal to its integral over *R*_0_,

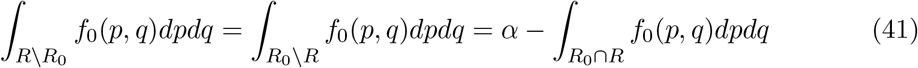

Hence

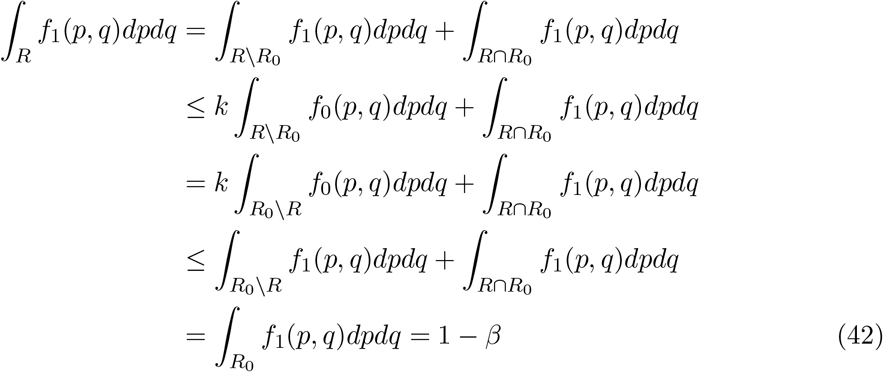

a contradiction of 40. Regions *R* ≠ *R*_0_ can satisfy 39 and

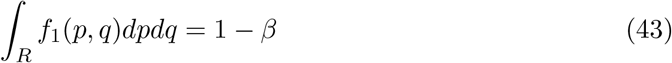

if and only if *R*_0_ \ *R* and *R* \ *R*_0_ have Lesbegue measure 0.

#### Corollary 5.1.

*If* 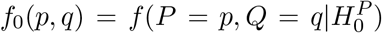 *and* 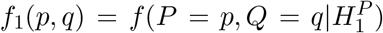 (*where* 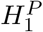 *is the alternative*) *then amongst all rejection regions R with fixed type-1 error rate* 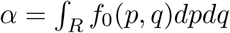, *power is maximised on a region inside a contour of f*_0_/*f*_1_, *if such a region exists*.

*Denoting f*(*p*, *q*) = *π*_0_*f*_0_(*p*, *q*) + (1 – *π*_0_)*f*_1_(*p*, *q*), *it is clear that a contour of f*_0_/*f*_1_ *is also a contour of f*_0_/*f and of* 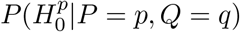, *so an optimal rejection region is given by*

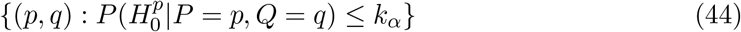

*for some k_α_*.

### 8.2 Failure of FDR control with 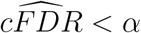

As described in section 2, rejection procedure (9) is similar to the B-H procedure, and it may be naively thought that it also controls the FDR at *α*. This is not the case, and indeed the FDR of such a procedure (and the corresponding procedure with 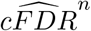) may exceed *α* by an arbitrary factor depending on *α* and *π*_0_.

This is most easily seen by considering the extreme case in which

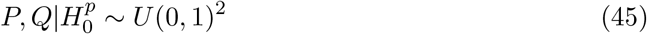

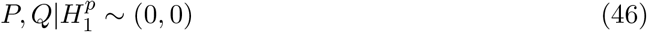

where 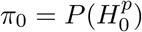 as usual. In this case we show:

#### Theorem 6.

*Under the above distribution of P, Q, as n* → ∞, *the FDR α_TRUE_ of rejection procedure* (9) *for* 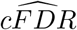 *satisfies*

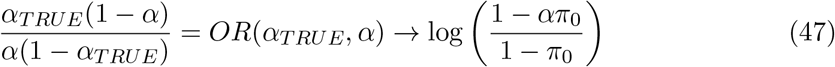

*and the corresponding procedure for* 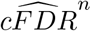 *satisfies*

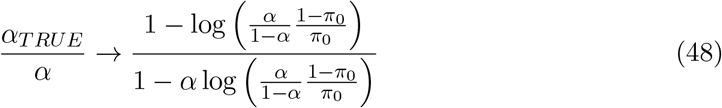

#### Corollary 6.1.

*For* 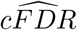, *the relative error in FDR (relative to α) can grow arbitrarily large as π*_0_ → 1, *α* → 0. *For* 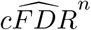, *the error can grow arbitrarily large as α* → 0, *regardless of π*_0_.

*Proof*. Suppose that we have a dataset *S* = {(*p_i_, q_i_*)}, *i* ∈ 1..*n* of draws from *P*, *Q* under 45,46. Due to assumption 45 we have 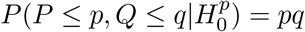 and due to 46 we have 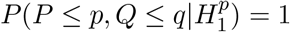. Now

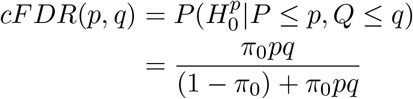

Now

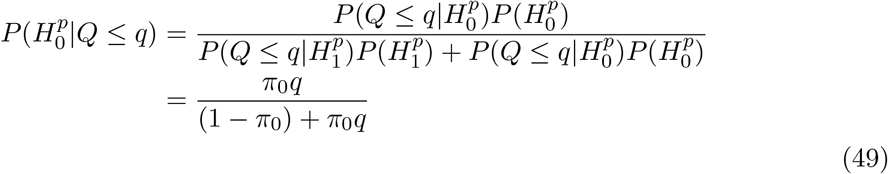

The estimate 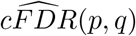 is proportional to a consistent estimator of

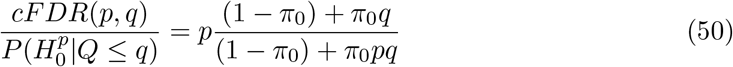

and since 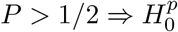, approximation 7 in the main paper is consistent, and 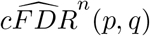 is a generally consistent estimator of *cFDR*(*p*, *q*).

The FDR of the rejection procedure 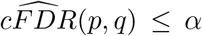 converges to the FDR of the rejection region 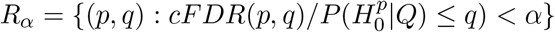 as *n* → ∞ (see diagram in figure 12). Since this rejection region contains (0, 0), all (1 – *π*_0_)*n* non-null hypotheses will be rejected, and the proportion of the total null hypotheses rejected will converge by the law of large numbers to

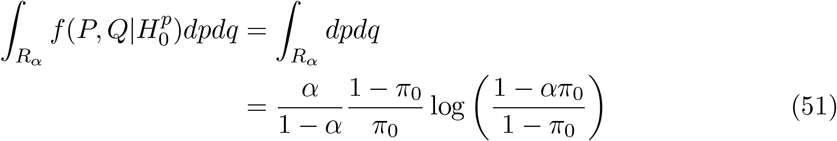

and thus the FDR *α_TRUE_* converges to

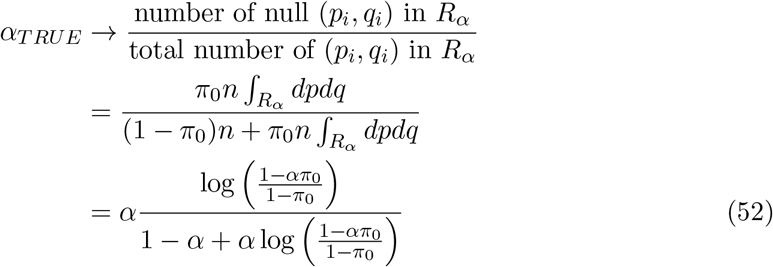

which can be written as

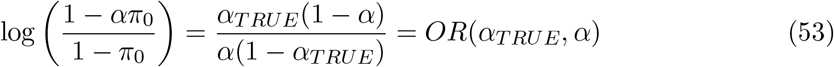

where OR is the odds ratio. Hence *α_TRUE_* can exceed *α* by an arbitrary degree when *pi*_0_ is close to 1.

The second part of the proof can be shown similarly. The RHS of equation 48 rises as −log(*α*) as *α* → 0, whatever the value of *π*_0_.

### 8.3 Convergence results

In these appendices, we omit the *X* from 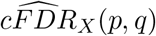 and other functions when it is clear. We consider *p*, *q* to be in (0, 1)^2^.

Set *n* = |*X*|, *F_n_*(*q*) = min(1,|{*i* : *q_i_* ≤ *q*, (*p_i_, q_i_*) ∈ *X*}|), *F_n_*(*p*, *q*) = min(1, |{*i* : *p_i_* ≤ *p, q_i_* ≤ *q*, (*p_i_, q_i_* ∈ *X*)}|), *F*(*q*) = *P*(*Q* ≤ *q*), *F*(*p*, *q*) = *P*(*P* ≤ *p*, *Q* ≤ *q*), and

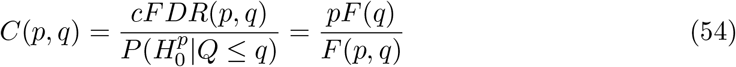

We will assume *∂F*(*p*, *q*)/*∂p* exists on (0, 1)^2^.

We define 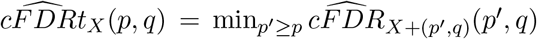, and 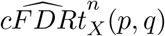 similarly for 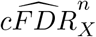 (‘t’ for ‘truncated’), so

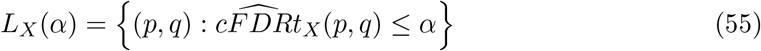

The boundary of *L_X_*(*α*) is continuous and piecewise-differentiable.

In this section, we show a series of results relating to convergence of cFDR estimates. We show results relating to convergence of 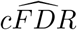 and 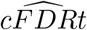 on a line *q* = *q*_0_, along with convergence of the co-ordinates of L-curves on such lines. We then show slightly weaker results regarding convergence across two-dimensional regions of the unit square.

#### Theorem 7.

*Suppose that on a line segment q* = *q*_0_, *p_γ_* < *p* < 1, *we have F*(*p*, *q*) ≥ *γ* > 0 *and F* (*q*_0_) > 0. *Then on this segment*, 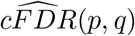 *converges uniformly to C* (*p*, *q*) *as n* → ∞. *If additionally we have ∂C*(*p*, *q*)/*∂p* ≥ 0, *then* 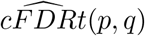 *converges uniformly to C*(*p*, *q*) *also*.

*Proof*. Condition on *q* = *q*_0_, and (for the moment) *F_n_*(*q*) = *m*. Set *ϵ* < *δ* and let

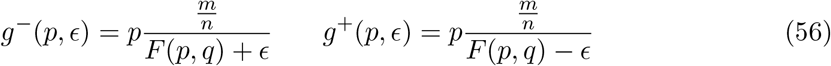

From the Dvoretzky-Kiefer-Wolfowitz (DKW) inequality we have

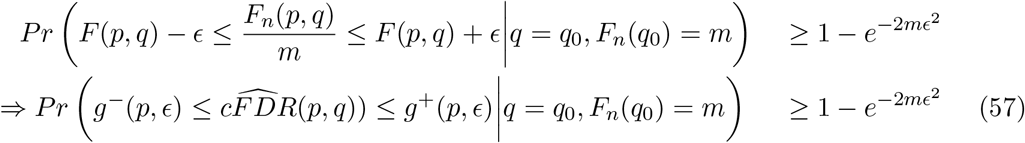

If *∂C*(*p*, *q*)/*∂p* ≥ 0, then (57) also holds for 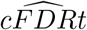. To see this, note 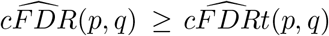, so if 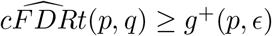 then 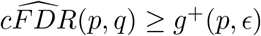 also. Now

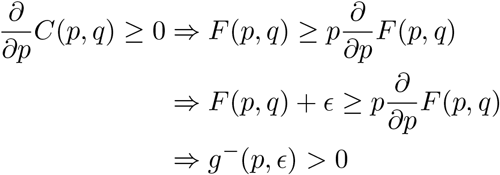

Suppose that for some *p* we had 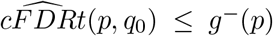. Then either 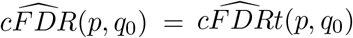 or 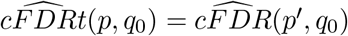 for some *p*′ > *p*. In the first case 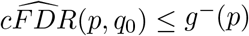, and in the second, 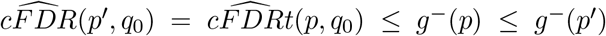; in either case, 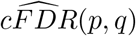 escapes the bound *g*^−^(*p*) somewhere. Thus the probability on the LHS of (57) can only increase if 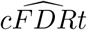 replaces 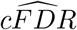, and 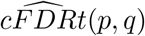 is contained within the bounds *g*^−^(*p*), *g*^+^(*p*) with probability at least 1 − exp(−2*mϵ*^2^).

We now move to remove the condition *F_n_*(*q*_0_) = *m*. Denote the events

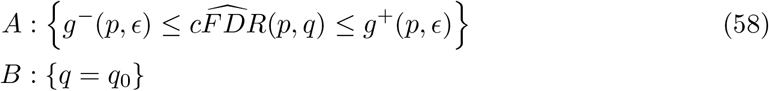

and, for some *ϵ*_2_ < *F*(*q*_0_)

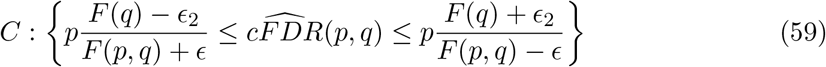

Denote by *S*(*ϵ*_2_) the set of integers in [*n*(*F*(*q*_0_) – *ϵ*_2_), *n*(*F*(*q*_0_) + *ϵ*_2_)] (and assume *n* is large enough that *S*(*ϵ*_2_) is nonempty). If *m* = *F_n_*(*q*_0_) ∈ *S*(*ϵ*_2_), the interval in event *A* is a subinterval of that in event *C*. Thus

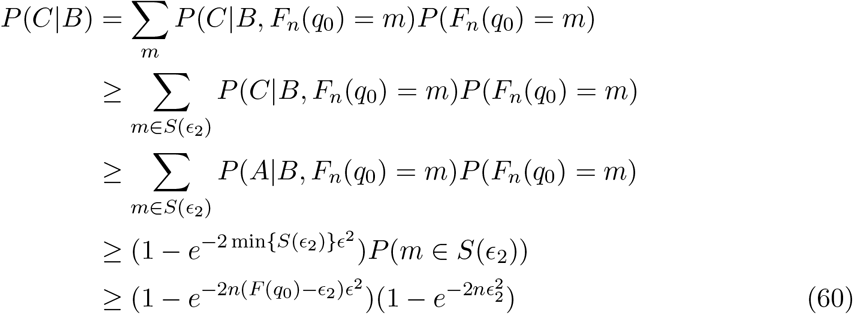

where the last inequality comes from the DKW inequality on *F_n_*(*q*). Since *p* ≥ *p_ϵ_*, and *F*(*p*, *q*) ≥ *γ* the widest part of the interval in event *C* can be made arbitrarily small on the interval (*p_ϵ_*, 1) and 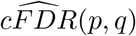 converges uniformly to *C*(*p*, *q*). If *∂C*(*p*, *q*)/*∂p* ≥ 0, then so does 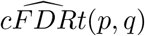.

#### Corollary 7.1.

*Under the assumptions in theorem 7*, 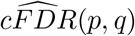 *and* 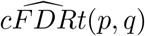 *are bound with fixed probability on the line segment q* = *q*_0_, *p_γ_* < *p* < 1 *in intervals of width O*(*n*^−1/2^)

*Proof*. In inequality 60, set

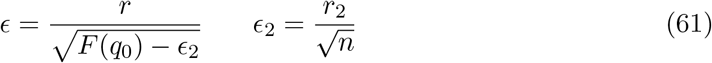

Then the RHS is 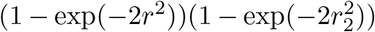 which may be made arbitrarily small by varying *r, r*_2_, and the difference between the upper and lower bounds in event *C|B* is

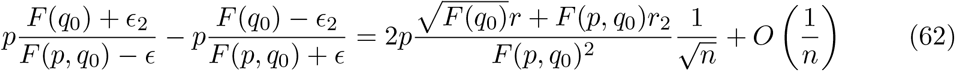

#### Theorem 8.

*Suppose that on a line segment q* = *q*_0_, *p_γ_* < *p* < 1, *we have F*(*p*, *q*) ≥ *γ* > 0, *F*(*q*_0_) > 0, *and ∂C*(*p*, *q*)/*∂p* ≥ *γ*_2_ > 0. *Denote by l*(*α*) *the value of p at the intersection of the L-curve L*(*α*) *with the line q* = *q*_0_, *so*

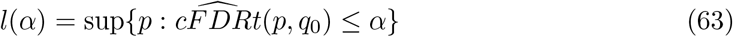

*and c*(*α*) *the value of p such that C*(*p*, *q*_0_) = *α* (*unique if it exists*). *For any δ* > 0, *the function* |*l*(*α*) – *c*(*α*)| *converges uniformly to* 0 *for α* ∈ [*C*(*p_ϵ_, q*_0_) + *δ*, 1].

*Proof*. Since *C*(*p*, *q*_0_) is continuous and increasing on [*p_ϵ_*, 1], the value *c*(*α*) exists for *α* ∈ [*C*(*p_ϵ_, q*_0_), *C*(1, *q*_0_)] ⊃ [*C*(*p_ϵ_, q*_0_) + *δ*, 1] by the intermediate value theorem. The function 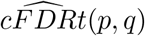 is continuous and nondecreasing on [0, 1] and hence *l*(*α*) exists for 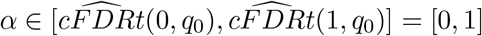.

Given arbitrarily small positive *ϵ*_3_, *δ*_2_ < *δ* choose *n* large enough that 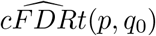 is contained in [*C*(*p*, *q*_0_) – *δ*_2_, *C*(*p*, *q*_0_) + *δ*_2_] for *p* ∈ [*p_ϵ_*, 1] with probability at least 1 – *ϵ*_3_. Then with probability ≥ 1 – *ϵ*_3_, whenever the curve 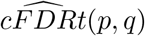 is in the region bounded by the rectangle *p_ϵ_* ≤ *p* ≤ 1, *C*(*p_ϵ_, q*_0_) + *δ* ≤ *q* ≤ 1, it is bounded by the curves *C*(*p*, *q*_0_) – *δ*_2_), *C*(*p*, *q*_0_) + *δ*_2_. The distance between the two curves in the *q*-direction is at most 2*γ*_2_*δ*_2_. Thus, if for some *α* ∈ [*C*(*p_ϵ_, q*_0_) + *δ*, 1], we have |*l*(*α*) – *c*(*α*)| > *γ*_2_*δ*_2_, the curve 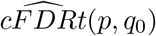 must escape the region bounded the curves *C*(*p*, *q*_0_) – *δ*_2_), *C*(*p*, *q*_0_) + *δ*_2_.

So with probability at least 1 – *ϵ*_3_ we have

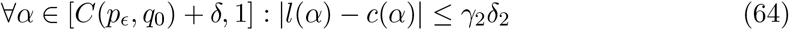

which proves the statement. This is illustrated in figure 9.

**Figure 9:**
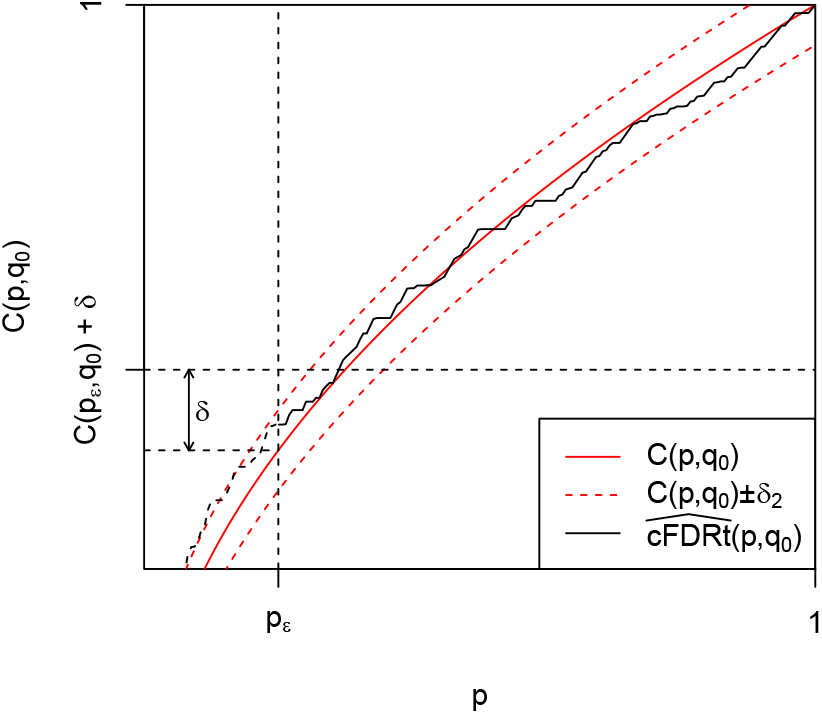
Convergence of intersections of L-curves with a line *q* = *q*_0_. Functions 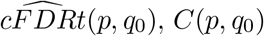, and *C*(*p*, *q*_0_) ± *δ*_2_ are shown. The vertical distance between dashed red lines is 2*δ*_2_, and since *∂C*(*p*, *q*_0_)/*∂p* ≥ *γ*_2_, the horizontal distance is at most 2*δ*_2_*γ*_2_. We must restrict the proof to *α* > *C*(*p*, *q*_0_) + *δ* because we cannot assert the behaviour of 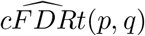 left of the line *p* = *p_ϵ_*.

We now proceed to the proof of theorem 2, restated here:

#### Theorem 2.

*Let R be the region of the unit square for which F*(*p*, *q*) ≥ *γ* > 0 *and F* (*q*) > 0. *Then on R*, 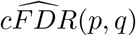 *converges uniformly to C* (*p*, *q*), *and if ∂C*(*p*, *q*)/*∂p* ≥ 0, *then so does* 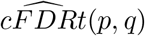

*Proof*. We proceed very similarly to theorem 7. We employ a result from [50] that for any *ϵ* > 0

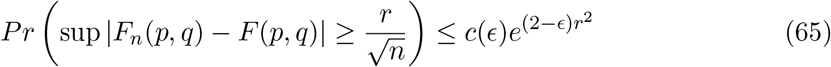

from which, wherever 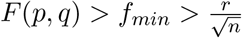

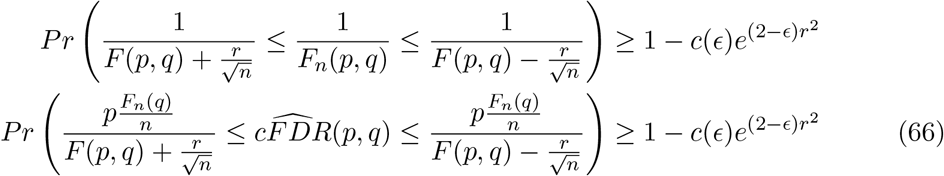

The values *F_n_*(*p*, *q*) and *F_n_*(*q*) are dependent. However, given some *r*_2_ > 0, we have for all *q* (by the DKW inequality)

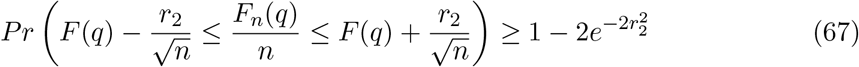

Denoting the event in (66) by *A*, the event in (67) by *B*, and *C* as:

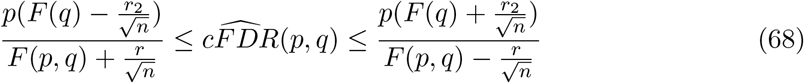

we have, since the interval in *A* is a subinterval of that in *C* when conditioning on *B*:

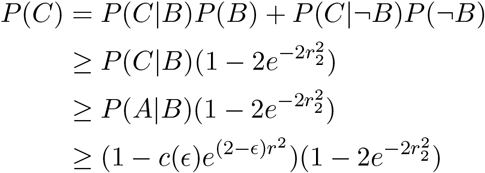

As before, this bound also holds for 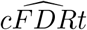 as long as *∂C*(*p*, *q*)/*∂p* > 0.

#### Corollary 8.1.

*Under the assumptions in 2*, 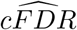 *and* 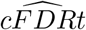 *are bound with fixed probability in R in intervals of width O*(*n*^−1/2^)

*Proof*. The difference between the upper and lower bounds in (68) is

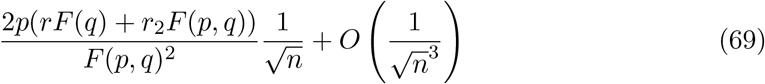

Our final result describes errors on v-values. Given an L-region *L*(*α*), we define the M-region as the ‘expected’ L-region:

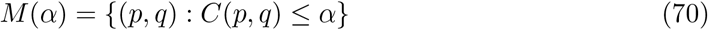

and the ‘error’ on the v-value 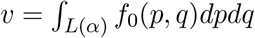 as

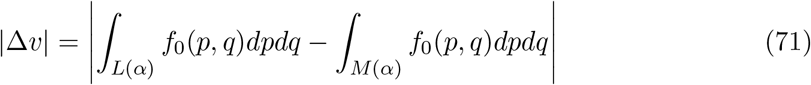

We now are now in a position to prove theorem 3.

#### Theorem 3.

*Define R as in theorem 2, and further assume that* 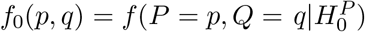 *is known and on R we have ∂C*(*p*, *q*)/*∂p* ≥ *γ*_2_. *Write R^c^* = [0, 1]^2^ \ *R. Then the maximum error on any v-value is*

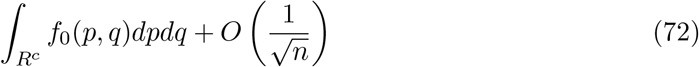

*Proof*. Using theorem 2, bound 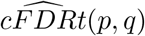 between *C*(*p*, *q*)–*δ, C*(*p*, *q*)+*δ* with probability ≥ 1 – *ϵ*_3_, where 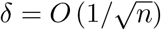.

Since *F*(*p*, *q*) is nondecreasing with p, we can describe *R* = {(*p*, *q*) : *F*(*p*, *q*) ≥ *γ*} as the union of line segments *q* = *q*_0_, *p_ϵ_*(*q*_0_) ≤ *p* ≤ 1. We now define *R*_1_ as the union of all line segments *q* = *q*_0_, *p_ϵ_*(*q*_0_) + *δγ_2_* ≤ *p* ≤ 1; that is, *R* with the leftmost border shifted *δγ_2_* to the right.

We show the result by firstly noting that if an L-curve intersects a line segment *q* = *q*_0_ at *l*(*α*) > *p_ϵ_*(*q*_0_) + *δγ_2_*, and we have that |*l*(*α*) – *c*(*α*)| > *δγ_2_* (where *c*(*α*) is the intersection of the border of *M*(*α*) with *q* = *q*_0_), then event *C* (equation 68) must have occurred in *R*, by the same argument as for theorem 8. Thus with probability at least 1 – *ϵ*_3_, every segment of a right-most border of an L-region *L*(*α*) in *R*_1_ is at a horizontal distance from the corresponding rightmost-border of *M*(*α*) of at most *δγ_2_*

We now write

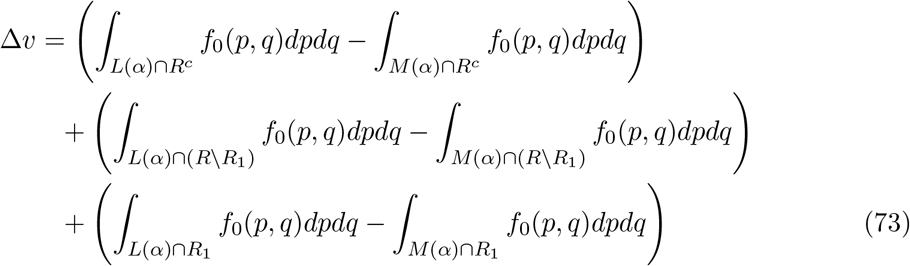

The first term is at most *∫_R^c^_ f*_0_(*p*, *q*)*dpdq*. The region *R* \ *R*_1_ has constant width *δγ*_2_, and since *f*_0_ only varies with *q*, hence the second term is at most *∫*_*R\R*_1__ *f*_0_(*p*, *q*)*dpdq* = *δγ*_2_ = *O*(*n*^−1/2^). Within *R*_1_, if the horizontal separation between curves at the rightmost border of *L*(*α*) and *M*(*α*) is greater than *δγ*_2_, then C has occurred, so this can happen with probability at most *ϵ*_3_. Thus with probability 1 − *ϵ*_3_, the third term is also bounded by *δγ*_2_ = *O*(*n*^−1/2^), establishing the result.

### 8.4 Influence of a single point

Intuitively, adding a single point to a map defined by *n* other points should have a small effect on that map, and hence on the resultant v-values. We show the following:

#### Theorem 4.

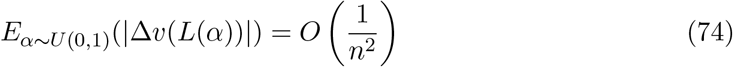

*Proof*. Consider the profile of 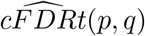 on a line *q* = *q*_0_, and how this changes with the addition of (*p*, q\**). The functions *F_n_*(*q*), *F_n_*(*p*, *q*) will be taken to be with respect to the *n* points (*p_i_, q_i_*) but not (*p*, q\**).

For *q*_0_ < *q\**, the addition of (*p*, q\**) changes neither *F_n_*(*q_0_*) nor *F_n_*(*p, q_0_*), so on lines *q* = *q_0_* < *q\** the profile of 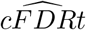 will remain the same.

Denote

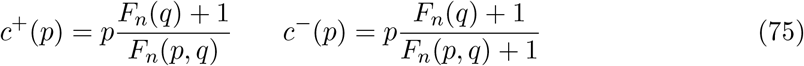

For *q_0_* > *q*, p < p\**, the value of 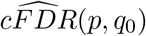 will increase by

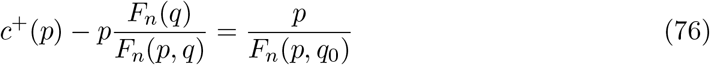

and for *q*_0_ > *q\**, *p* > *p\**, it will decrease by

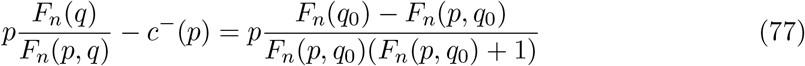

In either case, 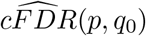 changes by 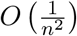. The behaviour of 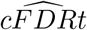 is a little more complex. If we define 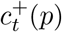 and 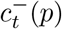 analogously to 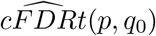, then for *p* > *p\**, 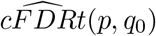 shifts to 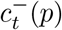, and for *p* < *p\**, it shifts to 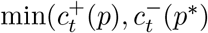 (see example in figure 10).

We can show that the absolute difference in 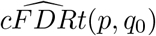 is always less than the absolute difference in 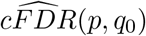 after adding (*p*, q\**). Denote these differences 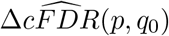 and 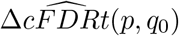. Since 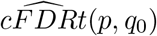 always shifts to between 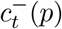 and 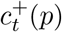, it suffices to show that

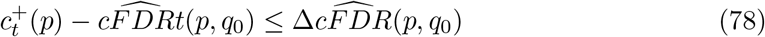

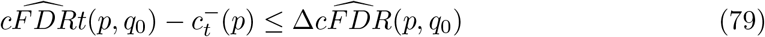

Inequality (78) follows from the observation that 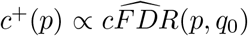, so order relations between 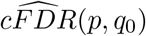 and *c^+^*(*p*, *q*_0_) are preserved. Thus

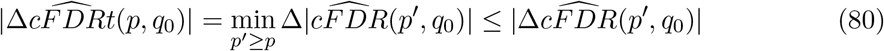

Order relations are not preserved between 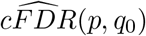 and *c*^−^(*p*), but the denominators increment at the same values of *p*. The functions 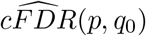 and *c*^−^(*p*) both rise linearly in *p* between successive increment points *p_a_, p_d_* of *F_n_*(*p*, *q*_0_), with 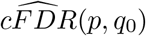 having the higher gradient, since

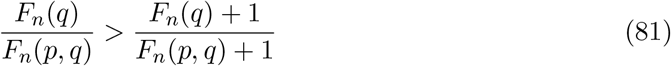

At *p_d_*, both functions are discontinuous and drop in value. On (*p_a_, p_d_*), the values of 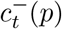 and 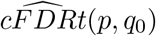 are either equal to *c*^−^(*p*), 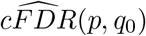, or ‘censored’ at some values *c*^−^(*p′*), 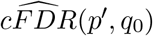 with *p′* > *p* (see the right-hand part of figure 10 for an example of this). We note that 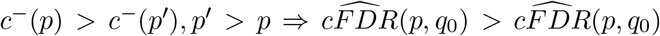, so the first point at which 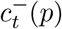 is censored on (*p*_1_, *p*_2_) is further right than the first point at which 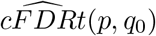 is censored. Denote the leftmost point at which 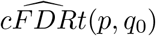 is censored as *p_b_* and the leftmost point at which 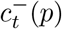 is censored as *p_c_*, so *p_a_ ≤ p_b_ ≤ p_c_ ≤ p_d_*.

On (*p_a_, p_b_*), where neither are censored, 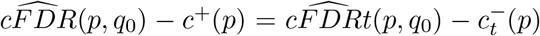 and 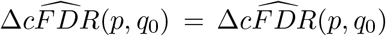. On (*p_b_, p_c_*), when only 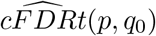 is censored, 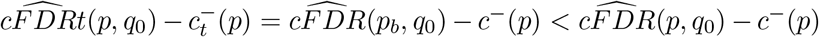, so 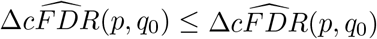. On (*p_c_, p_d_*), we have 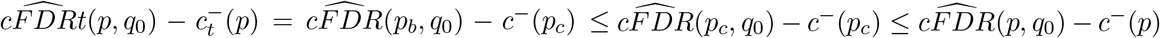, so again, 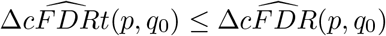. Thus, for all *p*,

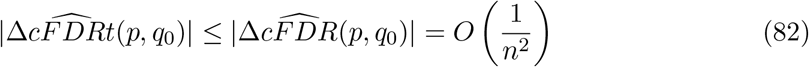

where the multiplicative factor in *O*(1/*n*^2^) is independent of *q*_0_. This inequality is demonstrated in the right panel of figure 10.

Denote by *l_*α*_* the value of *p* at the intersection of an L-curve corresponding to 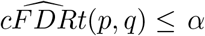 with the line *q* = *q*_0_. We have 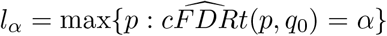. The value *l_α_* may shift substantially when adding *p*, q\**, as shown in figure 10 However, the effect is small on average. The plot of the function *l*(*α*) before and after adding (*p*\*, *q*\*) is identical to the plot of the function of *p* given by 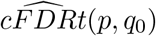 before and after adding (*p*\*, *q*\*) rotated by *π*/2. The average difference in movement of *l_α_* is the integral of the difference in *l_α_* with and without (*p*\**, q*\*). However, this is simply the area between the two curves, which is invariant under rotating *π*/2. Hence

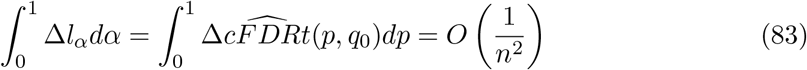

Denote the region 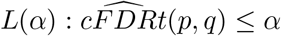 and the co-ordinates of its rightmost border (L-curve) (*q*, *l_α_*(*q*)), *q* ∈ (0, 1). Then, denoting the indicator function by *I*

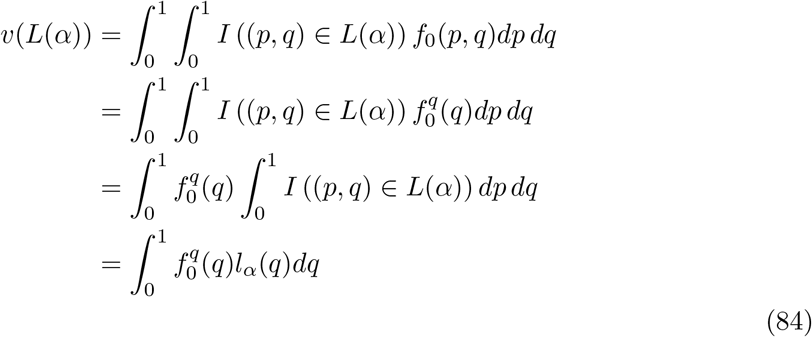

and the average error in v-values *v*(*L*(*α*)) over *α* ~ *U*(0, 1) is

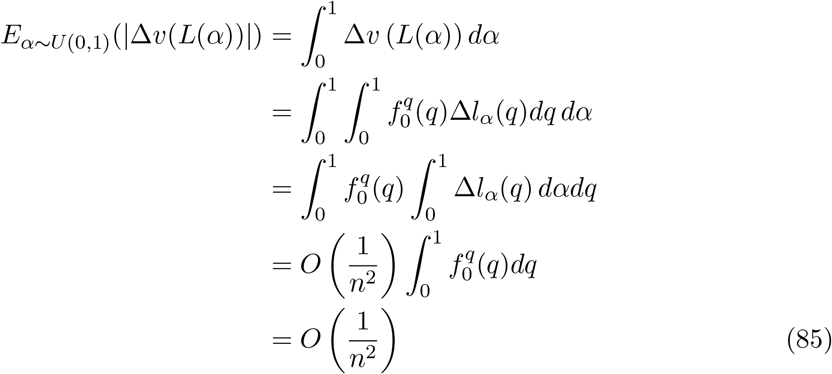

as required.

**Figure 10:**
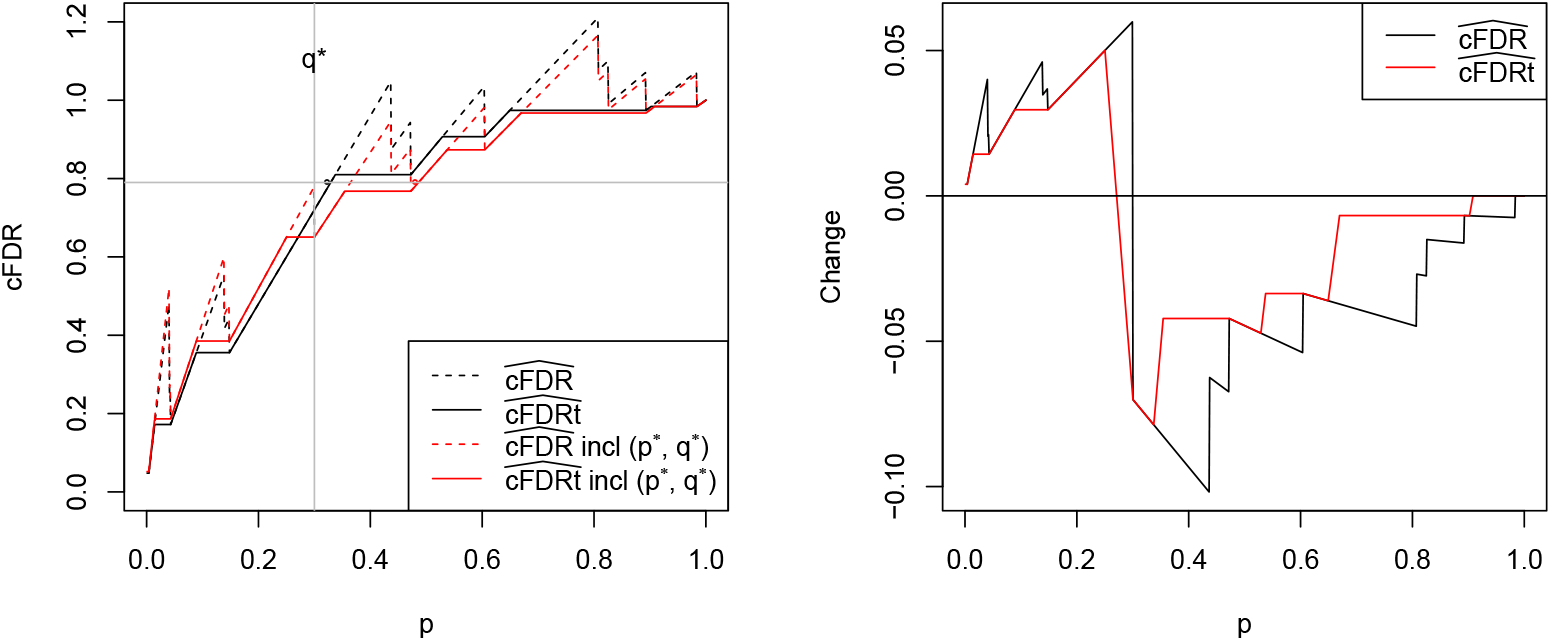
Behaviour of 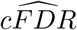 and 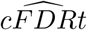 on a line *q* = *q*_0_ after adding a point (*p*\*, *q*\*) to a set of *n* points (*n* is considerably smaller in this example than in figure 9). In the left panel, curves of 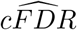 and 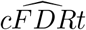 before and after adding (*p*\*, *q*\*) are shown. Adding (*p*\*, *q*\*) may have a substantial impact on the intersection of an L-curve with *q* = *q*_0_, such as that in the horizontal line: the black and red points show the intersection points of a curve before and after adding (*p*\*, *q*\*). However, the average effect across all curves is limited to the integral of the difference between the black and red lines, which is *O*(1/*n*^2^). The right panel demonstrates that 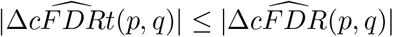.

### 8.5 Asymptotic equivalence of PDF- and CDF- based L-regions

We show in this section that under a fairly common condition L-regions based on the PDF of *p*, *q* are similar to L-regions based on the CDF. In this section, we generally work on the Z-scale rather than the p-value scale for convenience.

Denote a ‘fast-decreasing’ function as a function *g* such that for each *ϵ*_1_, *ϵ*_2_ > 0, there exists *δ* such that for all *X, Y* of distance at least *δ* from the origin, we have

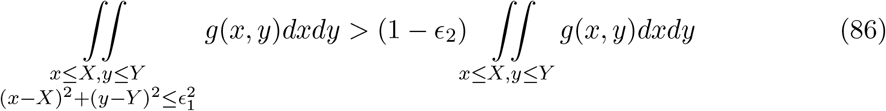

so for *x* ≤ *X, y* ≤ *Y*, the function *g* falls off rapidly enough as *x, y* decrease that we can disregard its value except when it is close to *X, Y*.

We show the following:

#### Theorem 9.

*Denote c*(*x, y*) = *f*_0_(*x, y*)/*f*(*x, y*) *and C*(*x, y*) = *F*_0_(*x, y*)/*F*(*x, y*). *Given a region of the* (−, −) *quadrant A_ϵ_*, = (−∞, 0] × (*I*_1_ – *ϵ*, *I*_2_ + *ϵ*) (*where ϵ* > 0 *is arbitrarily small), suppose that for x, y* ∈ *A_ϵ_ and for sufficiently small *α* we have*

1. *f*_0_ *and f are fast-decreasing continuous positive functions*
2. *Along horizontal rays in A, c*(*x, y*) *satisfies ∂*^2^ log (*c*(*x, y*)) */∂x*^2^ > 0
3. *The contour c*(*x, y*) = *α is continuous and bounded, and the rightmost bound increases to* ∞ *as α* → 0

*Then for each ϵ*_3_ > 0, *there exists an ϵ*_1_ *as above and an α*_1_ *such that whenever α* < *α*_1_, *there is a contour of C*(*x, y*) *is never further than ϵ*_3_ *from the contour c*(*x, y*) = *α in the region A*_0_.

*Proof*. Set *R*_3_ as the region defined by the union of all circles of radius *ϵ*_3_ with centres on points *y*, *l_α_*(*y*). Choose *ϵ*_1_ = *ϵ*_3_/2 (supposing that *ϵ*_1_ < *ϵ*), and define *R*_1_ similarly to *R*_3_ with radii *ϵ*_1_. Let *α*^+^ be the minimum value of *f*_0_/*f* on the rightmost border of *R*_1_, and *α*^−^ the maximum value on the leftmost border so *α*^+^ > *α* > *α*^−^.

Condition 2 implies that for fixed *y*

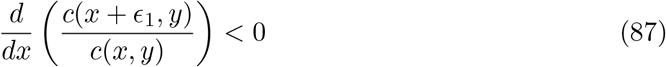

Since the horizontal distance between the rightmost border of *R*_3_ and the curve is at least 2*ϵ*_1_ and similarly from the leftmost border of *R*_3_, the values *α*^+^ – *α*, *α* – *α*^−^ must increase for fixed *ϵ*_1_ as we move left. Thus, for some fixed *ϵ*_2_ > 0, choose *δ*_2_ large enough that *α*^+^/*α*^−^ > 1/(1 – *ϵ*_2_)^2^ and larger than the *δ* corresponding to *ϵ*_1_, *ϵ*_2_ by assumption, and *α*_1_ large enough that the contour *c*(*x, y*) is entirely left of the line *x* = −*δ*_2_.

Let *X, Y* be a point in *A*_0_ to the right of *R*_3_, so a circle of radius *ϵ*_1_ centred at *X, Y* is in *A_ϵ_* but does not intersect *R*_1_. Thus across such a circle, the value of *c*(*x, y*) is at least *α*^+^. Similarly, across a circle of radius *ϵ*_1_ centred to the left of *R*_3_, the value of *c*(*x, y*) is at most *α*^−^

For *x, y* to the right of *R*_3_, denote by *H* the circle of radius *ϵ*_1_ centred at *x, y*. Now by the fast-decreasing property of *f*_0_ and *f*, we have

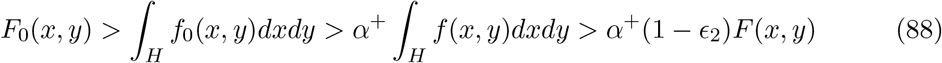

so *C*(*x, y*) > *α*^+^(1 – *ϵ*_2_). Similarly for *x, y* to the left of *R*_3_, we have *C*(*x, y*) < *α*^−^/(1 – *ϵ*_2_). By our choice of *α*_1_, we have *α*^+^(1 – *ϵ*_2_) > *α*^−^/(1 – *ϵ*_2_), so any contour of *C*(*x, y*) at a level between these values must pass within *R*_3_ through *A*_0_.

Contours of *F*_0_/*F* correspond to contours of *cFDR*, and contours of *f*_0_*/f* correspond to contours of 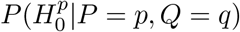. Theorem 8.5 has obvious analogies in other quadrants, and for the p-value rather than z-score scale.

The conditions in the theorem may seem restrictive, but they are satisfied by many distributions; for instance, when *f*_0_ and *f* are mixture Gaussian, and *f* dominates *f*_0_ as |*x*| → ∞. Figure 11 shows the similarity of a range of shapes of contours of *C* and *c*.

**Figure 11:**
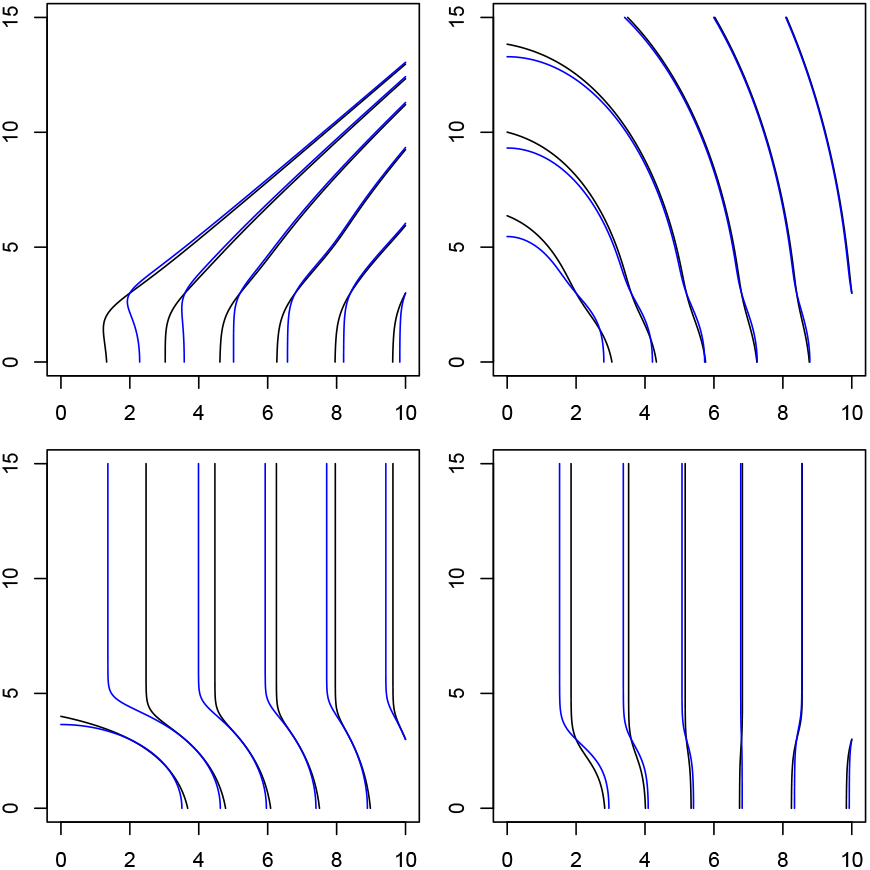
These plots show contours of CDF- based and PDF-based L-regions for a range of distributions of *P*, *Q*. Plots are on the Z-score scale (eg, rejection regions in terms of *Z_P_, Z_Q_*). The distributions are parametrised in terms of the mixture-Gaussian distribution detailed in supplementary material, section 9.4.1; parameters ((*π*_0_, *π*_1_, *π*_2_, *τ*_1_, *τ*_2_, *σ*_1_, *σ*_2_)) were (0.7,0.1,0.1,2,3,1.5,1.5), (0.7,0.1,0.1,2,2,3,3), (0.99,0.0005,0.0003,2,3,4,3), (0.7,0.1,0.05,3,2,2,2) respectively. Curves are generated passing through the points *Z_P_*, *Z_Q_* = (3,2..6), with curves further to the right corresponding to smaller *α*. As *α* gets smaller, contours *F*_0_(*x, y*)/*F*(*x, y*) = *α* (black lines) become closer to contours of *f*_0_(*x, y*)/*f*(*x, y*) = *α* (blue lines) under reasonably general circumstances

**Figure 12:**
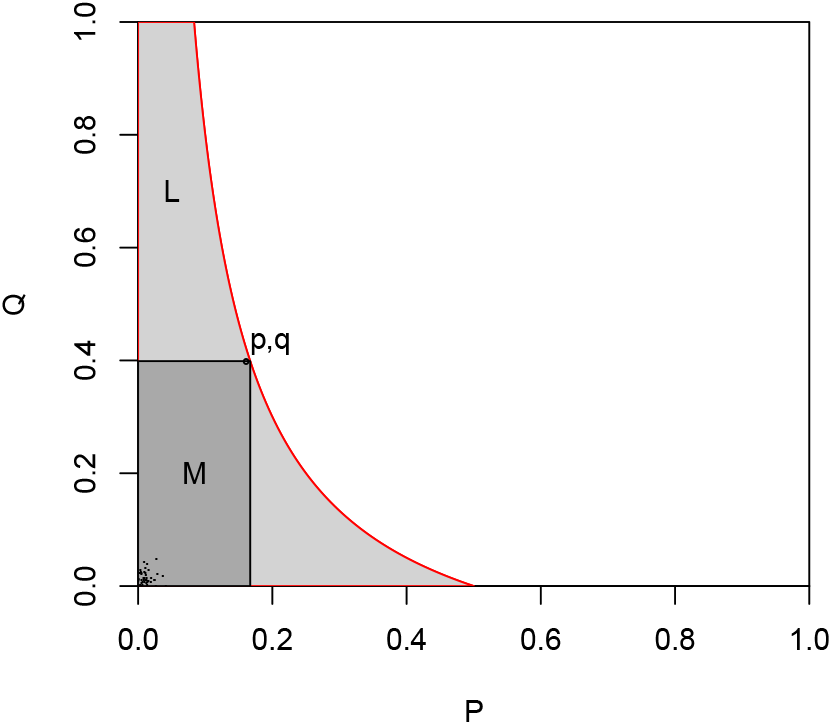
Rejection regions for rejection procedure (9) under assumptions in section 8.2. FDR is not controlled for either of the cFDR-based rejection regions. For reference, the B-H procedure applied to the set of (*p*, *q*) pairs with *q* ≤ *q*_0_ would reject everything in the dark gray rectangle, which includes all true positives, and this would control FDR at *α*. The cFDR based rejection regions reject the same number of true-positives, but far more false positives, so FDR control is lost.

## 9 Supplementary material

### 9.1 TWAS details

TWAS test association of gene expression (a biologically interpretable quantity) with some trait, when the trait and expression have not been measured on the same individuals, by using a reference expression-quantitative trait locus (eQTL) study and genome-wide association study (GWAS). A TWAS firstly uses eQTL data to learn rules to predict mRNA expression levels in a given tissue according to individual genotype, then applies these rules to predict expression for each individual in a GWAS, and finally compares these predicted expression levels across the GWAS trait of interest [40].

The results from a TWAS are a set of p-values corresponding to tissue-gene pairs. The object is to find which tissue-gene pairs are associated with the disease under consideration; that is, which p-values come from a distribution other than *U*(0, 1). In general, such tissuegene pairs are a small proportion of all those considered. We consciously ignore any prior information which could be derived from tissues likely to be BRCA or OCA associated, for purposes of demonstration.

We considered TWAS datasets for breast cancer (BRCA, [41]) and ovarian cancer (OCA, [42]), containing tests for varying numbers of genes across 54 tissues. BRCA and OCA have considerable phenotypic overlap [43], and we may hope that summary statistics for one disease may be useful for leverage in association analyses of the other. We considered RNA-tissue pairs available in both datasets, restricting our analysis only to pairs in which RNA expression was predicted using data from the GTEx consortium, comprising a total of *n* = 80222 hypotheses.

Given the GWAS-scale dimensionality of testing, we chose a conservative FDR control level *α* = 1 × 10^−6^. We used both 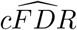 and 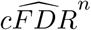 to generate v-values, and used the block-out method with blocks assigned according to genes, so expression levels for each gene were assigned a separate block (for 11327 folds in total).

### 9.2 Correlation of *P*, 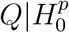

Throughout this paper we have largely assumed that 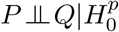, but methods can be easily adapted as long as the distribution 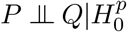 is known (or we are happy to assume it is known). An example where this occurs is in the GWAS literature in which the studies giving rise to *p_i_* and *q_i_* share samples (usually control samples), which induces a known correlation between *P* and *Q* under 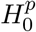.

This can generally be managed by using the true *f*_0_ (or an approximation allowing for dependence of *P*, *Q* under 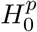) in equation 19 in the main paper and accounting for the true *f*_0_ in the computation of 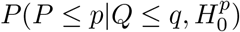 necessary to estimate cFDR (equation 6 in main paper) where it is otherwise equal to *p* under assumption (1) in the main paper. We demonstrate how to approximate 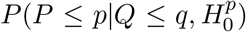 and *f*_0_ in the specific case of shared controls in a previous paper [23].

### 9.3 Simulations

### 9.4 Alternative cFDR estimators, and estimators of cfdr

In this section, we introduce new estimators of the cFDR 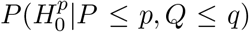 and cfdr 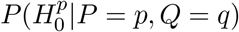, for use in simulations as detailed in section 5 of the main paper.

The main incentive for different estimators of cFDR is the tendency for the ECDF based estimator (equation (6) in the main paper) to have marked discontinuities at extremes of the unit square. This is illustrated in figure 13. The main incentive for estimators of cfdr is to allow comparison of PDF- and CDF- based estimators of the optimum rejection region detailed in section 2.2 in the main paper.

**Figure 13:**
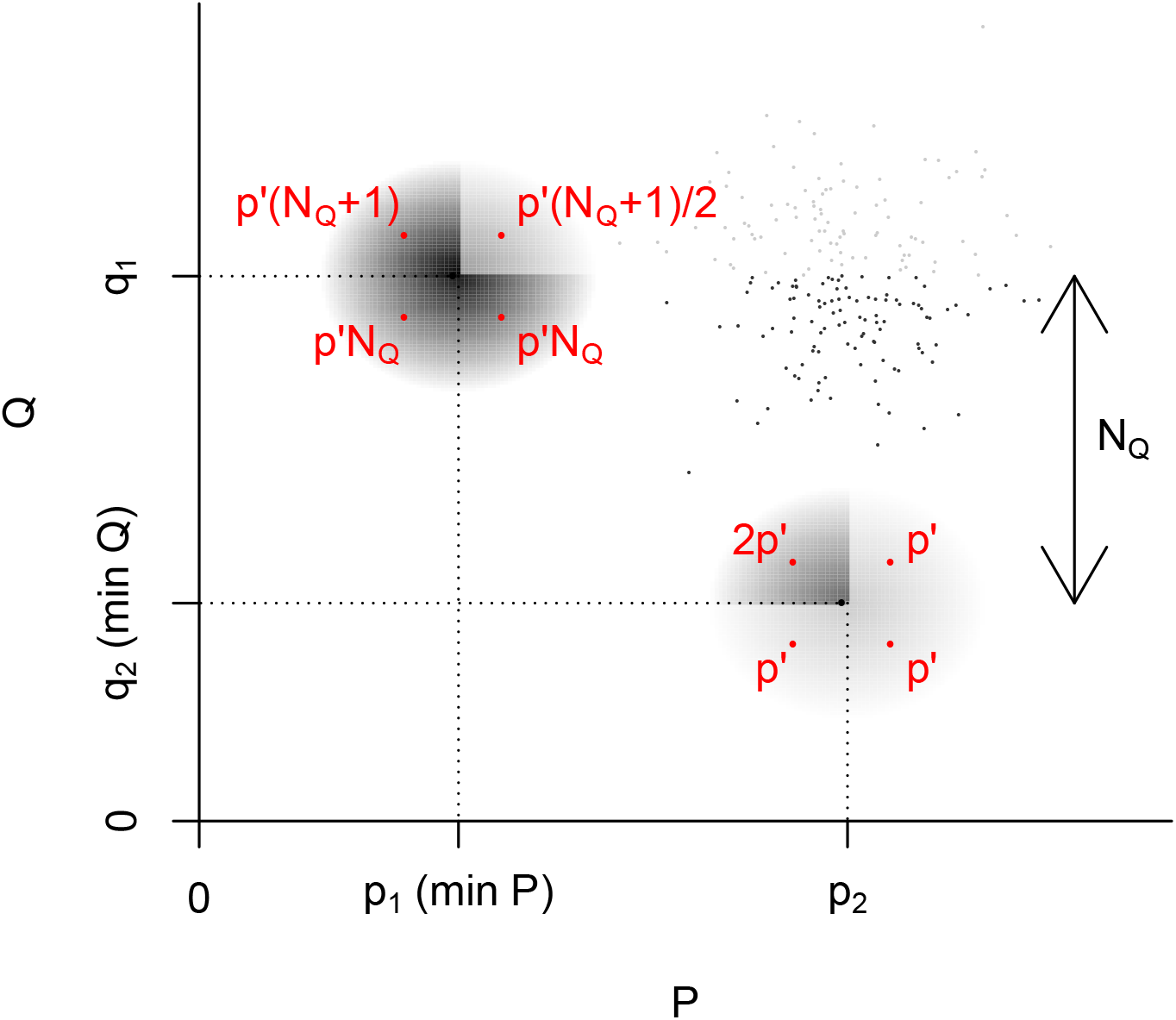
Dependence of 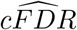 values on location of nearby points. In this example, we denote by (*p*_1_, *q*_1_), (*p*_2_, *q*_2_) the points at the (unique) left and lower extremes of the observed p-value distribution respectively; that is, *p*_1_ = min(*p_i_*), *q*_2_ = min(*q_i_*). We set *N_Q_* as the number of points with *q*_1_ ≤ *q_i_* ≤ *q*_2_ (small black points). If we add a test point (*p′,q′*) (shown in red) in a small neighbourhood of either (*p*_1_, *q*_1_) or (*p*_2_, *q*_2_), the estimated 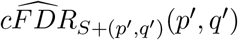 (shown in red next to the point) differs by a factor of 2 in different quadrants of the neighbourhood.

#### 9.4.1 Parametric estimators for cFDR and cfdr

The estimate 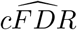 is based on empirical quantities estimated directly by empirical CDFs of (*P*, *Q*). We consider here estimators based on approximating the joint distribution of *P*, *Q* using a bivariate mixture-normal parametrisation. This estimator enforces continuity of 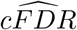 on the open unit square, and is robust to small deviations in p-values, overcoming the effect detailed in figure 13. It is easiest to visualise parametrisations as distributions over the unsigned Z scores (*Z_p_*, *Z_q_*) = (−Φ^−1^ (*P*/2), −Φ^−1^(*Q*/2)) with Φ^−1^ (*x*) denoting the standard normal quantile function at *x*.

We use a parametrisation with seven parameters: (*π*_0_, *π*_1_, *π*_2_, *τ*_1_, *τ*_2_, *σ*_1_, *σ*_2_), which parametrise a four-part bivariate mixture-Gaussian distribution over the (+, +) quadrant with PDF:

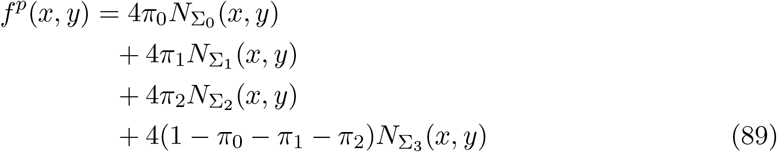

where *N*_∑_(*x, y*) is the PDF of the bivariate normal distribution centred at the origin with variance ∑, the factor of 4 is due to to only unsigned *Z*-scores being used, and

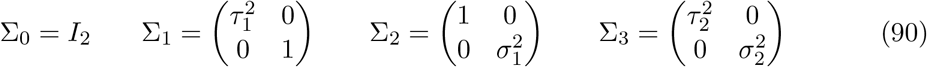

This model specifies a proportion *π*_1_ of study variables to be associated only with the trait of interest *P* (with *SD*(*Z_P_*) = *τ*_1_), a proportion *π*_2_ to be associated only with the second trait *Q* (with *SD*(*Z_Q_*) = *σ*_1_), and a proportion (1 – *π*_0_ – *π*_1_ – *π*_2_) to be associated with both (*Var*(*Z_P_*, *Z_Q_*) = Σ_3_). We can now write:

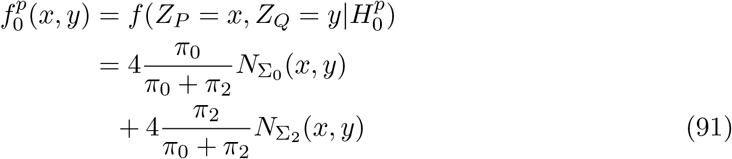

We allow different values of *σ*_1_, *σ*_2_ and *τ*_1_, *τ*_2_ to allow for potentially different reasons for shared (both *P* and *Q*) and independent (*P* XOR *Q*) associations.

Maximum-likelihood estimates of parameters (*π*_0_, *π*_1_, *π*_2_, *τ*_1_, *τ*_2_, *σ*_1_, *σ*_2_) can be obtained using an E-M algorithm [46]. Given these and corresponding estimates 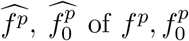 and 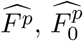 of *F^p^*, 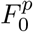 we can then define an estimate of cFDR (implicitly conditioning on parametric assumptions):

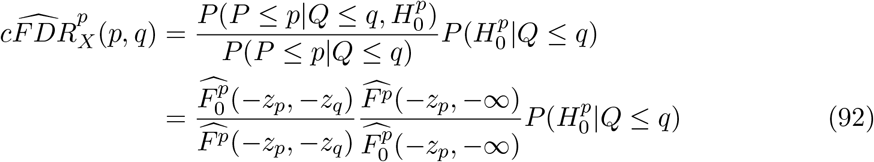

The quantity 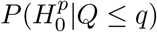 may be estimated up to directly from parametric assumptions as

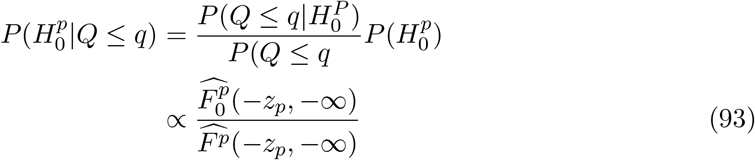

or may be estimated on the basis of the empirical CDF of *Q*|*P* > 1/2 as per equation 7 in the main paper. We found that the performance of 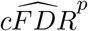 was stronger when using the empirical estimate in equation 7 than the parametric estimate in equation 93 (figure 14)

**Figure 14:**
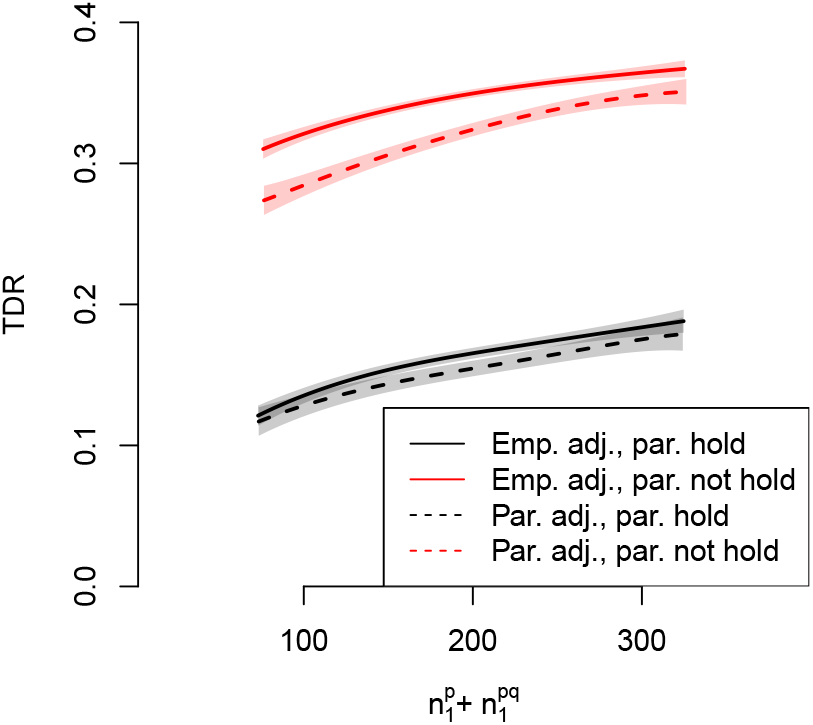
Performance (TDR) of parametric cFDR estimator against 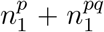, using either a parametric (‘Par. adj’) or empirical (‘Emp. adj’) estimate of 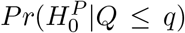, and separating cases where parametric assumptions hold (‘par. hold’) or do not hold (‘par. not hold’). See section 5 for further details. Shaded regions show 95% pointwise confidence envelopes. The empirical estimate leads to better performance when parametric assumptions are not satisfied, and equivocal performance when they are.

The local cfdr can be readily estimated as

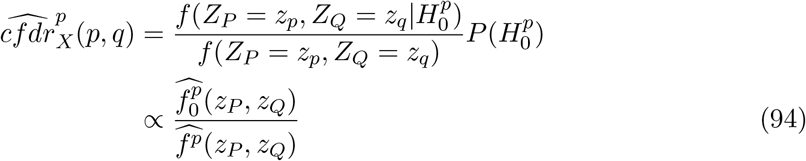

where *X* in this case is the set of points used in the estimation of parameters.

We note that estimator 92 implicitly includes an estimate of 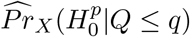, computed from parameter estimates. However, we found that performance of the estimator was improved when equation 7

#### 9.4.2 KDE-based estimators for cFDR and cfdr

To avoid distributional assumptions while maintaining a smooth form for the density of *p*, *q*, a second estimator of *P*(*P* ≤ *p*|*Q* ≤ *q*) can be derived from a two-dimensional kernel density estimator (KDE). We had no reason to prefer any kernel function over another, so opted to use a normal kernel with constant variance *I*_2_. The PDF corresponding to *Z_p_*, *Z_q_* at *x, y* was modelled in the usual way as

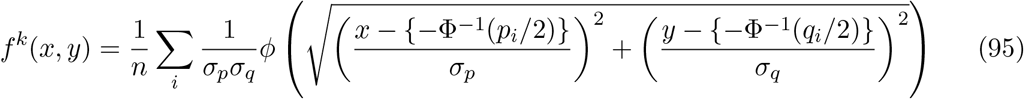

where *ϕ*(.) is the standard normal density. Values *σ_p_* and *σ_q_* are determined using a standard method based on the observations *p_i_*, *q_i_* ∈ *X* [47].

Unlike the parametric estimate above, this does not intrinsically specify the density of *P*, 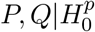. We thus incorporate the estimator 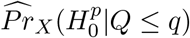 from equation 7 in the main paper, and write (implicitly conditioning on correctness of approximations)

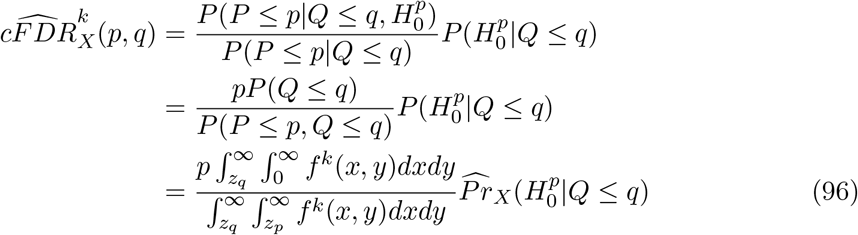

where *X* is the set of points used in the KDE in equation 95. We note that this estimator converges to 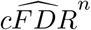 as *σ_p_*, *σ_q_* → 0.

Estimating local cfdr using KDEs requires estimation of 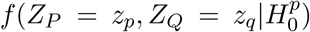. We use assumption 1 from the main paper, and as for equation 7 in the main paper we assume that 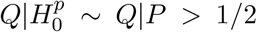. We then fit a one-dimensional KDE to the values *z_q_i__*|*p_i_* > 1/2, and denote the resultant function of q as 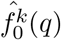. We then write (conditioning on assumptions)

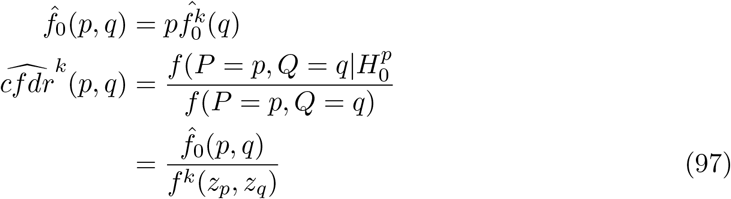

### 9.5 Analysis of cFDR estimators

We required that all estimators be nonincreasing in *p*, so all were censored when generating L-curves or designing rejection procedures in the same way as in 9 and 16 for 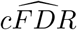 in the main paper.

We show in figures 15, 16, 17 a series of plots at different values of *n* which indicate the behaviour of L-curves as *n* increases. In all cases, *P*, *Q* are sampled under the parametric assumptions in supplementary section 9.4.1, with (*π*_0_, *π*_1_, *π*_2_, *τ*_1_, *τ*_2_, *σ*_1_, *σ*_2_) = (0.7, 0.1, 0.15,1.5, 2,1.5, 2). Curves are drawn through (0.1, 0.1), which would generally corresponds to a very high FDR level, so oracle PDF and oracle CDF curves are markedly different.

**Figure 15:**
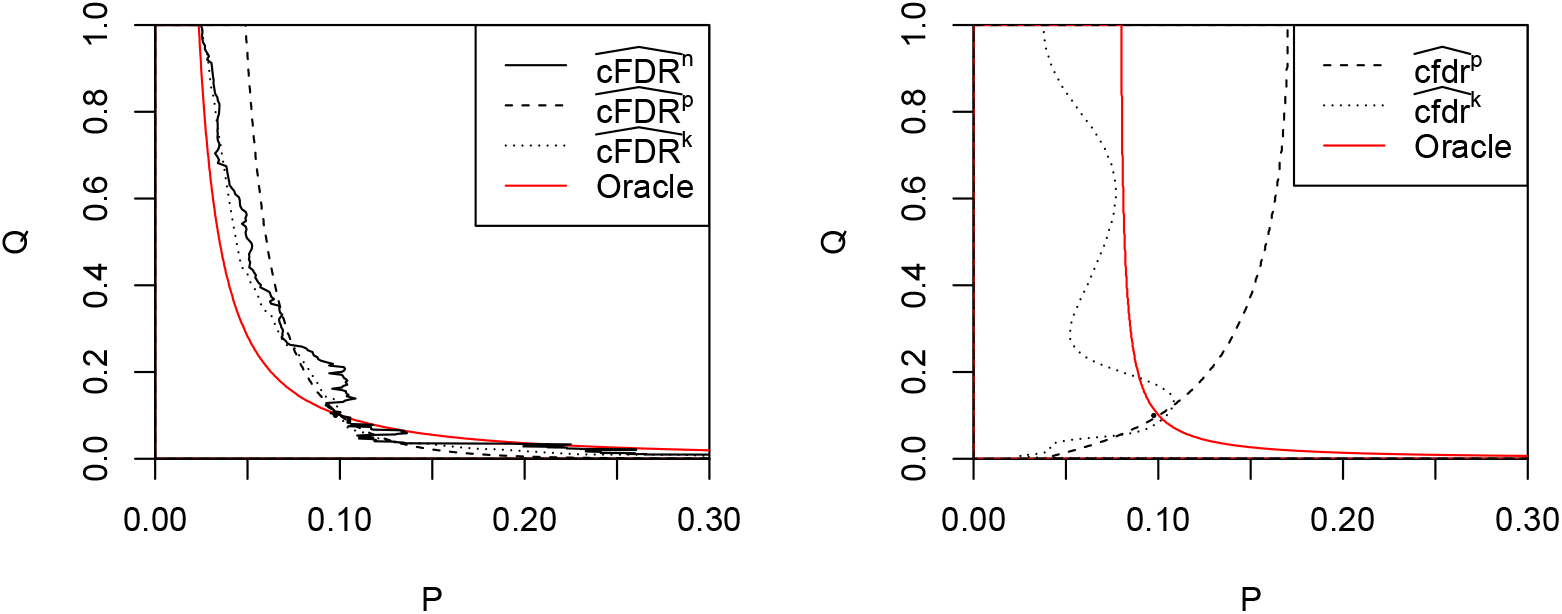
L-curves using various methods for cFDR estimation; n=1000. CDFs on left, PDFs on right.

**Figure 16:**
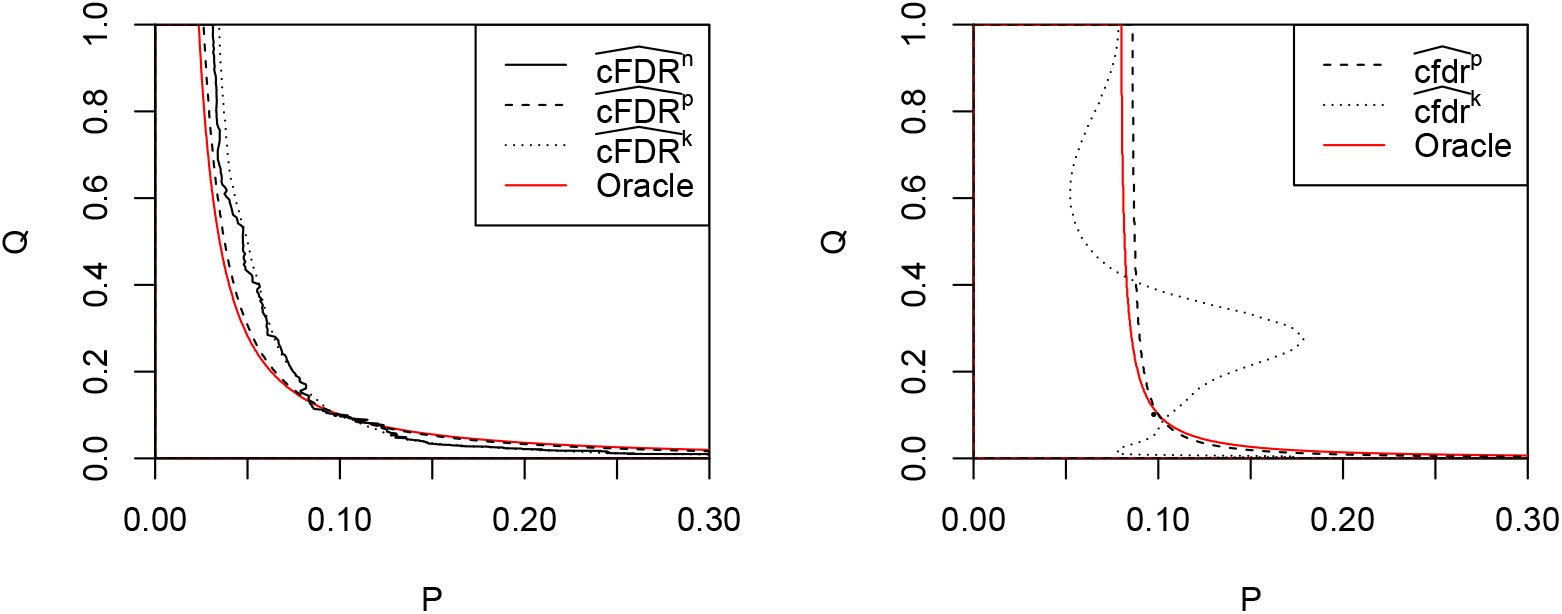
L-curves using various methods for cFDR estimation; n=10000. CDFs on left, PDFs on right

**Figure 17:**
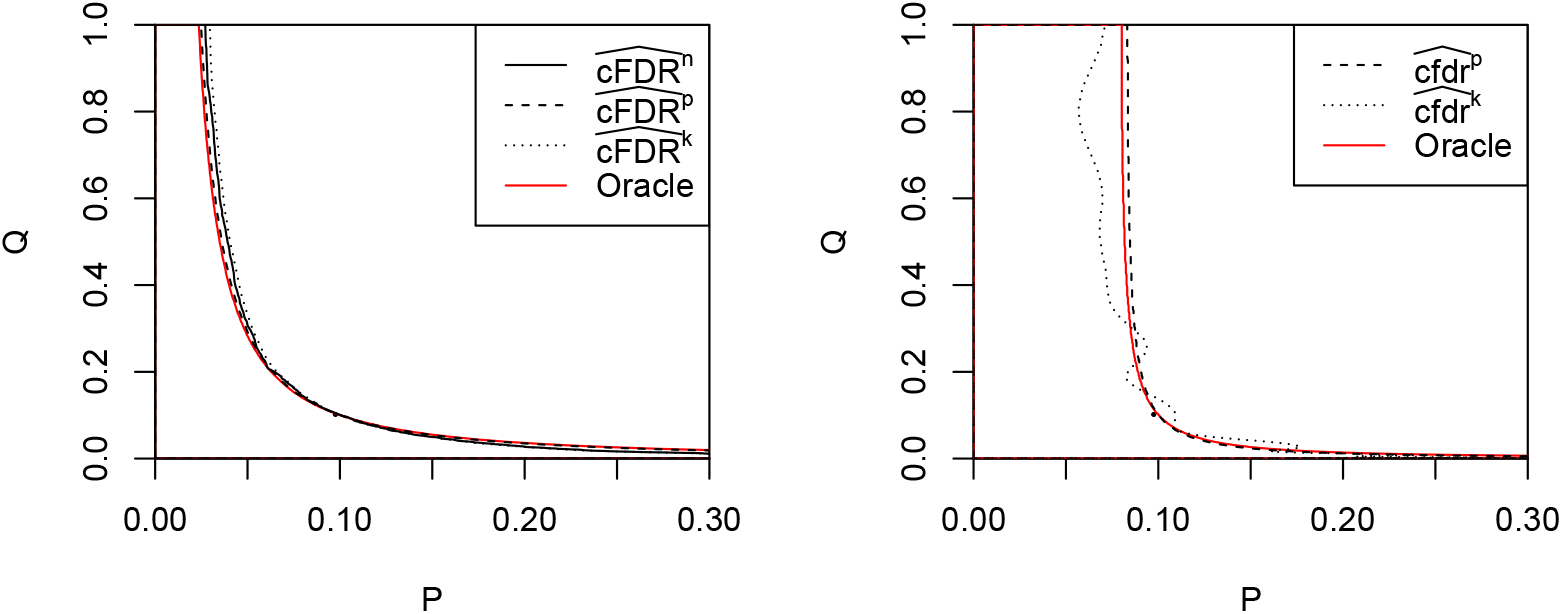
L-curves using various methods for cFDR estimation; n=100000. CDFs on left, PDFs on right.

Importantly, 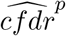 and 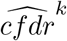 converge to the optimal rejection region (oracle PDF) while 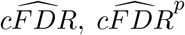 and 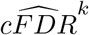 do not. However, the estimates of the latter are less noisy.

## 10 Supplementary figures

**Figure 18:**
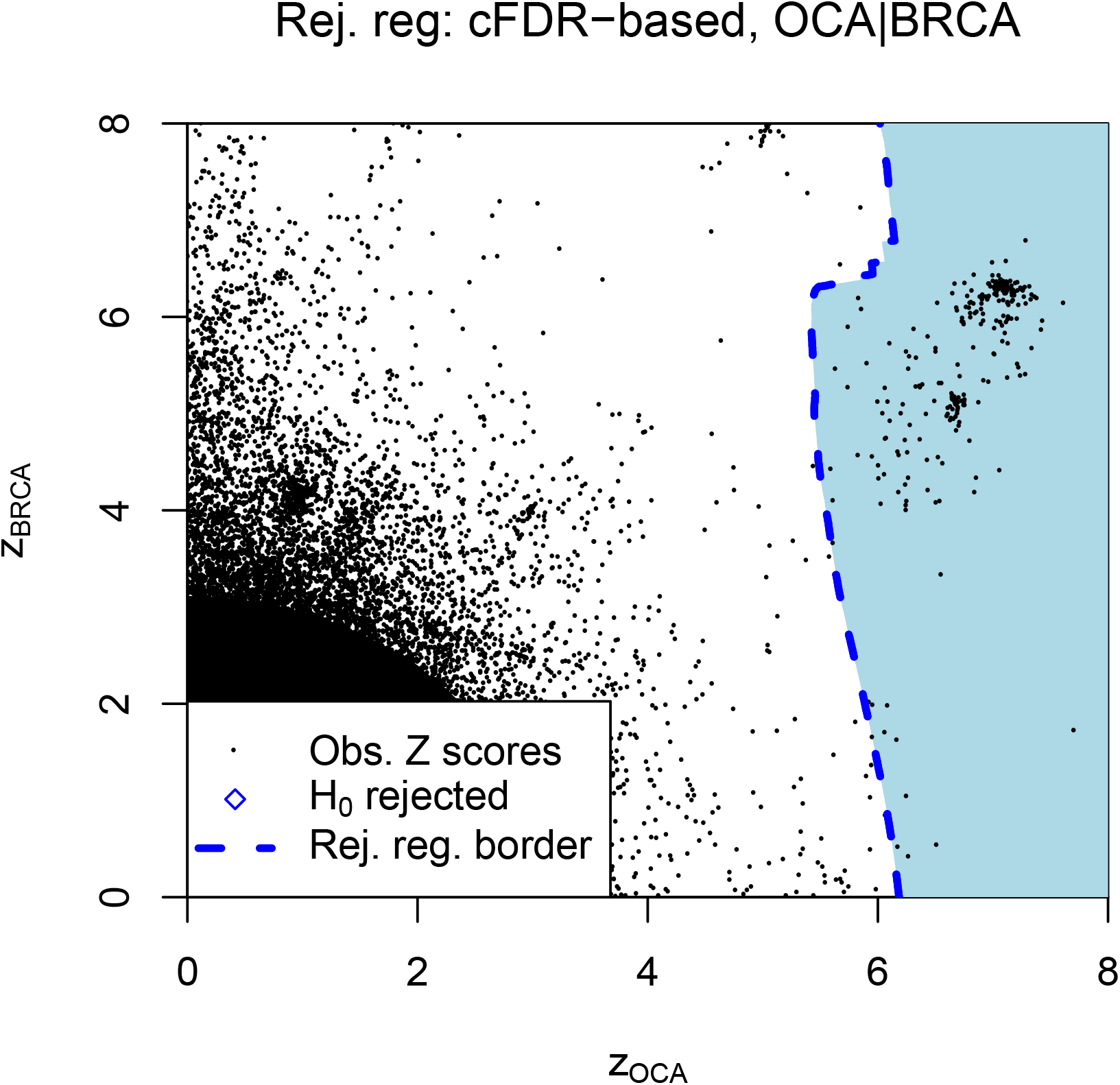
Association analysis of OCA using test statistics for BRCA as covariates. Variables and methods are similar to panel C in figure 1 in the main paper.

**Figure 19:**
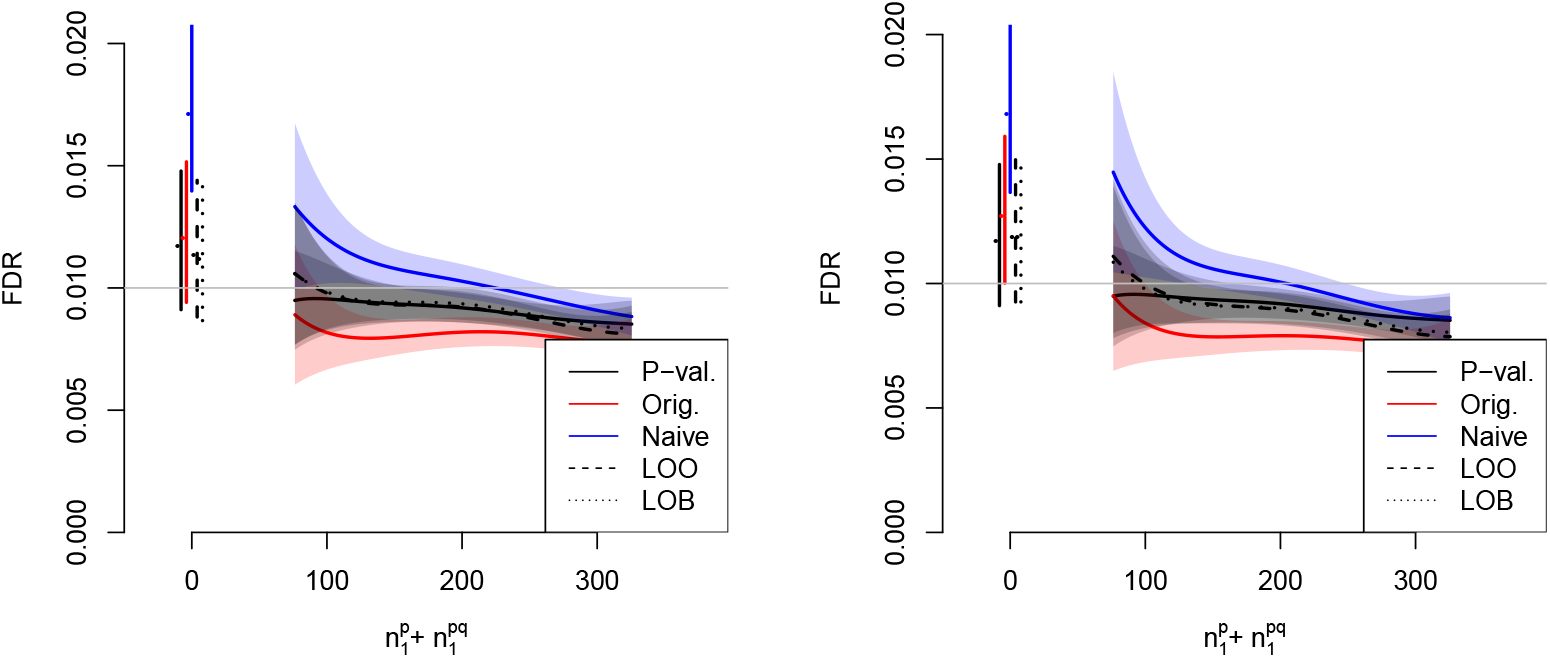
FDR control of various methods against 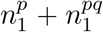, the total number of variables associated with *P* (the primary study under consideration). The horizontal line shows *α* = 0.01, the desired FDR control level. Simulations in the left panel integrate L-regions over the the true distribution *f*_0_; simulations in the right panel integrate over the estimated distribution as per equation (33). Shaded regions indicate 95% confidence envelopes. Curves show moving weighted averages using a Gaussian kernel with SD 15% of the X axis range. Lines on the left indicate FDR control with 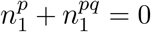.

**Figure 20:**
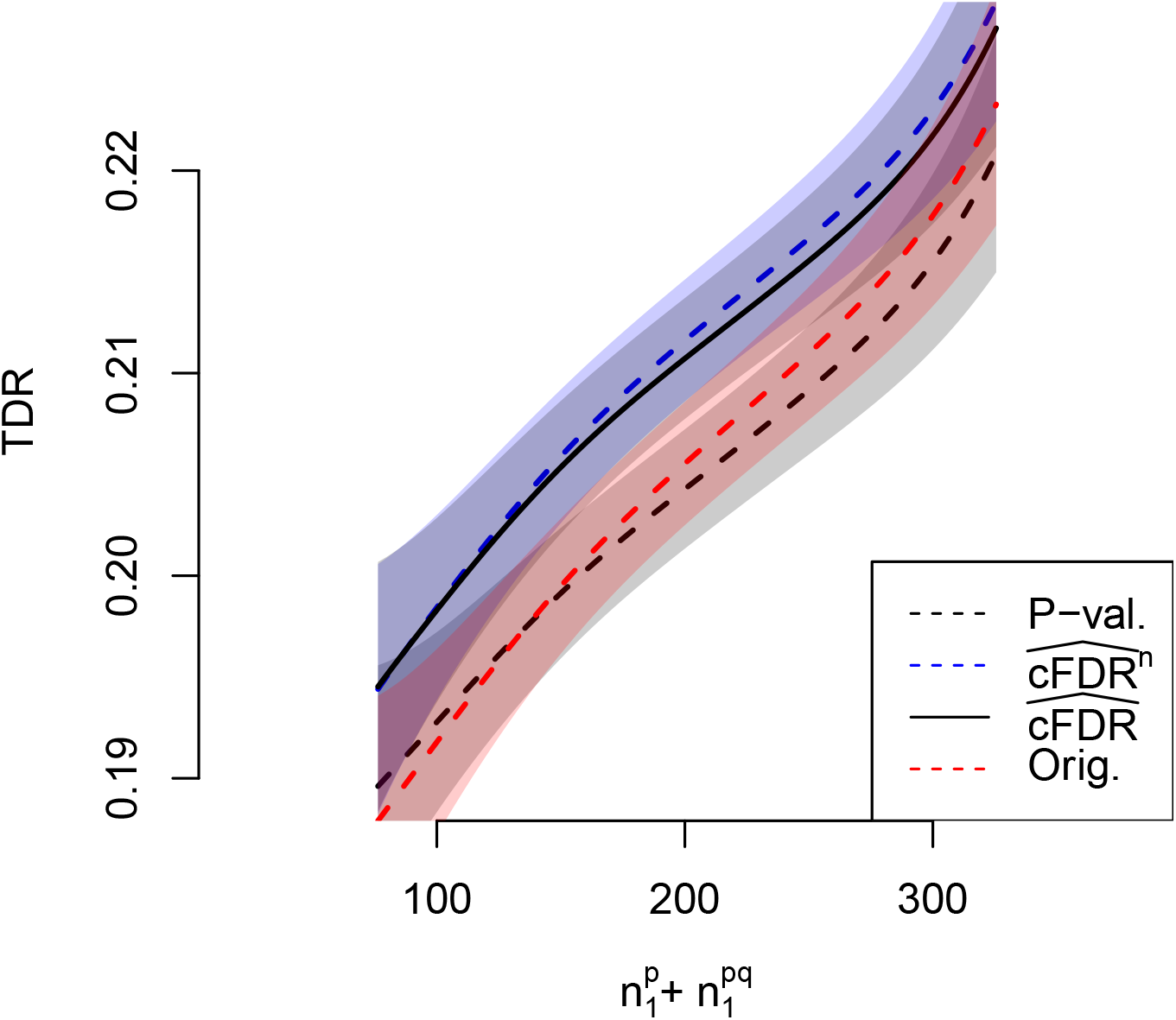
TDR of various methods against 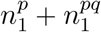, the total number of variables associated with *P* (the primary study under consideration), at FDR control level *α* = 0.01. Shaded areas show 95% pointwise confidence envelopes. Curves show moving weighted averages using a Gaussian kernel with SD 15% of the X axis range.

**Figure 21:**
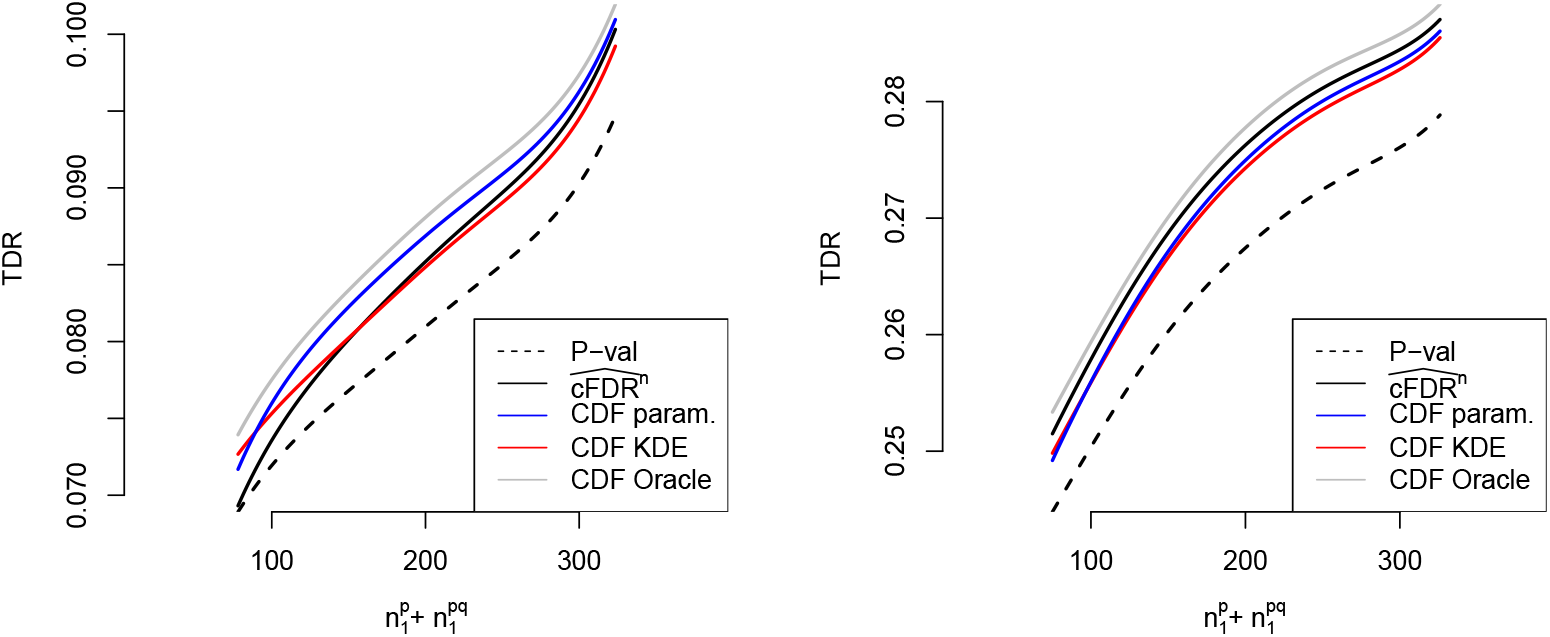
TDR of various methods against 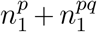, the total number of variables associated with *P* (the primary study under consideration), restricting to simulations in which parametric assumptions were satisfied (left panel) or were not satisfied (right panel), at FDR control level *α* = 0.01. Curves show moving weighted averages using a Gaussian kernel with SD 3/10 of the X axis range.

**Figure 22:**
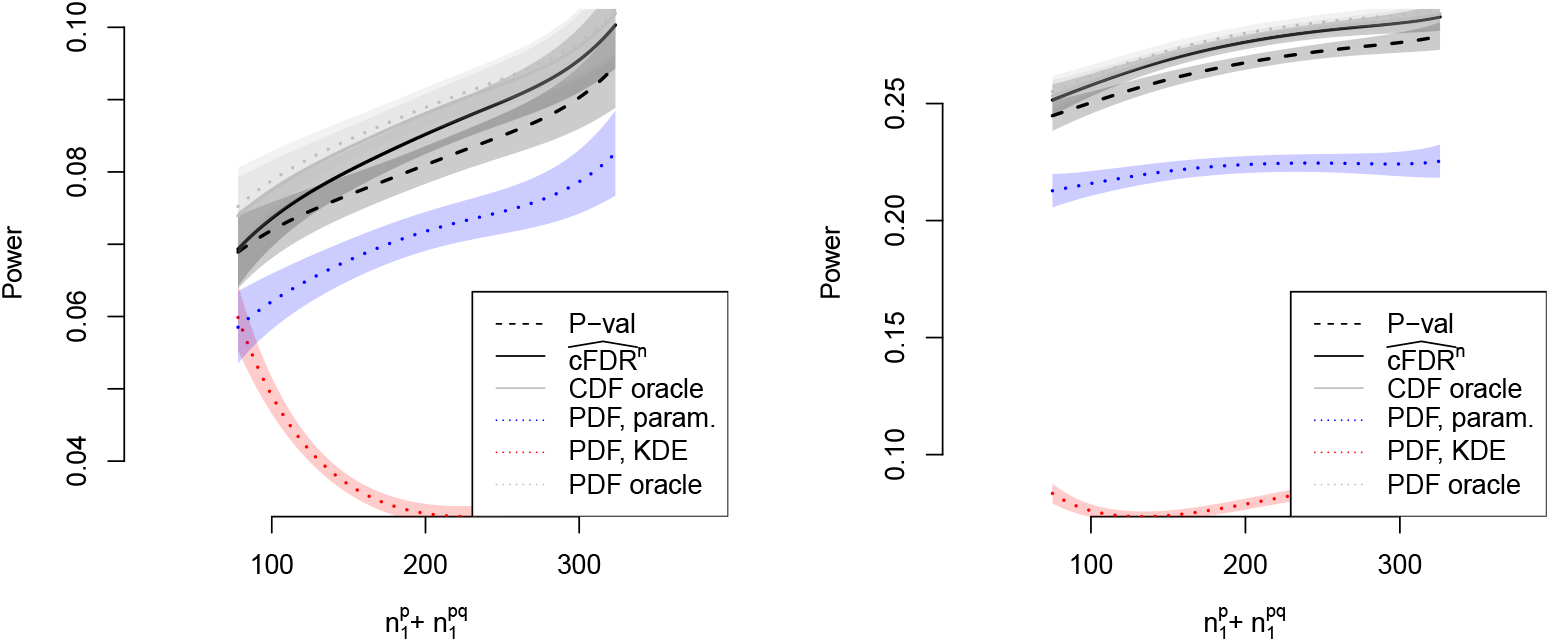
TDR of PDF-based methods against 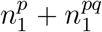, the total number of variables associated with *P* (the primary study under consideration), controlling FDR at *α* = 0.01. In the left panel, parametric assumptions were satisfied (ie *d* = 1 in table 1) and in the right panel, they are not (*d* = 2, 3). Shaded regions show pointwise 95% confidence intervals. Curves show moving weighted averages using a Gaussian kernel with SD 15% of the X axis range.

**Figure 23:**
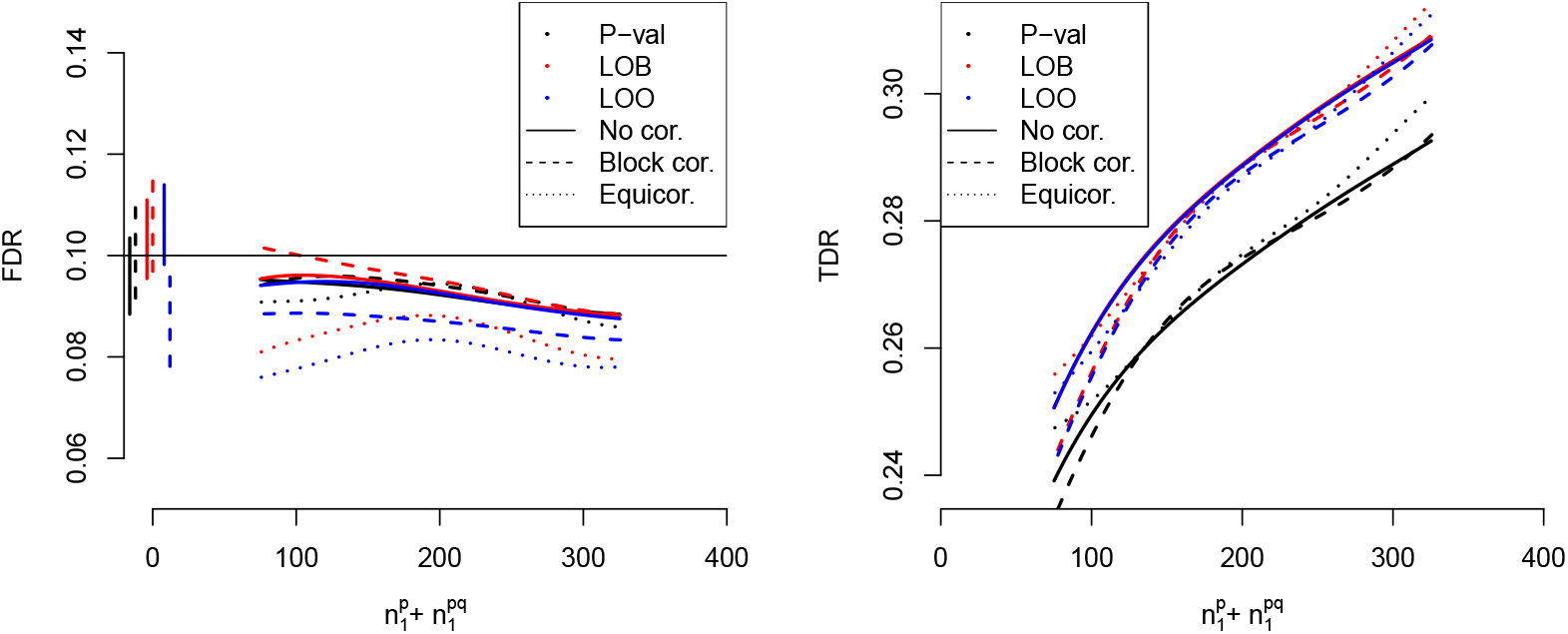
FDR (left) and TDR (right) of FDR-controlling methods leave-out-block (equation 26) and leave-one-out (equation 25) applied to 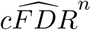, and the BH procedure applied to p-values, under several models of correlation between observations (*ρ* = 0.1). Confidence envelopes are omitted for visual clarity. Vertical lines show FDR with 95% confidence intervals at 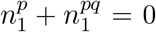. Curves show moving weighted averages using a Gaussian kernel with SD 15% of the X axis range.

**Figure 24:**
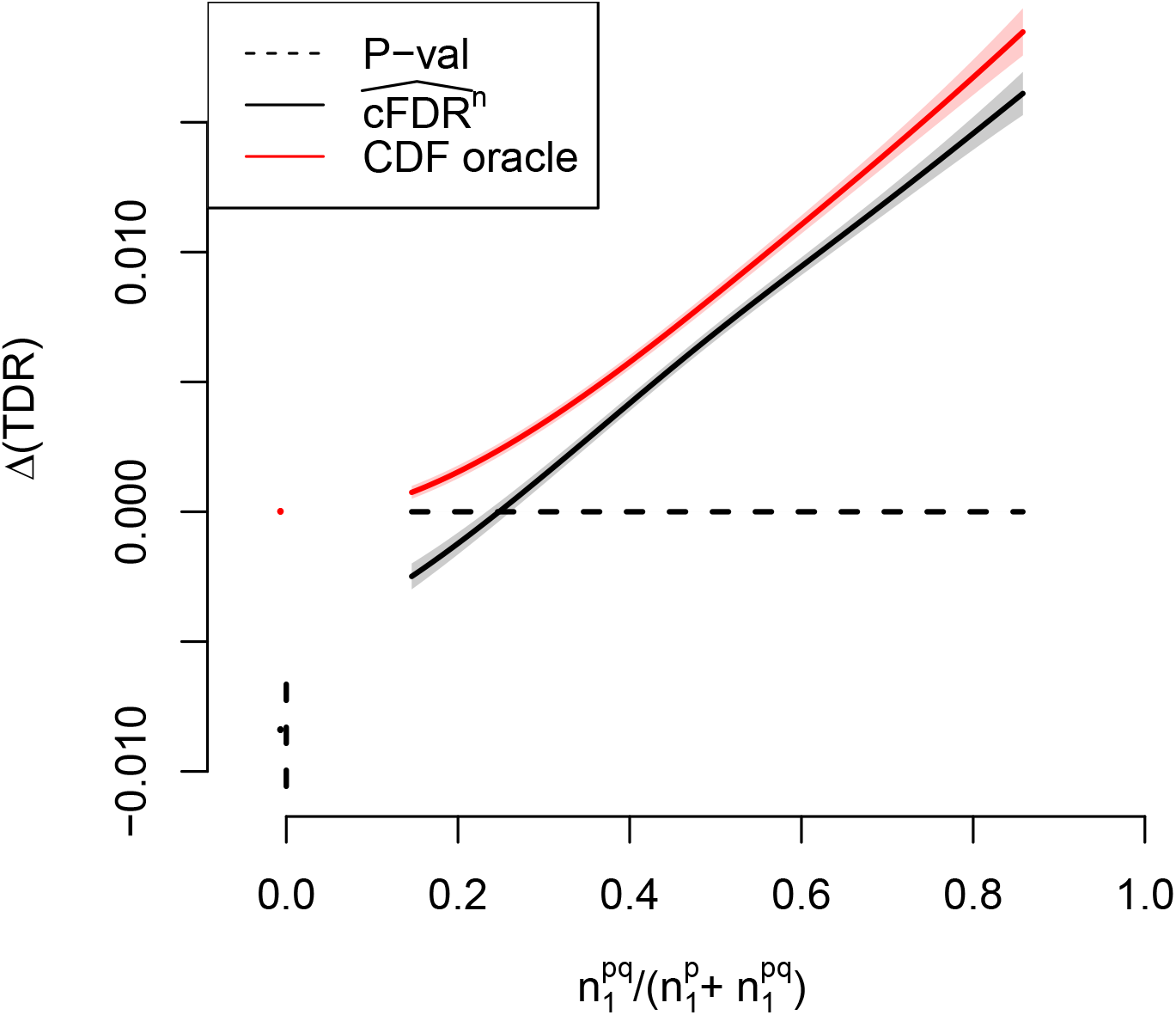
Difference in TDR between 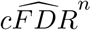 (assessed by leave-one-out v-values) and p-values, controlling FDR at *α* = 0.01, against 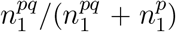 (proportion of non-null hypotheses for *P* which are shared with *Q*). The performance of the oracle CDF method is shown for comparison. Shaded areas show pointwise 95% confidence intervals. Points and lines at the leftmost edge show TDR values and 95% confidence intervals when 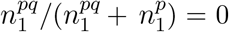. Curves show moving weighted averages using a Gaussian kernel with SD 15% of the X axis range.

## 11 Supplementary tables

**Table 3:**
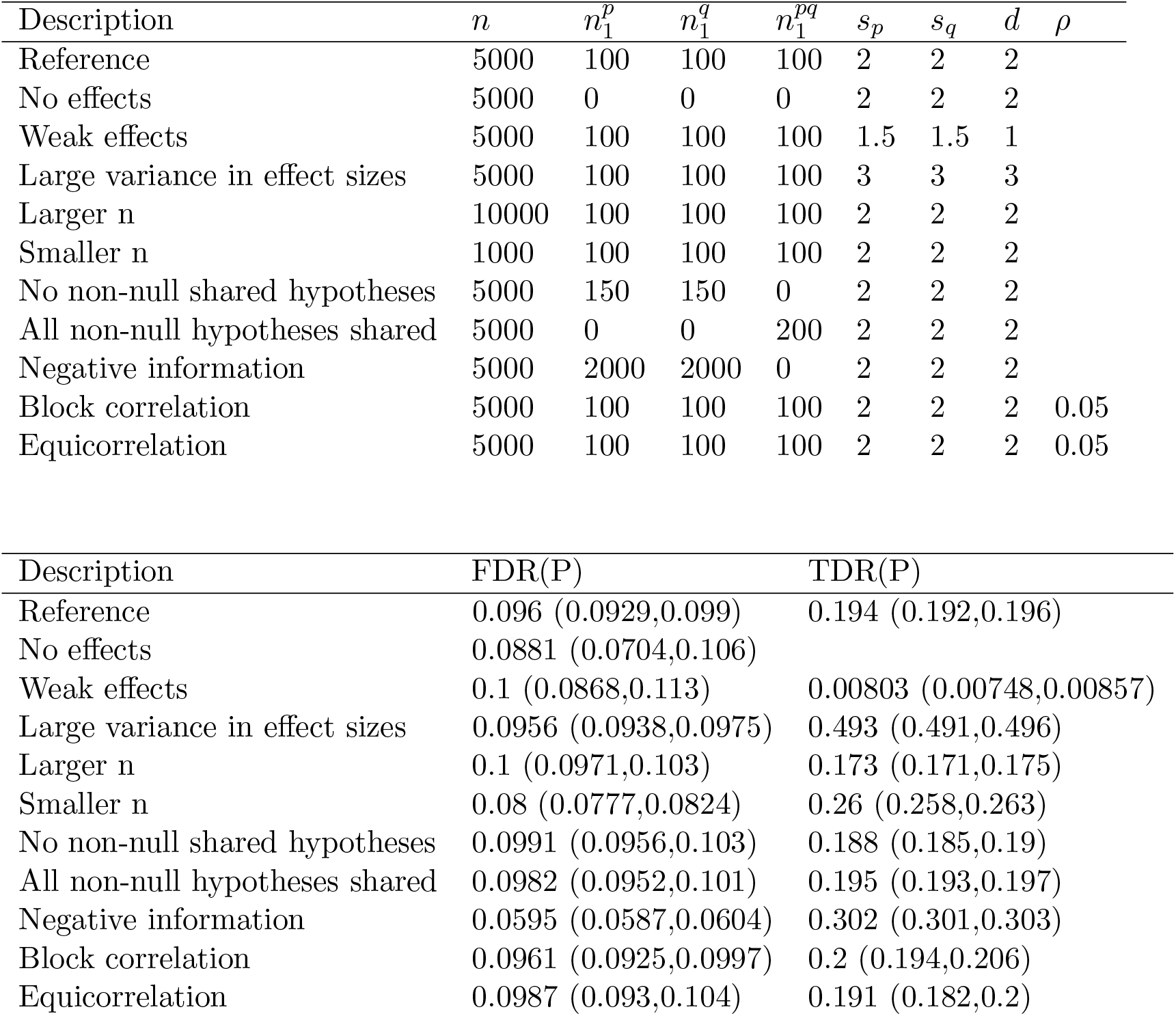
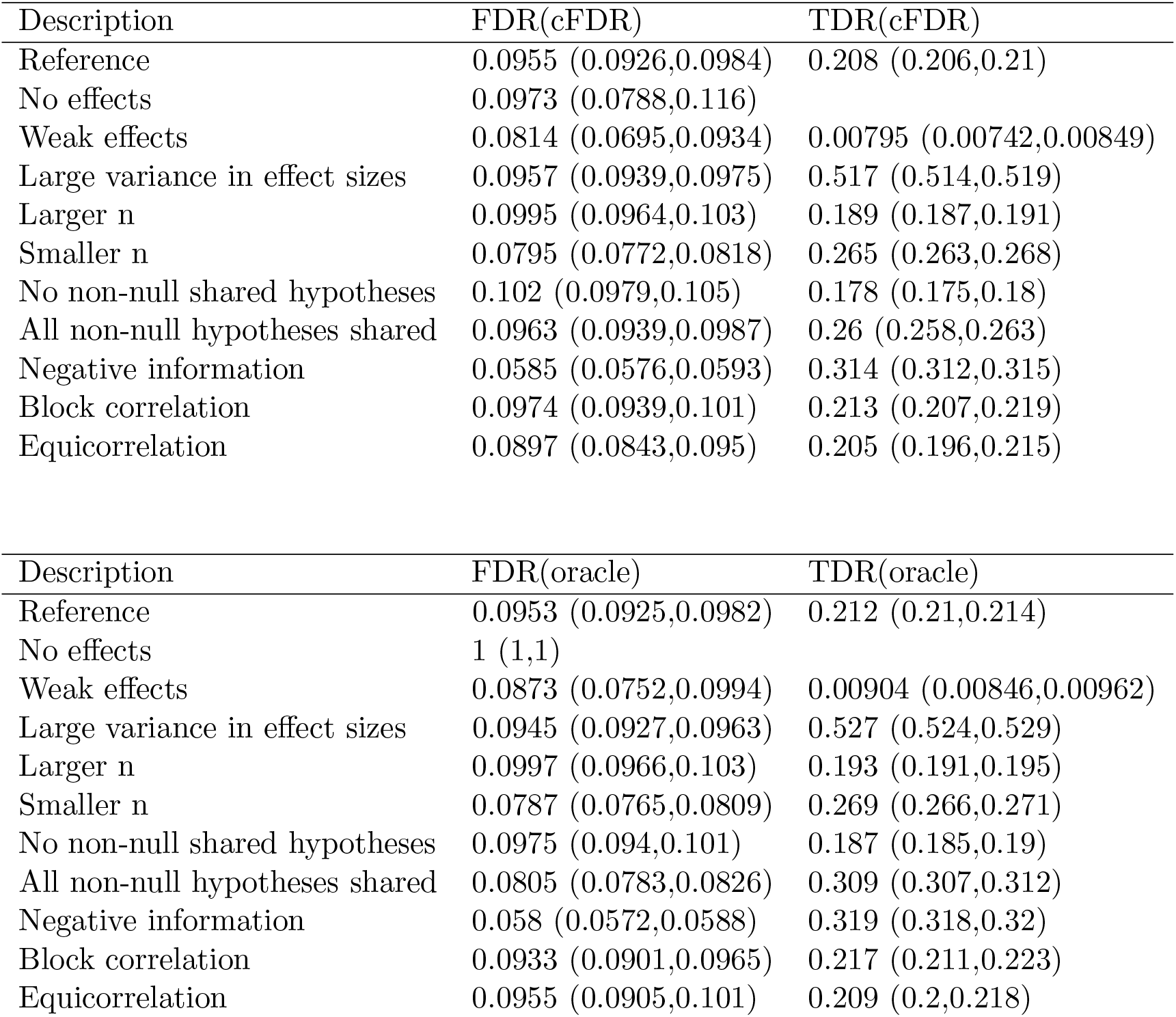
FDR and TDR of p-value, 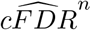, and oracle cfdr (best possible procedure) using leave-one-out v-values (equation 25) for a range of simulation parameters, controlling FDR at *α* = 0.1. Cells show mean and 95% confidence interval. TDR is undefined if 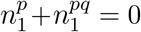.

| Description | $n$ | $n_1^p$ | $n_1^q$ | $n_1^{pq}$ | $s_p$ | $s_q$ | $d$ | $\rho$ |
| --- | --- | --- | --- | --- | --- | --- | --- | --- |
| Reference | 5000 | 100 | 100 | 100 | 2 | 2 | 2 |  |
| No effects | 5000 | 0 | 0 | 0 | 2 | 2 | 2 |  |
| Weak effects | 5000 | 100 | 100 | 100 | 1.5 | 1.5 | 1 |  |
| Large variance in effect sizes | 5000 | 100 | 100 | 100 | 3 | 3 | 3 |  |
| Larger n | 10000 | 100 | 100 | 100 | 2 | 2 | 2 |  |
| Smaller n | 1000 | 100 | 100 | 100 | 2 | 2 | 2 |  |
| No non-null shared hypotheses | 5000 | 150 | 150 | 0 | 2 | 2 | 2 |  |
| All non-null hypotheses shared | 5000 | 0 | 0 | 200 | 2 | 2 | 2 |  |
| Negative information | 5000 | 2000 | 2000 | 0 | 2 | 2 | 2 |  |
| Block correlation | 5000 | 100 | 100 | 100 | 2 | 2 | 2 | 0.05 |
| Equicorrelation | 5000 | 100 | 100 | 100 | 2 | 2 | 2 | 0.05 |

| Description | FDR(P) | TDR(P) |
| --- | --- | --- |
| Reference | 0.096 (0.0929,0.099) | 0.194 (0.192,0.196) |
| No effects | 0.0881 (0.0704,0.106) |  |
| Weak effects | 0.1 (0.0868,0.113) | 0.00803 (0.00748,0.00857) |
| Large variance in effect sizes | 0.0956 (0.0938,0.0975) | 0.493 (0.491,0.496) |
| Larger n | 0.1 (0.0971,0.103) | 0.173 (0.171,0.175) |
| Smaller n | 0.08 (0.0777,0.0824) | 0.26 (0.258,0.263) |
| No non-null shared hypotheses | 0.0991 (0.0956,0.103) | 0.188 (0.185,0.19) |
| All non-null hypotheses shared | 0.0982 (0.0952,0.101) | 0.195 (0.193,0.197) |
| Negative information | 0.0595 (0.0587,0.0604) | 0.302 (0.301,0.303) |
| Block correlation | 0.0961 (0.0925,0.0997) | 0.2 (0.194,0.206) |
| Equicorrelation | 0.0987 (0.093,0.104) | 0.191 (0.182,0.2) |

See continuation on following page
| Description | FDR(cFDR) | TDR(cFDR) |
| --- | --- | --- |
| Reference | 0.0955 (0.0926,0.0984) | 0.208 (0.206,0.21) |
| No effects | 0.0973 (0.0788,0.116) |  |
| Weak effects | 0.0814 (0.0695,0.0934) | 0.00795 (0.00742,0.00849) |
| Large variance in effect sizes | 0.0957 (0.0939,0.0975) | 0.517 (0.514,0.519) |
| Larger n | 0.0995 (0.0964,0.103) | 0.189 (0.187,0.191) |
| Smaller n | 0.0795 (0.0772,0.0818) | 0.265 (0.263,0.268) |
| No non-null shared hypotheses | 0.102 (0.0979,0.105) | 0.178 (0.175,0.18) |
| All non-null hypotheses shared | 0.0963 (0.0939,0.0987) | 0.26 (0.258,0.263) |
| Negative information | 0.0585 (0.0576,0.0593) | 0.314 (0.312,0.315) |
| Block correlation | 0.0974 (0.0939,0.101) | 0.213 (0.207,0.219) |
| Equicorrelation | 0.0897 (0.0843,0.095) | 0.205 (0.196,0.215) |

Table 3: FDR and TDR of p-value, $c\widehat{FDR}^n$ , and oracle cfdr (best possible procedure) using leave-one-out v-values (equation 25) for a range of simulation parameters, controlling FDR at $\alpha = 0.1$ .
| Description | FDR(oracle) | TDR(oracle) |
| --- | --- | --- |
| Reference | 0.0953 (0.0925,0.0982) | 0.212 (0.21,0.214) |
| No effects | 1 (1,1) |  |
| Weak effects | 0.0873 (0.0752,0.0994) | 0.00904 (0.00846,0.00962) |
| Large variance in effect sizes | 0.0945 (0.0927,0.0963) | 0.527 (0.524,0.529) |
| Larger n | 0.0997 (0.0966,0.103) | 0.193 (0.191,0.195) |
| Smaller n | 0.0787 (0.0765,0.0809) | 0.269 (0.266,0.271) |
| No non-null shared hypotheses | 0.0975 (0.094,0.101) | 0.187 (0.185,0.19) |
| All non-null hypotheses shared | 0.0805 (0.0783,0.0826) | 0.309 (0.307,0.312) |
| Negative information | 0.058 (0.0572,0.0588) | 0.319 (0.318,0.32) |
| Block correlation | 0.0933 (0.0901,0.0965) | 0.217 (0.211,0.223) |
| Equicorrelation | 0.0955 (0.0905,0.101) | 0.209 (0.2,0.218) |

